# Learning cell-specific networks from dynamics and geometry of single cells

**DOI:** 10.1101/2023.01.08.523176

**Authors:** Stephen Y Zhang, Michael P H Stumpf

**Author notes:** stephenz <at> student.unimelb.edu.au (Stephen Y Zhang).

## Abstract

Cell dynamics and biological function are governed by intricate networks of molecular interactions. Inferring these interactions from data is a notoriously difficult inverse problem. The majority of existing network inference methods work at the population level to construct population-averaged representations of gene interaction networks, and thus do not naturally allow us to infer differences in gene regulation activity across heterogeneous cell populations. We introduce locaTE, an information theoretic approach that leverages single cell dynamical information together with geometry of the cell state manifold to infer cell-specific, causal gene interaction networks in a manner that is agnostic to the topology of the underlying biological trajectory. We find that factor analysis can give detailed insights into the inferred cell-specific GRNs. Through extensive simulation studies and applications to three experimental datasets spanning mouse primitive endoderm formation, pancreatic development, and haematopoiesis, we demonstrate superior performance and the generation of additional insights compared to standard static GRN inference methods. We find that locaTE provides a powerful, efficient and scalable network inference method that allows us to distill cell-specific networks from single cell data.

**Graphical abstract:** **Cell-specific network inference from estimated dynamics and geometry**

LocaTE takes as input a transition matrix *P* that encodes inferred cellular dynamics as a Markov chain on the cell state manifold. By considering the coupling (*X*_*τ*_, *X*_−*τ*_), locaTE produces an estimate of transfer entropy for each cell *i* and each pair of genes (*j, k*). Downstream factor analyses can extract coherent patterns of interactions in an unsupervised fashion.

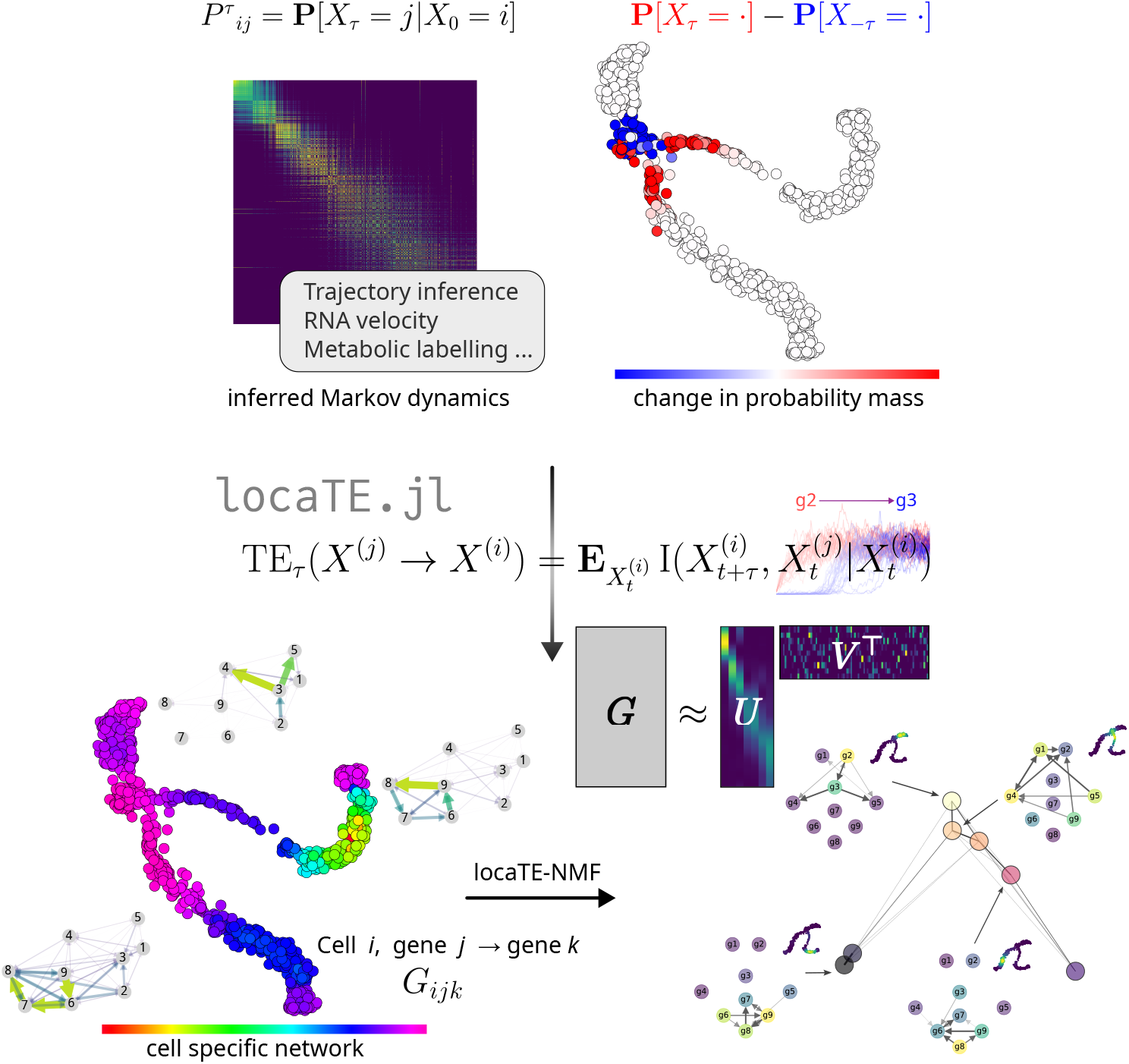

## 1 Introduction

Cell identity is determined by the dynamics underlying gene expression. Gene expression, in turn, is controlled by gene regulatory networks that direct the activity of genes over time and set the differences between cells. Identifying and characterising gene regulatory interactions has become a central aim of cell and systems biology. Single cell assays have been enabling us to capture population heterogeneity in gene expression, allowing for the detection and study of rare and transient cellular phenotypes that would be unobservable from bulk studies [1]. This has demonstrated variability within groups of cells which would conventionally have been regarded as being of the same “type”.

Modern single cell datasets are of a size that has allowed us to overcome statistical problems that have plagued network inference from bulk data [2, 3] In this vein, a collection of methods have been developed for network inference from single cell data, some of which build on methods originally intended for bulk expression data, and others which are specifically tailored to single cell data. The recent surveys in [4–6] provide an overview of the available methodology. The overwhelming majority of network inference methods aim to reconstruct a single *static* network [4,6] that describes the set of possible interactions occurring within an observed population of cells. This is a missed opportunity: cellular heterogeneity at the transcriptional level is ubiquitous and functionally important, and we might expect that variations in (cell state dependent) regulatory interactions play a part in generating and propagating this heterogeneity. Evidence for cell and cell-type specific networks has been observed empirically in previous studies [7, 8], but static networks are not able to capture such dynamics [3].

The need for methods that can learn cell- or phenotype-specific networks [3, 9, 10], is now leading to new classes of network inference methods [11–14]. These methods seek to infer multiple networks (e.g. one per cell type, or even one per cell) from single-cell resolved data, thus allowing us to learn about variation in gene interactions over time and between conditions. Such methods generally rely on neighbourhood or cluster information to construct networks that are cell, time, or context-specific. Existing methods for cell-specific network inference have focused on inferring undirected networks [11–13]. If we hope to infer *causal* or *directed* interactions, then information about underlying dynamics in a cell population is essential [15, 16].

In order to infer cell specific networks we need to learn how far along a (e.g. developmental) trajectory a cell has travelled; which cells represent earlier stages; and which cells represent later stages. Pseudotemporal ordering is one approach, and many methods for static network inference take advantage of such an ordering [15, 17–20]. However, the one-dimensional nature of pseudotime necessarily imposes a total ordering over cells, resulting in a loss of finer structure of the cell state manifold. Recent developments in trajectory inference from a Markov process viewpoint [21–24] depart from this framework and are able to model more complex dynamics on the set of observed cell states directly, and are free from assumptions on the trajectory topology. Manifold learning approaches [25,26] construct a cell-state graph by learning local neighbourhoods. They avoid clustering of cell states and can model arbitrarily complex trajectory structures (see Box 1 for further discussion).

In this article, we place transfer entropy (TE) [27] as an information-theoretic measure of causality into the context of a manifold in gene expression space. Transfer entropy, like other information-theoretical measures [28], is a model-free framework for measuring dependencies between random variables. It has been used widely in contexts such as neuroscience where abundant time-series data are available. In the Gaussian case it is equivalent to Granger causality [29]. Transfer entropy has been used to predict static networks using a pseudotemporal ordering [15, 17]. Here we demonstrate that the transfer entropy can be adapted to infer cell-specific networks directly from dynamics on the cell-state manifold. Crucially we do not have to impose an ordering of cells. The ability to infer cell-specific will, for example, allow us to tackle problems in developmental biology. Here it is often the transient dynamics that are the most interesting [30, 31] since these are likely to correspond to major epigenetic changes [3]. Our approach is tailored to such problems and we make no restrictive simplifying assumptions: the cell state (gene expression) space is treated as a continuum; we do not require *a priori* partitioning or clustering to group or classify cells.

Having introduced locaTE as a dynamics and geometry-driven network inference framework, we first illustrate its accuracy and utility in two in-silico examples. In these systems the underlying regulatory interactions are dynamically rewired, making the ability to infer context-specific interactions crucial to achieving good performance.

### Box 1: Beyond pseudotime: geometry of the cell-state manifold

Developmental systems are by nature dynamic. C. H. Waddington’s classical landscape metaphor [32] conceptualised biological processes as cells traversing down a potential landscape from an uphill progenitor state to mature cell types. In this landscape, attractors corresponding to biological cell types are separated by saddle nodes, which can be thought of as transitory states at which “fate decisions” are made by differentiating cells. Single cell profiling makes it possible to measure not only cells within attractors, but also transient cell states at these decision points [30, 33]. These intermediate states were invisible to bulk analyses [34].

Identification of transient cell states requires knowledge of the (temporal) structure of the underlying biological process. The problem of recovering this information from single cell measurements is called trajectory inference [35]. For example, pseudotime imposes an ordering of cells based on relative progression through the developmental process. Pseudotime is fundamentally limited to a small set of trajectory topologies that it can handle: a total ordering of cells can only accurately reflect a linear topology. Branched pseudotime methods produce a partial ordering of cells which can describe tree-like topologies. In two or more dimensions, trajectories can exhibit cycles; in four or more dimensions, more complex topologies are theoretically possible (the Klein bottle provides an example of one such “esoteric” structure). Trajectories that depart from linear and tree-like topologies cannot be accurately represented by pseudotime methods. Approaches based on manifold learning, by contrast, aim to construct local neighbourhoods of cells. They can handle arbitrary topologies and are only limited by the amount of available data from which to learn the manifold. Since single cell datasets are naturally embedded in ∼ 10^4^ (the number of discernible transcripts) dimensions, there are a vast number of degrees of freedom and hence ways in which linear or tree-like topological assumptions may be violated.

### Box 2: Single cell dynamics and transfer entropy

Trajectory inference methods aim to recover dynamics from single cell data, but have limitations on the dynamics they can model. From geometry of single cell measurements alone, construction of a random walk over observed cell states is consistent with an inferred drift of **v =** ∇ log *ρ* where *ρ* is the density of observed cells [34, 36] – if the observation process draws from a closed system at equilibrium then this recovers the potential Ψ. Methods such as pseudotime, optimal transport and population balance analysis implicitly assume potential-driven dynamics **v = −**∇Ψ and thus cannot model behaviours such as cycles [21]. Other approaches, such as RNA velocity, metabolic labelling and live imaging, enable direct quantification of the vector field [24, 37]. These methods have the potential to identify dynamics that are not captured by geometry or trajectory inference, for example the periodic dynamics of the cell cycle. We discuss this further in 4.1.

The mutual information between two random variables is a non-parametric measure of statistical dependence that is free of distributional assumptions and can capture non-linear relationships between variables. For two random variables, *X, Y*, with joint distribution, *p*_*XY*_, the mutual information is defined as

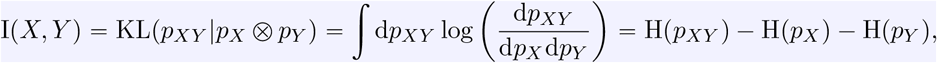

where H(*p*) is Shannon’s (neg)entropy, 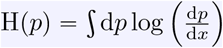. We have *I*(*X, Y*) ⩾ 0, and *I*(*X, Y*) = 0 if and only if *X* and *Y* are independent, i.e. **P (***X, Y) =* **P (***X*) **P (***Y*). For a third random variable *Z*, the *conditional* mutual information can be defined in terms of a conditional expectation

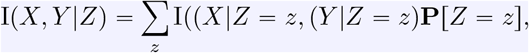

where *X* |*Z = z, Y* |*Z = z* denote random variables conditioned on the event {*Z= z*}.

The *transfer entropy* (TE) between two time-dependent random variables (*X*_*t*_, *Y*_*t*_) measures information transfer from *X*_*t*_ to *Y*_*t*_ over time. For a chosen time lag parameter, *τ >*0, the transfer entropy TE [*X*_*t*_ *→Y*_*t*_ ]is defined as I (*X*_*t*_, *Y*_*t τ*_ |*Y*_*t*_), the mutual information between *X*_*t*_ and *Y*_*t+ τ*_ conditioned on *Y*_*t*_ in order to remove the dependence of *Y* on itself. We refer the reader to Section 4 for a extended discussion of these topics.

## 2 Results

### 2.1 Overview of locaTE

LocaTE models biological dynamics as Markov processes {*X*_*t*_, *t* ⩾0} supported on a cell state manifold ℳ ⊆ 𝒳, a subset of the space 𝒳 of single-cell gene expression profiles (see Box 1). Specifically, locaTE is designed for single cell snapshot experiments, where cells are sampled asynchronously from ℳ across a spectrum of developmental stages [21]. We assume that dynamics are cell-autonomous and well-described by Markov transition kernels {*P*_*τ*_ *(x, x*^′^) : *τ* ⩾0 supported on ℳ × ℳ. In practice, the cell state manifold ℳ can be approximated by a number of manifold learning approaches [25]. Estimates of the transition kernel *P*_*τ*_ can similarly be obtained from experimental and computational approaches. These include but are not limited to: pseudotime estimation [38], population dynamics inference [21, 39], RNA velocity [37, 40, 41], and metabolic labelling [24, 42]. In scenarios where dynamics cannot be readily inferred, we propose a simple construction for *P*_*τ*_ which has a theoretical motivation and yields good results in practice. We provide an in-depth discussion of dynamical inference approaches and their relative advantages in Section 4.

In practice, for a set of measured single cell profiles 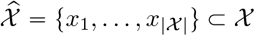, we construct an approximation 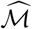 of the cell state manifold ℳ, together with a transition matrix 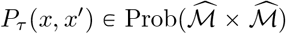. Taken together, these encode stochastic dynamics which are constrained to lie on a manifold of cell states. Given measurements of interest (e.g. gene expression) for each cell 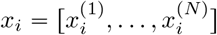, locaTE infers cell-specific networks that capture directed information flows between variables in the system. To achieve this, locaTE uses the dynamics information encoded in *P*_*τ*_ and a generalisation of the transfer entropy (see Box 2). Under the Markov model of cell dynamics, these allow us to measure the localised flow of information between variables in this high-dimensional stochastic dynamical system.

Raw cell-specific TE estimates are noisy, and the signal may be degraded by transcriptional noise and technical dropouts. Motivated by classical graph signal processing techniques, we propose to distil meaningful interaction signals from noisy TE estimates by utilising the manifold hypothesis. Encoding geometric information of the empirical cell state manifold 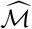 in terms of the graph Laplacian, cell-specific networks are denoised using a Laplacian-regularised regression. Intuitively, this is motivated by the expectation that cells that are proximal 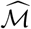 in should have similar interaction networks. Following the sparsity-of-effect principle [43] we impose sparsity of the learned networks via a Lasso regularisation. Taken together, locaTE produces for each cell *x*_*i*_ a corresponding sparse matrix *A*_*i*_ encoding local interactions. Viewed differently, this associates each point 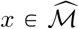 with an interaction network *A (x*). We refer to Section 5 for a in-depth discussion of locaTE, its assumptions and implementation.

### 2.2 LocaTE uncovers state-dependent logic in synthetic trajectories

We first consider two simulated trajectories where the true regulatory interactions are known by design. To illustrate the advantages of cell-specific networks, we consider examples where the active network is rewired in a cell-state dependent fashion. We consider a bifurcating system where regulatory interactions differ between branches, and a switch-like system where regulatory interactions vary continuously along a linear trajectory. Conventional approaches that infer population-level networks are insufficient to fully describe these systems. We show in Figure 3(a) the bifurcating network, in which a toggle-switch feeds into one of two gene expression modules, denoted A and B. Each of these modules involve the same genes {*g*6, *g*7, *g*8, *g*9}, but in each module the flow of information in each module is the mirror image of the other. Alternatively, the “context-dependence” of interactions in this system can be understood as arising from the presence of higher-order interactions. For instance, gene 7 may either be activated by gene 6 (in module A) or by gene 8 (in module B). This can be understood a *hyperedge* with a Boolean rule

**Figure 1:**
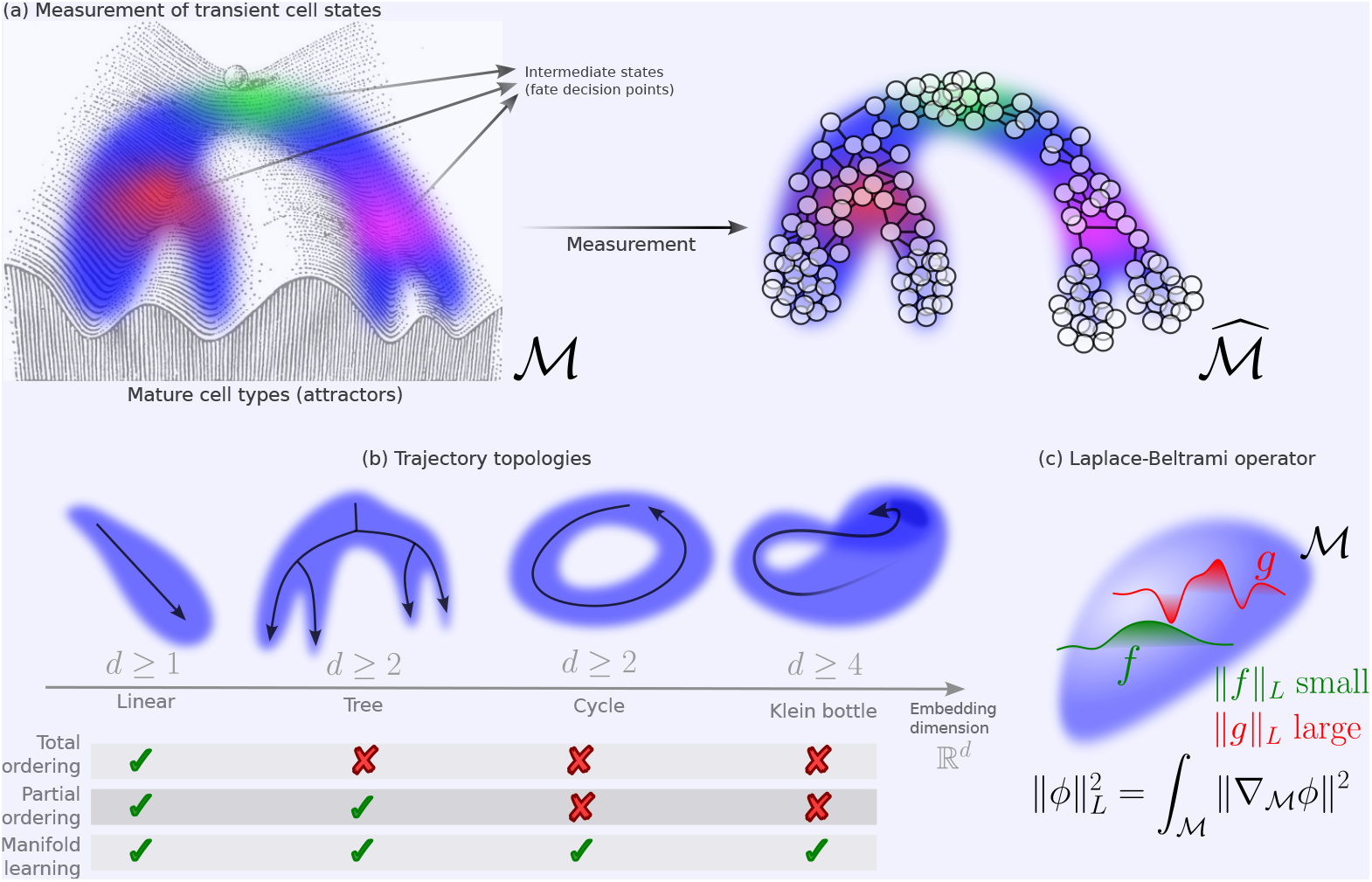
(a) Manifold ℳ of cell states overlaid on the classical Waddington’s landscape: intermediate cell states correspond to points of fate decision. Measurement by single-cell technologies produces a set of observed cell states, from which an approximation, 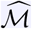, of the underlying cell state manifold can be learned. (b) Total and partial orderings (and thus, pseudotemporal orderings and extensions thereof) are limited to linear and tree-like trajectory topologies, while manifold learning approaches, which rely on construction of *neighbourhoods* of points, are by nature topology-agnostic. (c) The Laplace-Beltrami operator is a central object describing a manifold ℳ and in particular compares functions in a manner adapted to the underlying geometry.

**Figure 2:**
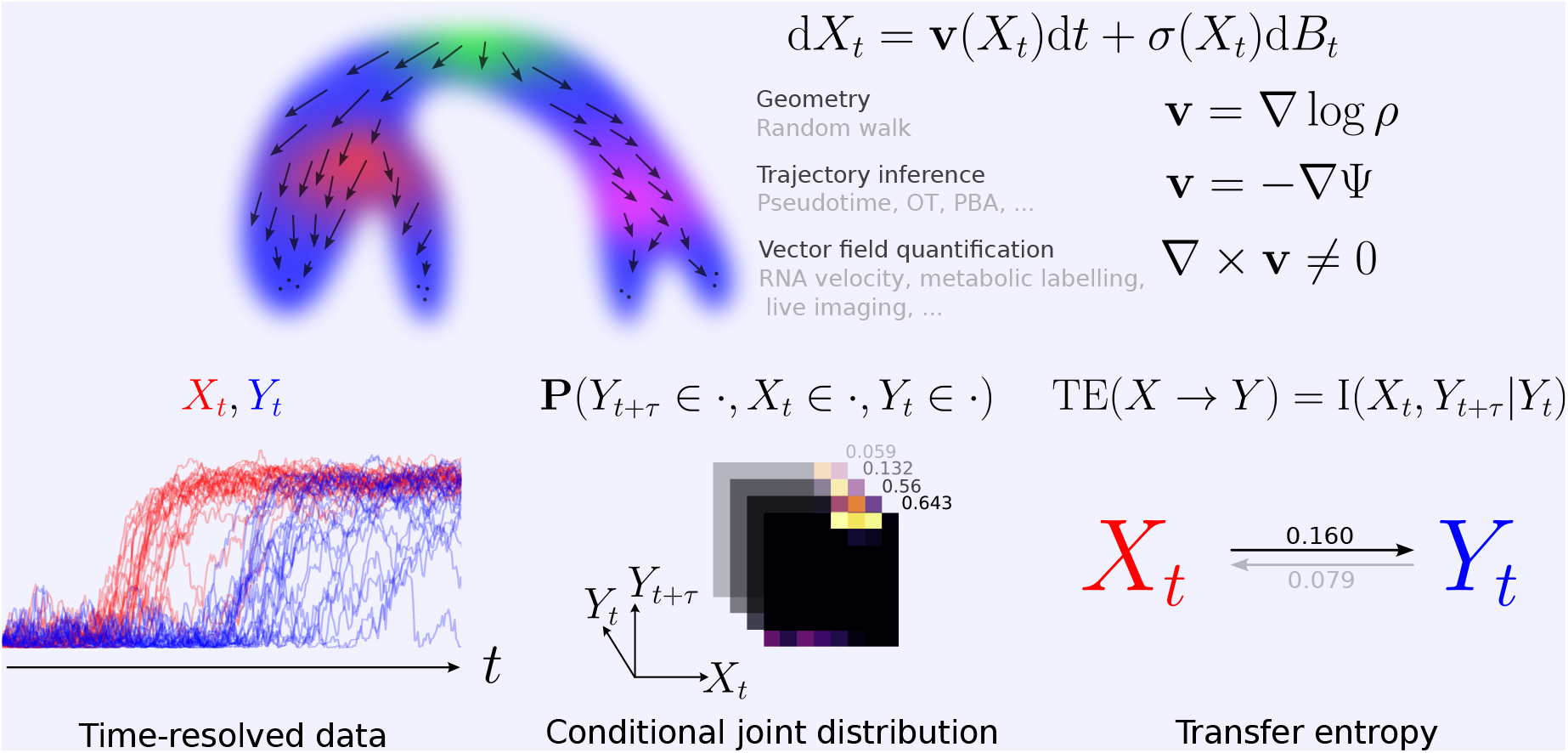
Top: Single cell dynamics reflect the underlying developmental program. Dynamics can be inferred using computational and experimental techniques, which may incur various structural limitations on the inference. Bottom: Transfer entropy is an information theoretic approach for quantifying information flow through a stochastic dynamical system.

**Figure 3:**
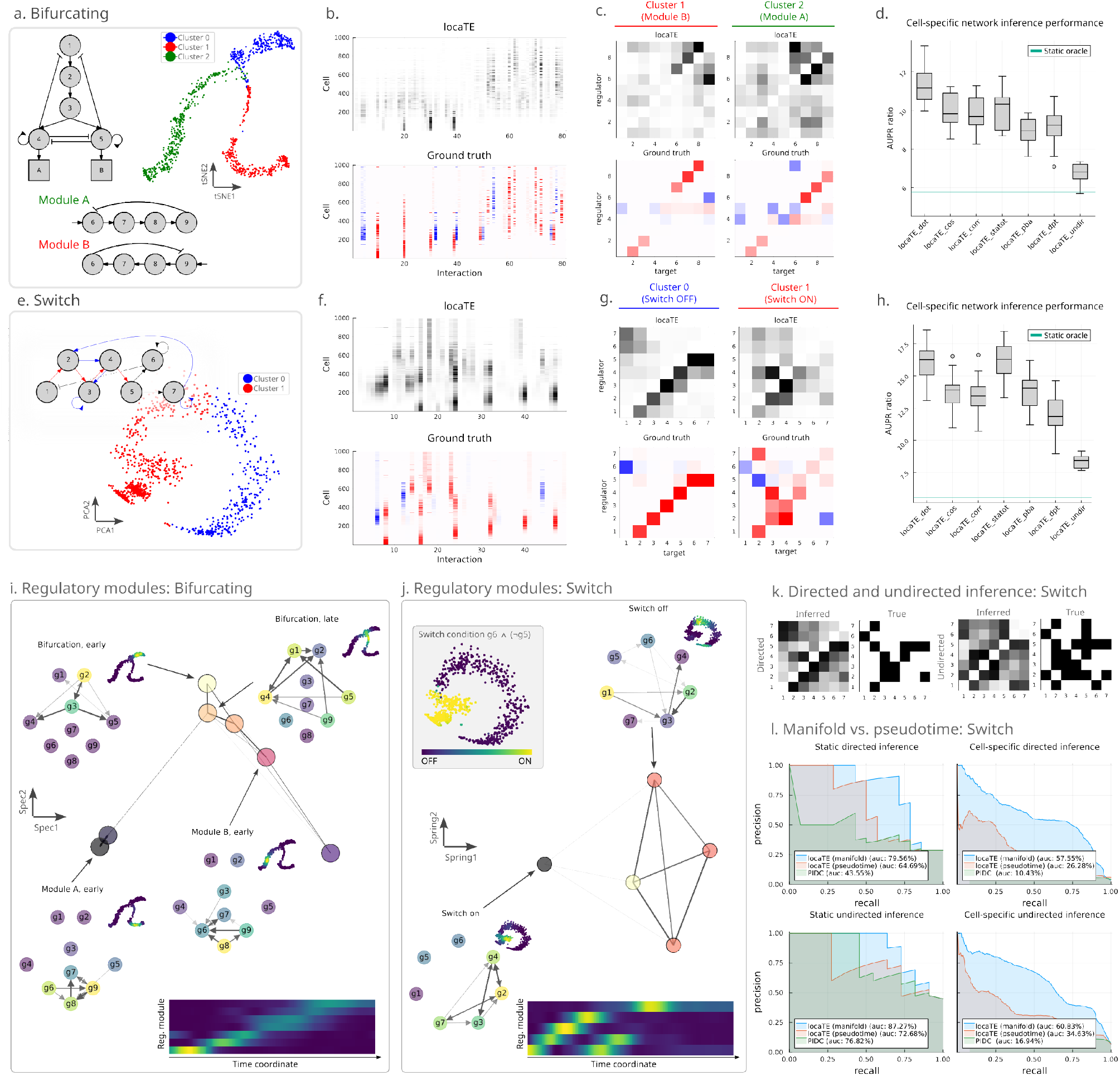
Simulated networks. **(a)** Bifurcating network: 5-gene bifurcating subnetwork feeding into two modules A, B involving the same 4 genes *g*6, *g*7, *g*8, *g*9. 4-gene modules A, B each involve different regulatory interactions involving the same species; tSNE embedding of simulated cell profiles, coloured by inferred diffusion pseudotime cluster. **(b)** LocaTE output as a *N* × *d*^2^ matrix in order of increasing pseudotime (top), shown alongside the ground truth Jacobians (bottom). **(c)** Interactions inferred by locaTE averaged by cluster (top) shown against ground truth Jacobians averaged by cluster. **(d)** AUPRC ratio for locaTE and other methods across 10 sampled simulation datasets, in the setting of cell-specific inference. We show as a horizontal line the AUPRC ratio achieved by perfect knowledge of the static network (static oracle). **(e)** Switch network: Blue (resp. red) interactions are active when (*g*6 ^ (⌐*g*5)) is FALSE (resp. TRUE). Notably, gene 2 interacts both directly and indirectly with gene 4, depending on the system state. **(f)** LocaTE output as a *N* ×*d*^2^ matrix in order of increasing pseudotime (top), shown alongside the ground truth Jacobians (bottom). **(g)** Interactions inferred by locaTE averaged by cluster (top) shown against ground truth Jacobians averaged by cluster. **(h)** AUPRC ratio for locaTE and other methods across 10 sampled simulation datasets, in the setting of cell-specific inference. We show as a horizontal line the AUPRC ratio achieved by perfect knowledge of the static network (static oracle). **(i)** Top: Regulatory modules for the simulated bifurcating network found by locaTE-NMF with *k* = 8, shown as a graph spectral layout. Selected modules are shown (top 10% of edges), along with the module activity on tSNE coordinates in inset. Bottom: smoothed module activities against simulated time. **(j)** Top: Regulatory modules for the simulated switch network found by locaTE-NMF with *k* = 5, shown as a graph spring layout. Selected modules are shown (top 20% of edges), along with the module activity on PCA coordinates in inset. Bottom: smoothed module activities against simulated time. **(k)** Comparison of population-averaged networks for switch system for inference of directed interactions utilising dynamical information (left) and inference of undirected interactions utilising only geometry (right). **(l)** Comparison of population-averaged and cell-specific inference, both in the directed and undirected settings, for switch system using graph construction from PCA coordinates (manifold) and pseudotime coordinates (pseudotime). We show for reference PIDC [28], an information-theoretic approach closely related to locaTE that does not use geometric or context-specific information.

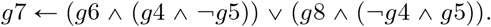

Inference of higher-order interactions is known to be more challenging than inference for pairwise interactions [16, 28]. Cell-specific networks allow us to disentangle interactions for a certain subclass of higher-order interactions: those which can be understood to be locally first-order, conditional on information that is encoded in the cellular context.

LocaTE depends on a user-specified transition kernel *P*. In order to gain insight into how inference accuracy depends on this choice, we consider several choices for constructing the transition kernel: where direct estimation of the velocity is feasible, we try to construct *P* using dot-product (velo-dot), cosine (velo-cos) and Pearson correlation (velo-corr) similarities [44], which are common options available in RNA velocity analysis pipelines [23, 24, 37]. In the absence of direct velocity estimates, trajectory inference methods are used to infer dynamics: we consider constructions of *P* based on diffusion pseudotime [38] (dpt), optimal transport [39] (statot), population balance analysis [21] (pba). Finally, we consider a reversible random walk construction that does not use dynamics information (undir). We summarise in brief the assumptions and limitations of dynamical inference approaches in Box 2 and provide a more detailed discussion in Section 4.

Figure 3(b) illustrate the inference results at the cell level, in the case of direct quantification of dynamics and using a dot-product velocity kernel. For the sake of visualisation, cells are ordered along the trajectory and each cell-specific network is shown as a row. Comparing to the ground truth, we find that locaTE is able to resolve the differential activity of interactions across the continuous trajectory. Figure 3(c) shows cluster-averaged networks for each branch corresponding to one of the two regulatory modules A and B – importantly, locaTE correctly recovers the mirrored regulation pattern between the two branches of the bifurcation from the cell state distribution alone, without using any clustering information. To quantitatively gauge accuracy, we compute area under the precision-recall curve (AUPRC) for cell-specific interaction scores with respect to the simulation ground truth. In Figure 3(d) we summarise the AUPRC ratio relative to a random predictor for cell-specific network inference using locaTE, for each choice of dynamical inference method. As a baseline, we consider a static network oracle in which the ground truth static network is *known* and every cell is predicted to have the static network. We reason that this represents the best-case prediction scenario possible for any static network inference approach. LocaTE achieves a performance gain over the static oracle in every case, giving us confidence that locaTE is able to detect true signal for cell-state specific interactions.

We next consider a scenario where differential regulation occurs along a linear trajectory instead of a bifurcation. This inference problem is more subtle than a bifurcating process given the lack of a clear branching or clustering structure upon which regulatory interactions depend. We consider a model network depicted in Figure 3(e): this network either contains a feed-forward loop *g*2 → *g*3 → *g*4, or *g*2 → *g*4 → *g*3, depending on the position of a cell along the trajectory. The interaction *g*2 → *g*4 is first *indirect* and then becomes *direct*: the nature of this differential regulation is such that static GRN inference methods would observe a superposition of these interactions in the best case scenario, i.e. *g*2 → *g*3 → *g*4 as well as *g*2 → *g*4. As an example, this could potentially lead to the *g*2 → *g*4 interaction being erroneously identified as an indirect interaction and thus being pruned, if filtering heuristics such as CLR or the data-processing inequality were used [17, 45]. Cell-specific networks may be able to disentangle these two effects.

Applying locaTE as in the previous example, we find from Figure 3(f) that the inferred cell-specific interactions again accurately capture the true directed interactions. Cluster-averaging of inferred networks (Figure 3(g)) clearly uncover the network rewiring. We demonstrate again that locaTE offers significant performance gains relative to the static network oracle in terms of AUPRC for cell-specific inference in Figure 3(h).

LocaTE-NMF is an extension of locaTE that allows for unsupervised discovery of regulatory modules from cell-specific networks (see Section 5). Applying locaTE-NMF to both the bifurcating and switch systems, Figure 3(i, j) shows a graph layout of the recovered regulatory modules. For the bifurcating system, we illustrate the network modules that drive the bifurcation, as well as Modules A and B which correspond to each branch. For the switch system, we label modules corresponding to regulatory logic when the switch condition *g*6 ^ (⌐*g*5) is, respectively, on and off. In contrast to the visualisations in Figures 3(c, g) which required prior cell clustering from gene expression, these results illustrate how locaTE-NMF can identify distinct regulatory states in an unsupervised manner independent of clustering.

In Figure 3(k) we show that locaTE can be applied *without* dynamics information, using an *undirected* transition kernel relying only on a graph approximation to the cell-state manifold 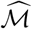. This construction assumes that the cell population is at equilibrium. Where information on the dynamics is available, directed interactions are inferred using a transition matrix to capture these dynamics. When no dynamical information is available, using the undirected transition kernel reveals the presence of interactions but cannot extract causal relationships since temporal information is lost under an equilibrium assumption.

Finally we investigate the importance of accounting for the cell state manifold, rather than just orderings. The switch-like network exhibits a linear trajectory topology, allowing for a fair comparison against pseudotemporal ordering. We apply locaTE to the same simulated data as previously, but construct neighbourhood kernels and *k*-NN graphs based on the true simulation time of each sampled cell instead of PCA coordinates. We reason that using the ground-truth simulation time for the ordering provided a “best-case” scenario for pseudotime-inference approaches. In Figure 3(l) we compare the performance of locaTE using PCA-based manifold approximation (locaTE-manifold) against locaTE using pseudotime neighbourhoods (locaTE-pseudotime) as well as PIDC [28] as an additional baseline. We consider the directed setting, using velocity information with the dot-product kernel, as well as the undirected setting using an undirected transition matrix. Parameters for each case are kept the same as for the previous examples. LocaTE-manifold outperforms locaTE-pseudotime significantly in all cases: the performance gain is especially dramatic for cell-specific inference, with the AUPR roughly doubling. Of particular interest is that locaTE-manifold outperforms PIDC for the task of *undirected static* network inference: locaTE-manifold does not use velocity information in this setting, and so has exactly the same input data as PIDC. This provides further evidence that appropriately modelling the underlying geometry of cell states is crucial to obtaining good results for network inference.

Together, our simulated network analyses demonstrate that locaTE is able to combine dynamical and geometric information to infer both cell-specific and static networks, often with a substantial margin compared to existing approaches. Unsurprisingly, better estimates of dynamics lead to better results for network inference. Perhaps what is more remarkable is that, even after aggregating cell-specific networks produced by locaTE, the resulting static networks generally outperform competing static inference methods (see Figures S1(d, e), S2(d, e)).

### 2.3 Systematic benchmarking of algorithm performance

To quantitatively demonstrate the advantages of dynamical and geometric information, we carry out a systematic comparison of locaTE against a selection of existing network inference methods across an array of test instances. In addition to the two systems from Section 2.2, we consider a range of systems with varying trajectory behaviours including linear, multifurcating, cyclic and combinations thereof. In total, we construct 24 test systems across two independent trajectory simulation tools. We simulate 17 systems using BoolODE [6] which uses a continuous, stochastic differential equation (SDE) formalism for modelling gene expression dynamics. These systems include the instances from Section 2.2, networks presented in existing literature [6, 8], as well as additional instances. The remaining 7 systems are simulated using dyngen [10], which models single cell gene expression at the level of discrete counts using the Gillespie algorithm. We show in Figure 4(a) examples of these trajectories and their corresponding networks. For trajectories simulated using dyngen, dynamics in the form of a cell-state transition matrix were estimated from spliced and unspliced counts using the dynamo analysis pipeline [24].

**Figure 4:**
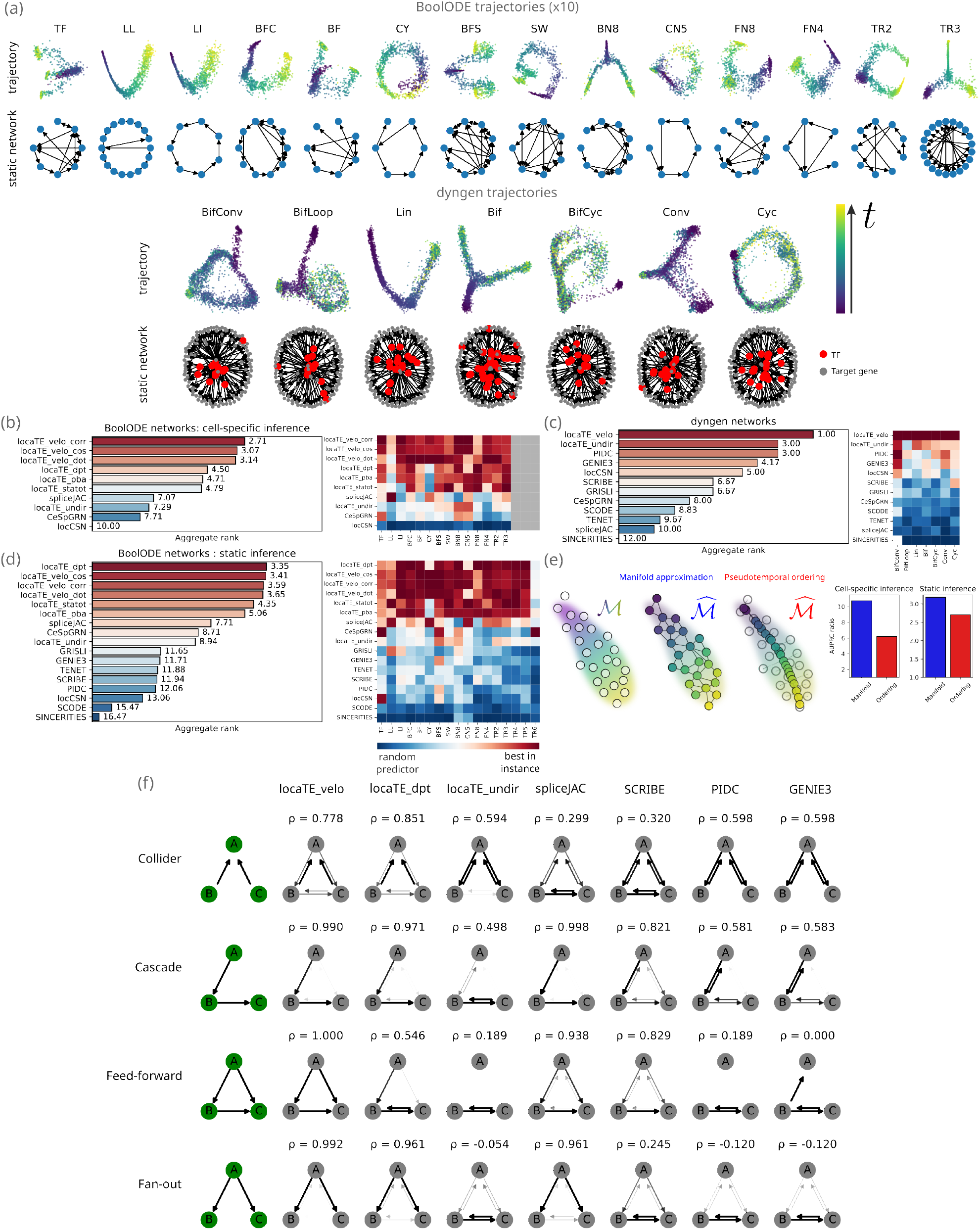
Systematic benchmarking instances and results. (a) Dimension-reduced embeddings of 14 out of 17 BoolODE trajectories (top) and 7 dyngen trajectories, colored by the true simulation time, shown alongside ground truth static networks. For dyngen instances, TFs are shown in red and target genes in grey. (b) Cell-specific inference summary for BoolODE networks. (c) Static inference summary for dyngen networks. (d) Static inference summary for BoolODE networks. (e) Effect of graph manifold approximation versus pseudotemporal ordering for network inference accuracy. (f) Summary of inference performance for 3-node network motifs across BoolODE static networks. 33

For cell-specific network inference, we compare to CeSpGRN [11], locCSN [12] and SpliceJAC [14]. SpliceJAC uses spliced and unspliced count data to infer directed networks; CeSpGRN and locCSN do not model dynamics and are thuslimited to inferrence of undirected networks. Since cell-specific networks inferred by locaTE can be aggregated to produce population-level “static” networks, we considered accuracy for inference of static networks as well as for cell-specific networks. From the many static network inference approaches in the literature, we select 7 methods to compare to. Our selection is based on the relevance of the method to ours, as well as previous benchmarking results from [6], and we provide detailed descriptions and rationale for method selection in Section 5. In summary, we compare locaTE to TENET [17] and SCRIBE [15], which use transfer entropy with pseudotime information to infer directed networks; and PIDC [28] and GENIE3 [46], which were found to perform best overall in the prior benchmarking study [6]. We additionally consider SCODE [18], GRISLI [20] and SINCERITIES [47], which use dynamical information to infer directed networks.

We apply locaTE to each system using the transition kernel constructions summarised in Section 2.2. Inference accuracy is measured in terms of the AUPRC ratio, which corresponds to the gain in performance relative to a random predictor. In Figure 4(b-d) we show results for cell-specific and static network inference in BoolODE and dyngen trajectories. To construct an overall ranking of inference algorithms across all test instances considered, we rank methods for each test instance and aggregate across instances using the Borda count method; see Section 5 for further details on benchmarking.

Our results show that locaTE consistently performs well across systems of different sizes and with different trajectory topologies, both for cell-specific and population-averaged, static inference. For cell-specific inference, we find that locaTE with velocity-based kernels perform best, followed by trajectory inference-based kernels. SpliceJAC, CeSpGRN and locaTE-undir show mixed results. In particular, we remark that CeSpGRN and locaTE-undir show similar patterns of accuracy across test instances, with locaTE-undir having a slight performance edge. This is interesting since neither method uses dynamical information, but both use a context-specific inference approach. This lends additional evidence to the benefit of context-specifc inference, as was observed in Figure 3(l). We find that locCSN consistently performs poorly.

Considering population-averaged static network inference, we find that all locaTE methods that use dynamics information perform strongly. While SpliceJAC performs relatively well in systems simulated using BoolODE, it performs consistently poorly in the dyngen instances. Inconsistent performance may be due to differences in the simulator that can violate assumptions of the inference method. Since SpliceJAC is developed by linearisation of a kinetic model of splicing dynamics [14], the inherent assumptions of this kinetic model is a likely culprit. Again, we find that locaTE-undir (locaTE with an undirected kernel) either outranks or is on-par with PIDC and GENIE3 overall for both BoolODE and dyngen trajectories. Since all of these methods use the same input data without dynamics information, this provides evidence indicating that network inference in general may benefit from a cell-specific approach, independently of dynamics information.

To illustrate the advantage of using a graph approximation to the cell state manifold, we ran locaTE using a dot-product velocity kernel for the SW and BFS trajectories (Figure 3) simulated by BoolODE, using a *k*-NN graph constructed from either PCA embeddings or pseudotemporal ordering of cells. We illustrate the difference between these approaches in Figure 4(e): using the more flexible graph approximation (manifold) allows more subtle aspects of the data to be captured, compared to using only an ordering along a unidimensional axis of variation (ordering) as provided by principal curves, pseudotemporal ordering, or similar. For manifold approximations we observe major improvements in inference accuracy compared to pseudotime, with a nearly twofold improvement in accuracy for the cell-specific setting, as measured by AUPRC ratio.

As an additional perspective on inference performance for different methods, we show in Figure 4(f) four archetypal 3-node motifs and corresponding inference results summarised across 12 BoolODE trajectories (excluding TR2-6 due to network size) for locaTE, SpliceJAC, SCRIBE, PIDC and GENIE3. We measure accuracy of motif recovery in terms of the Pearson correlation between the inference output and the ground truth adjacency. Overall we find that locaTE achieves a high level of accuracy for motif recovery. In particular, we find that locaTE-velo has a significant advantage over locaTE-dpt in the case of the feedforward loop – this suggests that direct measurement of dynamics may help to resolve direct interactions from indirect ones, while dynamical inference may at times fail to be sufficiently accurate for this purpose. We find that locaTE-undir again performs similarly to PIDC and GENIE3; the collider motif is the most challenging to infer accurately.

Overall, our benchmarking results showcase the consistently strong performance of locaTE across different simulators, trajectories, and topologies.

### 2.4 Embryonic stem cell to primitive endoderm differentiation

We first consider the dataset of Hayashi et al. (2018) [48], consisting of 456 cells and 100 TFs. This dataset is a time-series of mouse embryonic stem cells differentiating through to primitive endoderm, sampled at timepoints 0, 12, 24, 48, 72h. Cells in this differentiation process follow a linear trajectory, as can be seen in Figure 5(a). To infer trajectories, we apply StationaryOT [39] using quadratically-regularised optimal transport to ensure a sparse transition kernel (see Methods for details). Using this transition matrix, we infer cell-specific networks with locaTE, resulting in a 456 × 100 × 100 array of interaction scores.

**Figure 5:**
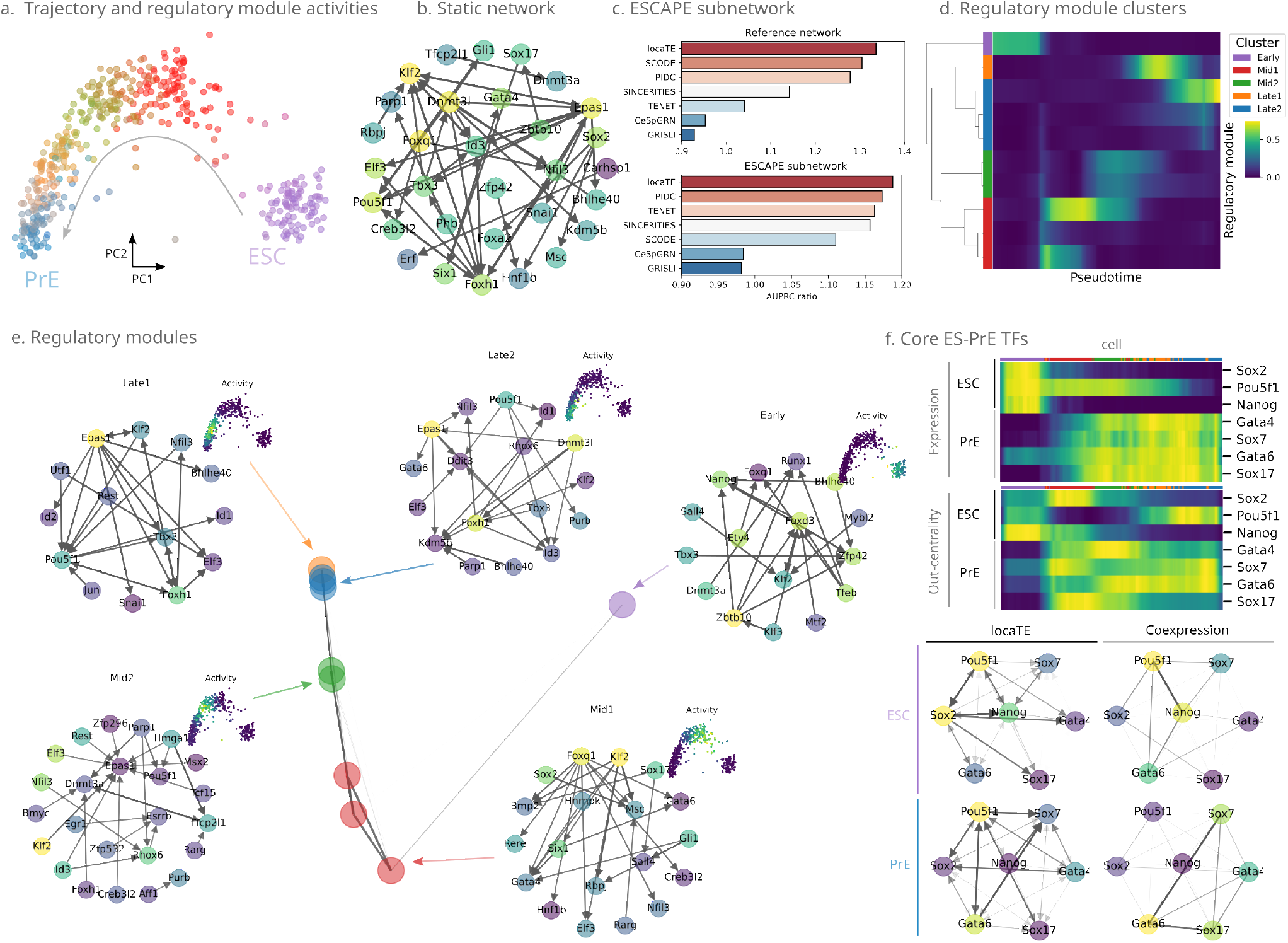
ES-endoderm development dataset. **(a)** PCA embedding of 456 cells along an embryonic stem cell (ES)-primitive endoderm (PrE) differentiation trajectory, coloured by soft assignment to regulatory modules. **(b)** Static network produced by averaging locaTE cell-specific networks (top 0.5% of all edges by confidence). **(c)** Area under precision-recall curve (AUPRC) ratios against (i) the full reference network constructed by [18] and (ii) the ESCAPE reference subnetwork [49]. **(d)** Regulatory module activities found using locaTE-NMF, clustered by activity along the linear differentiation trajectory. **(e)** Graph layout of regulatory modules found by locaTE-NMF coloured by cluster together with inferred networks (top 0.25% of all edges by confidence) averaged over each cluster. Network nodes (TFs) are coloured by out-edge eigenvector centrality. Overall activity for each cluster of modules is shown (inset). **(f)** Set of 7 core transcription factors known to be involved in murine ES-PrE differentiation, shown (i) by expression level and out-edge eigencentrality over the differentiation process and (ii) subnetworks corresponding to ESC and PrE celltypes, compared to co-expression (Pearson correlation) subnetworks. See main text for detailed discussion.

Since no ground truth is available for cell-specific networks in experimental datasets, we can only validate the output of locaTE against a static network benchmark. Unfortunately, “ground truth” reference networks are generally not available. Even when relevant data exists, there is no certainty that they reliably reflect the *activity* of interactions given the experimental conditions and developmental timepoints captured in a particular experiment. For instance, reference networks constructed from epigenetic accessibility assays or TF-binding motifs provide a set of *biophysically possible* interactions, but do not tell which of those interactions are in fact actively influencing cell dynamics. Notwithstanding, we provide some brief results for two reference networks that were previously used in the context of this dataset. In [18] the authors use a reference network constructed from DNaseI footprint data, and in [19] the authors constructed a reference network from known regulatory interactions in the embryonic stem cell atlas from pluripotency evidence (ESCAPE) database [49] which consists of 19 regulators and 100 targets. Starting from the array of cell-specific interaction scores, we obtain a static 100 × 100 GRN by averaging interaction scores across cells and compute precision-recall curves each benchmark network (Figure 5(c-d)). Comparing to CeSpGRN, TENET, SCODE, GRISLI, PIDC and SINCERITIES, we find that locaTE performs comparably to, or better than, other methods in terms of AUPRC for each reference network. We additionally find that other information theoretic methods (PIDC and TENET) show strong performance, especially with PIDC consistently among the best performers for both reference networks.

Since locaTE infers cell-specific networks, we can ask questions at a much finer resolution than possible at the population level. To demonstrate this we use locaTE-NMF with *α* = 0.75 to find *k* =10 regulatory modules that are active at different stages along the trajectory (Figure 5(d)). Hierarchical clustering of the inferred factor activities produces five groups, which we label as “Early”, “Mid(1,2)”, and “Late(1,2)” trajectory stages. The inferred network for each cell can be expressed as a weighted linear combination of each of the these modules. In Figure 5(a), cells are coloured by their regulatory module activities, highlighting that the inferred regulatory modules are specific to different regions of the trajectory. For each cluster we compute a representative network by taking weighted averages of the NMF factors, and the top 0.25% confidence edges for each stage are shown in Figure 5(e). It is clear that the nature of the top inferred regulatory interactions changes drastically along the trajectory. Included among these are the canonical ones previously reported to be associated with maintenance of pluripotency and embryonic development, including Nanog-Zfp42 [50], Foxd3-Nanog [51] and Sall4-Klf2 [52] which are active early in the trajectory. Top inferred interactions for the late trajectory include Epas1 (Hif2a)-Pou5f1 (Oct4), which is known to play a role in embryonic development [53]; and Epas1 (Hif2a)-Bhlhe40 (Dec1), which plays a role in suppression of Deptor [54], a stemness factor known to be downregulated during ES differentiation [55].

For each stage, we rank potential regulators by their out-edge eigenvector centrality and show the 25 top-ranked genes for each stage (see Figure S10(a)). We find that many of the top genes are known to be associated with stem cell development, including Nanog, Pou5f1 (Oct4), Sox2 [51], Gata6 [56], Zfp42 [50]. These gene lists are input to Metascape [57] to find corresponding Gene Ontology (GO) annotations (Figure S10(b-d)). Among the gene set annotations found for the “Early” and “Mid1” trajectories, we find that enrichment for pluripotency and embryonic morphogenesis. Annotations for the “Mid2” and “Late(1,2)” trajectories include epithelial cell differentiation and organ development. This reflects the developmental progression of embryonic stem cell differentiation to primitive endoderm and suggests that locaTE is able to identify key molecular drivers of the underlying process.

Finally, we focus on a subnetwork of 7 core transcription factors known to drive different stages of the ES-PrE differentiation process [58]. In Figure 5(f) we contrast the measured gene expression to the out-centrality of the corresponding node in the cell-specific locaTE network. We find that the patterns of gene expression and regulatory activity are highly distinct, indicating that locaTE is able to extract signal from the dynamics of gene expression rather than just reflecting expression or coexpression level. We show cluster-averaged subnetworks for the ESC (earliest) and PrE (latest) states, further illustrating the difference between locaTE networks and simple coexpression networks. Notably, in the locaTE network Pou5f1 (Oct4) is predicted to have a strong affinity with Sox2 in the ESC cluster, while in the PrE cluster it is predicted to additionally interact with Sox17 and Sox7. It was reported by [59] that Oct4 switches binding partners from Sox2 (in ESC) to Sox17 (in PrE), and that Sox7 and Sox17 may behave in a similar, redundant manner.

### 2.5 Pancreatic development

Next we consider the murine pancreatic development dataset of Bastidas-Ponce et al. [60]; here cellular dynamics are captured by RNA velocity estimates. The dataset consists of 2,531 cells, and we filter for 82 transcription factors following the preprocessing of [20]. We apply the scVelo [37] package to produce RNA velocity estimates, and CellRank [23] with a cosine kernel to produce a transition matrix encoding the inferred dynamics on the cell set. In Figure 6(a) we show a UMAP embedding of the full dataset with streamlines visualising the estimated dynamics. We apply locaTE with the RNA-velocity derived transition matrix, and obtain a 2, 531 × 82 × 82 array of cell-specific interaction scores. To illustrate the heterogeneity in the networks inferred by locaTE, we colour cells by their corresponding cell-specific GRN using the first diffusion component of the inferred cell-specific networks (see Figure S12(b)). This reveals a clear gradient along the differentiation trajectory, suggesting that the networks inferred by locaTE capture a progression in regulatory interactions over developmental time. The highest-scoring interactions involve the genes Fos and Egr1, which are known to play a role in stress response [62]. Fos expression has been documented to be an dissociation-induced artifact in scRNA-seq [63], and we therefore conservatively filter out interactions involving Fos or Egr1 in subsequent analyses and for visualisations.

**Figure 6:**
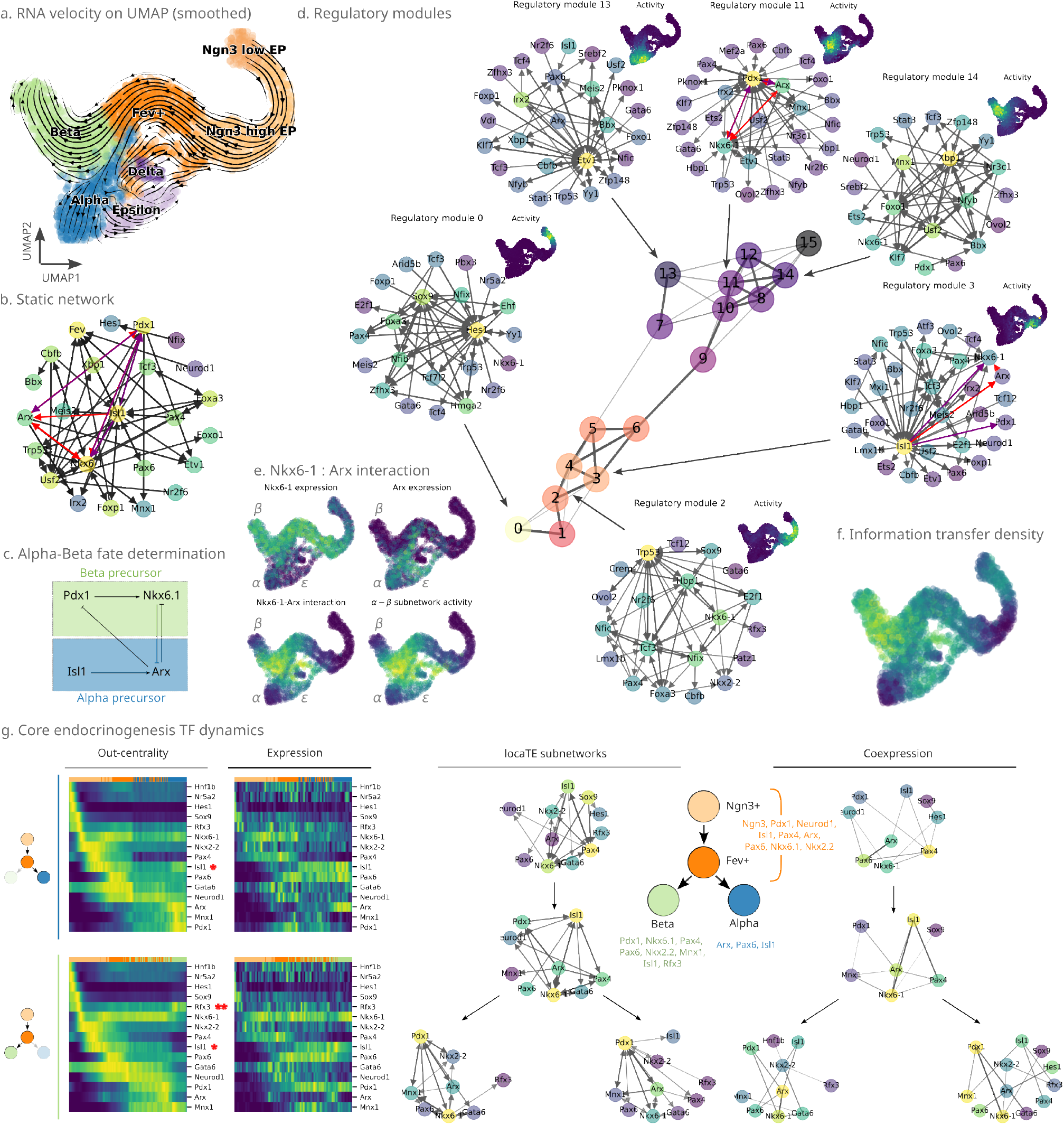
Pancreatic development dataset. **(a)** UMAP embedding of murine pancreatic development dataset [60], with smoothed RNA velocity vector field shown as a streamplot. Cells are coloured according to their cluster annotation from the original publication. **(b)** Static GRN constructed as population average of cell-specific networks (top 1% of edges by confidence shown, after filtering for stress response genes). **(c)** Alpha-beta fate determination network of [61]. **(d)** Graph layout of regulatory modules found using locaTE-NMF, coloured by proximity to early trajectory. Selected regulatory modules are shown (top 1% of edges shown by confidence). **(e)** Nkx6.1-Arx expression and interaction scores in UMAP coordinates, and average activity of the 4-gene *α*/*β* subnetwork of [61]. **(f)** Information transfer density (root-mean-square of total TE scores) shown in UMAP coordinates. **(g)** Core endocrinogenesis transcription factors: (i) shown along pancreatic alpha and beta cell lineages by out-edge eigencentrality and expression level, (ii) subnetworks corresponding to Ngn3+, Fev+, Beta and Alpha clusters found by locaTE, compared to coexpression (Pearson correlation) networks. See main text for detailed discussion.

As a first summary of the inference output we produce an 80 × 80 static GRN by population averaging. Figure 6(b) shows the top 1% confidence edges. The genes with highest out-edge eigenvector centrality include Nkx6-1, a known master regulator of pancreatic beta cell development [61, 64], as well as Isl1 and Pdx1, known determinants of alpha and beta cell fate [65]. In [61] the authors report evidence for a 4-gene network (henceforth referred to as the *α*/*β* network) involving Nkx6-1, Pdx1, Isl1 and Arx, that drives alpha and beta fate determination (Figure 6(c)). All 4 genes have high out-edge centrality in the inferred static network, and, furthermore, we find that the *α*/*β* network is recapitulated in the top-1% edges inferred network, as highlighted by the coloured edges in Figure 6(b, c). Comparing to the networks inferred by PIDC and TENET (Figure S14), we find that both approaches detect aspects of the same fate-determination network within the top scoring edges.

Applying locaTE-NMF with *α* = 0.75, we find 16 regulatory modules spanning the set of transcriptomic states. Computing pairwise similarities between regulatory modules as before, we show in Figure 6(d) a graph layout of the learned regulatory modules in which each node is coloured by its activity over pseudotime. From these we illustrate six modules (top 1% edges by confidence), which correspond to key features of the developmental trajectory. Regulatory module 0 is active near the Ngn3 − EP (endocrine progenitor) cell cluster, including a predicted interaction Sox9–Hes1. Sox9 is known to play a key role in maintaining the pancreatic progenitor cell state by regulating Hes1 [66], an inhibitor of Neurog3 (Ngn3) [65]. This interaction is no longer present in regulatory module 2, which is most active near the Ngn3+ cell cluster, further evidence for its role in maintaining the Ngn3 − state. This interaction does not appear in the static GRN in Figure 6(b), illustrating the need for cell-specific networks in capturing localised signals, which would be lost in static networks or population averaging. Out-edge eigenvector centrality of module 0 identifies additional potential regulators including Foxa3, known to be implicated in endodermal organ formation [67]; and Hmga2, a chromatin-associated protein expressed during development that does not directly regulate downstream genes, but appears to have widespread indirect effects via chromatin interactions [68].

Modules that correspond to alpha and beta committed cells yield networks that are distinct from each other as well as from the progenitor clusters. The activity of module 13 is concentrated near the alpha cluster: examining out-edge eigenvector centrality, we find that Irx2 is featured prominently, and has been suspected to be involved in alpha fate specification [69]. In module 14, which is concentrated near the beta cluster, we find that Xbp1 is a top regulator recently found to be critical to maintenance of beta-cell identity [70].

Not all regulatory modules correspond to annotated cell type clusters, however: for instance, regulatory interactions that drive fate determination may instead be active at the boundary between cell states (thought of as a saddle point between two attractors in the Waddington’s landscape, see Box 1). We highlight module 11, which is active at the boundary between the alpha and beta lineages. In this module we find interactions between Pdx1, Arx and Nkx6.1 to be among the top inferred edges. This module is consistent with a fate decision boundary between the two lineages. Additionally in module 3, located between the Ngn3+ EP and Fev+ clusters, Isl1 is identified as a top regulator along with Isl1-Arx. This agrees with the early behaviour of the fate determination network of [61] in which Isl1 activates Arx (consistent with module 3), which in turn competes with Nkx6.1 (consistent with module 11). To investigate this at a finer scale we visualise expression levels and interaction scores for the Nkx6.1-Arx interaction in Figure 6(e). As before, we observe a fate decision boundary from expression levels that reflects the competition between Nkx6.1 and Arx, corresponding to beta and alpha fate. The interaction scores for both the Nkx6.1–Arx edge and the entire *α*/*β* subnetwork are strongest near this boundary.

Finally, we consider a subset of 15 TFs implicated in pancreatic alpha and beta cell development from endocrine progenitors [60]. In Figure 6(g) we contrast regulatory activity (as measured by node out-centrality) against gene expression for each of the alpha and beta lineages. As in the ES-PrE example, we find that the out-centrality patterns are different from gene expression across the trajectory. Notably, the alpha lineage driver Isl1 (*) shows a decreased regulatory activity in the beta trajectory compared to the alpha trajectory, and this trend is not discernible from the gene expression alone. On the other hand, Rfx3 (**), implicated in beta cell development, shows an increased activity in the beta trajectory compared to alpha. We construct cluster-averaged subnetworks for these TFs for the Ngn3+, Fev+, Beta and Alpha cell states, and compare to the corresponding coexpression networks. In the locaTE subnetworks, Nkx6-1 has a higher regulatory activity in the beta subnetwork, while Arx is more active in the alpha cell state. This is in agrement with biological prior knowledge (Figure 6(c)). We find that this trend is reversed in the coexpression networks, with Arx having a higher centrality in the beta subnetwork, and Nkx6-1 having a higher centrality in the alpha subnetwork.

### 2.6 Haematopoiesis

Haematopoiesis is a biological process that has long been regarded as a canonical system for studying complex cell fate decisions and is far from being fully understood [71]. In what follows we apply locaTE to one simulated and one real dataset modelling this process.

First we consider a simulated dataset using the literature-derived gene regulatory network from [6, 72], which models the differentiation of Common Myeloid Progenitors (CMPs) towards Monocyte, Granulocyte, Megakaryocyte and Erythroid lineages. As with our earlier benchmarking study, rigorous analysis of network inference performance is only possible in simulation settings where the ground truth system is fully specified. We simulate 2, 500 cells from the system (Figure 7(a)) and run locaTE using a transition kernel derived from simulated velocities and a cosine similarity kernel. Using locaTE-NMF with *α* = 0.6 we derive *k* =5 regulatory modules, and we display cells colored by their weights towards each regulatory module. This illustrates the variation of the inferred networks across the trajectory and, notably, the branching of distinct lineages. We illustrate one regulatory module associated to the CMP stage, showing the top 10% of edges and find that a large fraction of the top inferred edges are genuine. Of note, PU.1 (Spi1) is assigned the highest regulatory activity in terms of out-edge eigencentrality, reflecting its well known central role in the Myeloid-Erythroid fate determination process [71, 72]. In Figure 7(b) we show the ground truth network along with the inferred static network (top 20% of edges by confidence), in which edges are colored by their region of activity across the trajectory. Finally in Figure 7(c) we compare the static network inferred by locaTE to the alternative inference methods from Section 2.3 in terms of AUPRC ratio. Our results are largely consistent with our benchmarking findings: we find that locaTE out-performs competing methods, with CeSpGRN and PIDC also showing strong performance.

**Figure 7:**
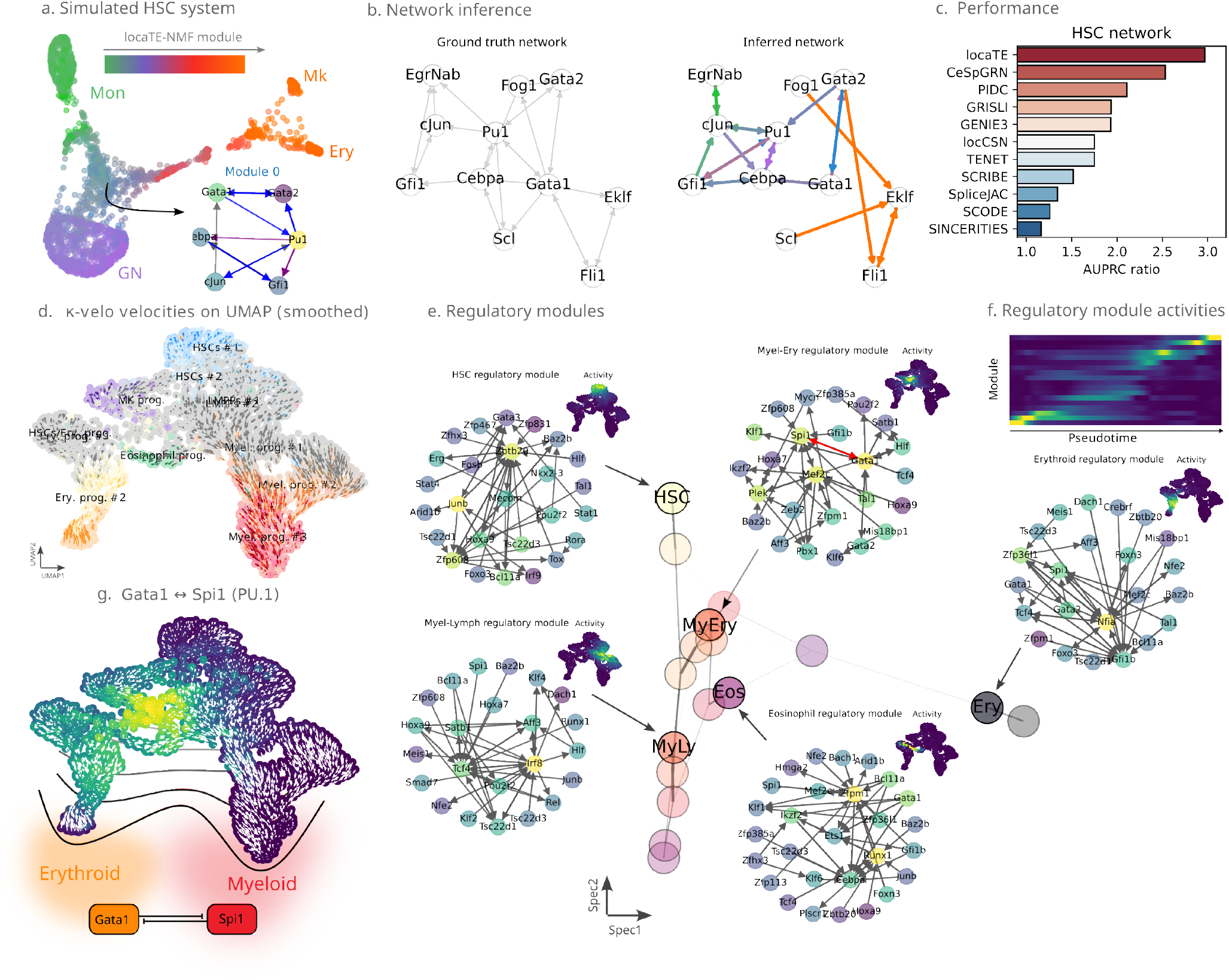
Haematopoiesis systems. **(a)** SPRING layout of simulated cells from HSC system, colored by relative activity of regulatory modules found by locaTE-NMF. Module 0, corresponding to CMPs, is illustrated (top 10% by confidence) with true positive edges colored in blue, and indirect (1-hop) edges colored in purple. **(b)** Ground truth static network for simulated HSC system, compared to the inferred, population-averaged network (top 20% by confidence). Inferred edges are colored by their region of activity. **(c)** Static network inference performance for locaTE and competing methods, as measured by AUPRC ratio. **(d)** Smoothed velocities found by *κ*-velo for the dataset of Marot-Lassauzaie et al. [41] shown on UMAP coordinates, coloured by cluster annotation. **(e)** Graph layout of regulatory modules found by locaTE-NMF coloured by proximity (measured by mean first passage time) from the HSC module, shown alongside representative networks of selected modules (top 0.5% of all filtered edges) corresponding to HSC, Myeloid-Erythroid decision boundary, Erythroid, Eosinophil and Myeloid-Lymphoid. **(f)** Smoothed regulatory module activities shown against pseudotime. **(g)** Average locaTE score for Gata1-Spi1 interactions shown on UMAP coordinates along with smoothed velocities, shown alongside a Myeloid-Erythroid fate determination schematic.

One advantage of employing general, transition-kernel based Markov modelling framework is that it is highly flexible and agnostic to how the dynamics are inferred. Currently, inference, interpretation and downstream use of RNA velocity estimates is a contentious topic [40, 73] that remains an active research area. Marot-Lassauzaie et al. [41] propose *κ*-velo, an alternative method for estimating RNA velocity and provide an example application to a haematopoiesis dataset, where the more established method scVelo [37] fails to recover expected trajectories. We apply locaTE to this dataset using inferred dynamics computed by *κ*-velo, producing a 2, 430 × 100 × 100 array of cell-specific interaction scores (see Section 5 for details). Initial observation of the inferred networks showed that the top inferred interactions were dominated by a handful of TFs: Hmgb2, Hmgb3, Myb, and Myc, all of which are known to be implicated in haematopoiesis [74–76]. For visualisation purposes we excluded interactions involving these TFs to prevent overcrowding of the displayed networks.

A bifurcation in the trajectories between myeloid and erythroid lineages is a prominent feature in this dataset (see Figure 7(a)). Erythroid-myeloid cell fate determination is governed by a genetic switch comprised of mutually antagonistic Gata1 and Spi1 (known also as PU.1) [77] (see illustration in Figure 7(f)). In Figure 7(b) we show the locaTE interaction score for the Gata1–Spi1 interaction alone along with the smoothed RNA velocity vector field. We find that the peak activity corresponds to a “dead zone” in the velocity field, while in the surrounding regions downstream, the velocity fields are coherent and indicative of lineage commitment. This is of practical interest [41], as one interpretation of this could be that it corresponds to cells which have high fate plasticity and are still in the “decision-making” process at a saddle node in the epigenetic landscape.

We next apply locaTE-NMF with *α* = 0.75 to find *k* = 18 regulatory modules, and in Figure 7(d) we show a layout of the graph constructed from modules in the same manner as previously done in Figure 6 (see Figure S15(a) for all modules). From this we can see two clear branches corresponding to erythroid and myeloid fate. We find a module (see Figure 7(d), labelled Myel-Ery) that appears concentrated near the bifurcation point on the UMAP coordinates. This module is centered around activity of Spi1 and Gata1, as well as Mef2c, which is known to be involved in lymphoid-myeloid fate determination [78], and locaTE-NMF predicts the key Gata1-Spi1 (PU.1) interaction (edge highlighted in red) as one of the top interactions. In Figure 7(d, e) we display other regulatory modules that correspond to various degrees of fate commitment, including haematopoietic stem cells (HSCs). These networks uncover known hallmarks of haematopoietic development: the myeloid-lymphoid progenitor (labelled Myel-Lymph) module prominently features Irf8, understood to play a key role in the development of myeloid and lymphoid lineages [79, 80] (note that the lymphoid lineage is not captured in this dataset). The erythroid module is centered on a different set of genes such as Gfi1b [81]. The HSC module features Junb prominently, which has been implicated in homeostasis in the long-term HSC population [82]. The HSC module contains interactions involving Baz2b, a master regulator which has been found to reprogram lineage-committed cells to a multipotent state [83].

In the static network produced by aggregating locaTE cell-specific networks (see Figure 7(c)) we find that interactions such as Gata1–Spi1 are obscured due to the population averaging. Comparing to the output of PIDC and TENET (Figure S17), we find that PIDC successfully uncovers this interaction within the top 1% confidence interactions, while TENET misses it. On the other hand, Gata1–Spi1 features prominently in the interaction module uncovered by locaTE, with Gata1 and Spi1 being among the top four nodes by out-edge centrality in that module. Similarly, in Figure S16 we show cluster-specific networks produced by aggregating locaTE outputs across annotated cell types, and it is clear that Gata1-Spi1 does not appear among the top cluster-averaged interactions. This again illustrates the potential for important interactions to occur at the boundary between two defined cell types (and therefore missed by cluster-wise analyses), and the value of locaTE-NMF as an unsupervised approach for finding them.

We additionally applied locaTE to the dataset of Tusi et al. [84] which profiled mouse haematopoiesis and performed dynamics inference using population balance analysis (PBA) [21]. This example illustrates the versatility of locaTE in that it can use any dynamics inference approach, rather than being limited to just a single class of methods like RNA velocity. Similarly to the previous experimental example, this dataset (Figure S19(a)) captures a continuum of cell states from HSC to any one of six lineages: erythrocyte, basophil, megakaryocyte, lymphocyte, monocyte, and granulocyte. Using the PBA potential inferred by the analysis in [84] (Figure S19(a)(ii)), we construct a transition matrix encoding the estimated dynamics. Applying locaTE, we find *k* = 8 regulatory modules with varying activities across the cell population. The inferred modules highlight key drivers for developing lineages, some of which overlap with the Marot-Lassauzaie dataset. Notably, the Gata1-Spi1 interaction is once again detected in a module spanning the myeloid and erythroid fate boundary (Myel-Ery), along with Klf1 as a top regulator. Due to discrepancies in captured celltypes between the Marot-Lassauzaie and Tusi datasets, direct comparison of the inferred networks and modules is not possible or appropriate. However, for overlapping genes between the two analyses we calculate the similarity between regulatory modules found by locaTE-NMF in each dataset: this is calculated as the Spearman rank correlation between gene eigencentralities (Figure S19(c)). We find that many of the uncovered regulatory modules are similar between datasets, suggesting that locaTE is able to produce biologically consistent results across datasets and dynamical inference methods.

## 3 Discussion

Biology is dynamic. A static representation of a network can never do justice to what we observe phenotypically. Discerning molecular causes requires experimental and theoretical advances to progress in lock-step. The power of single cell data, and the insights we can gain from such data, rely on flexible statistical analyses that allow us to dissect the differences between gene regulatory networks (and programs) and how they differ between cells. Gaining deeper mechanistic insights into single-cell data requires a framework to reconstruct cell-specific, causal networks. Our method, locaTE, computes the transfer entropy (TE) [27, 85] for pairs of genes to detect causal relationships. The approach uses approximations of the underlying dynamics but makes no extraneous assumptions on linearity of interactions, the distribution of gene expression counts, or the topology of the trajectory.

By using neighbourhoods in gene expression space we can infer cell-specific networks without going to the extreme *N* = 1 case. How to pool information across cells most efficiently will ultimately decide on the success of such methods and the insights they provide. Temporal ordering helps enormously, but we are still, by and large, not able to collect meaningful single cell transcriptomic time-course data. Pseudo-temporal ordering or some of the many versions of RNA velocity entering the literature can provide useful information, as can, related multi-omic data, such as scATAC-seq.

LocaTE is not limited to any single dynamical inference method: while RNA velocity data are currently among the most abundant in the literature, metabolic labelling datasets hold the promise of more accurate vector field quantification. Nor is our approach limited to inference for transcription factor networks: as an information theoretic network inference approach, locaTE aims to uncover information flow between observed random variables. Thus locaTE can in principle be applied to any gene expression data, but also surface marker and protein expression data. As an illustration, in Section S1.9 we applied locaTE to a metabolic labelling dataset of the cell cycle [86] to infer context-specific and causal coexpression networks between 200 genes found to be highly correlated with cell cycle. This highlights the flexibility of locaTE: using a learned vector field based on metabolic labelling data [24], non-linear trajectories can be handled with ease.

As with any other statistical inference method, locaTE relies upon underlying modelling assumptions: the Markovian model of cellular dynamics, and consequently the information theoretic approach for quantifying statistical dependency between measured variables. As a result, we identify several potential limitations that may affect the applicability of locaTE (or indeed, other related network inference approaches) in different datasets. Among others, the manifold assumption underlying locaTE requires the presence of continuous variability of observed gene expression states in a single cell population. LocaTE therefore is most suited to single-cell snapshot experiments, where cells are sampled from across a developmental spectrum. Time-series datasets in which there exist large “gaps” in gene expression space between successive observations may thus not be amenable to analysis using locaTE. We provide further discussion of these issues in Section 5.

Directions for future work include investigating approaches using which local information can help dealing with technical zeros (dropout) in single cell data. We investigated the application of data-imputation methods for the pancreatic dataset; however, we found that imputation worsened performance for all inference methods considered. This effect has been reported previously to be due to spurious signals being amplified [87] between non-interacting genes. Additionally, extensions of mutual-information based approaches such as partial information decomposition [28] and part mutual information [88] have been applied to undirected network inference – it remains to be seen whether similar adaptations are possible for directed inference in the transfer entropy setting.

The problem of inferring cellular dynamics, upon which application of methods such as locaTE depend, is far from solved. In particular, accurately modelling and quantifying RNA velocity remains challenging in practice [40, 44]. Data from additional modalities will help greatly: metabolic labelling assays can provide a more accurate source of dynamical information [24], and lineage-resolved data [89] can provide dynamical information beyond that which can be obtained from gene expression data alone. Since locaTE requires only that dynamics are described by a Markov process, we expect that advances in trajectory inference can be used to inform cell-specific network inference. Finally, a further route for improving locaTE and network inference in general is to use relevant prior information to guide network inference.

## 4 Theory

In this section, we provide an introduction to key concepts underpinning cell dynamics reconstruction and information theory which motivate the design choices behind locaTE. Within each subsection, we provide references to articles and textbooks which may provide the interested reader with a more in-depth discussion.

### 4.1 Markov process approximations of cell dynamics

At the core of the locaTE algorithm is the use of a Markovian model of the transcriptional dynamics of cells in a biological process, a framework which has been increasingly adopted for modelling cell dynamics at high (e.g. single-cell) resolution [21–24, 39]. For a set of measured cells 𝒳 = {*x*_1_, …, *x*_| *𝒳* |_} ⊂ ℝ^*d*^ where *d ≫* 1 is the dimensionality of the system (i.e. the number of measured genes), the Markov chain is specified by a | 𝒳 | × | 𝒳 | transition matrix *P* which corresponds to a discrete time random walk on 𝒳. The primary modelling assumption of locaTE is that the true dynamics can be adequately described as a Markov process. Theoretically, this assumption is valid for an observed system when there are no hidden variables [7,21]. Since experimental measurements are always partial observations of cell state, in practice this is an approximation. No assumptions are made on the structure of *P*, and any one of the many dynamical inference methods in the literature can be used to construct *P*. This contrasts with parametric trajectory inference methods which rely on an underlying linear or tree-like structure [90]. In this section we provide a brief overview of some dynamical inference methods, as well as theoretical motivations for their use. Throughout this section, we consider a general model of single cell dynamics described by the SDE

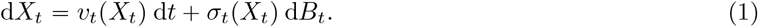

#### Construction from direct measurement of velocity

We first consider the setting where direct quantification of the velocity field *v* is possible – this is the setting with the fewest structural limitations on the form of the reconstructed dynamics [24,44,91], compared to “trajectory inference” approaches to reconstruct dynamics from population measurements, which are restricted by their assumption existence of a potential function or pseudotemporal ordering. Practical approaches for measuring *v* directly at scale include RNA velocity [37, 40, 41, 73], metabolic labelling [24, 86], or lineage tracing [89, 92, 93]. We note that live imaging [15], while being a limited throughput technique, also falls in this category as it allows the ground truth temporal coupling to be observed. In what follows, we provide a discussion of RNA velocity and metabolic labelling approaches since they require relatively sophisticated processing prior to downstream use. Both RNA velocity and metabolic labelling techniques aim to measure the transcriptomic velocity of single cells, in terms of the rate of transcription of genes. Both approaches aim to achieve this by distinguishing between “new” and “old” transcripts together with a kinetic transcriptional model: RNA velocity uses intron sequence information to identify nascent, unspliced mRNAs and use a splicing kinetics model, while metabolic labelling approaches chemically modify nascent mRNAs to allow for unbiased quantification [24].

RNA velocity quantification remains a very active field of research. [40] and [73] provide in-depth discussions of potential pitfalls and failure modes. In summary, quantification methods for RNA velocity rely on assumptions of some underlying model of RNA metabolism as well as kinetic parameters across genes. Additionally, since RNA velocity requires detection of introns, transcript length may introduce additional biases to the inference [94]. A typically observed failure mode of RNA velocity is “backflow” estimated velocity field against the biologically known direction of cell state progression [41] due to kinetic modelling assumptions being violated.

Metabolic labelling approaches provide an experimental route towards direct quantification of nascent mRNAs, circumventing sources of error in RNA velocity, and therefore do not suffer from the same limitations as RNA velocity. While single cell metabolic labelling data for developmental systems are currently relatively scarce due to the need for a new experimental technique, recent works have demonstrated its significant advantages over standard RNA velocity for dynamics modelling [24, 95].

For our purposes, we can be agnostic to the dynamical inference method and assume that we have access to estimated high dimensional cell state and velocity vectors {*x*_*i*_}, *{v*_*i*_}*∈* ℝ^*d*^, whatever their origin. From these, we construct a Markov transition matrix *P*. The task of constructing *P* was investigated in detail from a mathematical perspective by [44, Section 4], who considered several choices of velocity kernels and derived their corresponding differential operators in the continuum limit. We summarise their results below, where we adopt their notation of denoting displacements between expression states by *δ*_*ij*_ = *x*_*j*_ − *x*_*i*_.

- Cosine kernel cos(*δ*_*ij*_, *v*_*i*_): recovers effective vector field *v*(*x*)/‖*v*(*x*) ‖ [44, Theorem 4.1].
- Correlation kernel corr(*δ*_*ij*_, *v*_*i*_): recovers effective vector field *P*_1_*v*(*x*) / ‖*P*_1_*v*(*x*) ‖ where *P*_1_ denotes a linear projection operator *P*_1_*v* = (*I* − **11**^**⊺**^)*v* [44, Theorem 4.2].
- Inner-product kernel x*δ*_*ij*_, *v*_*i*_y: recovers vector field *v*(*x*) + *C*∇_*x*_ log *p*(*x*) where *p*(*x*) is the density of observed cells and *C* > 0 is a constant [44, Theorem 4.3].

We note that there is a bias introduced by the density in the setting of the inner product kernel, in that it is consistent with the superposition of the underlying vector field together with a constraining potential that is proportional to − log *p*(*x*), which is commonly referred to as a “quasipotential landscape” [96] in classical systems biology literature. As we discuss later, this interpretation promises to be useful in the undirected setting.

From a theoretical point of view, the primary advantage of direct velocity quantification is that it enables measurement of the *non-conservative forces* at play in a developmental system, such as cell cycle [24, 37]. Fundamental identifiability issues [21] mean that recovering non-conservative dynamics from population snapshots is typically not possible.

#### Construction from trajectory inference methods

When direct quantification of the velocity field *v* is not feasible or not reliable, a variety of *inference* approaches may be employed to estimate *v* from other observables or prior biological knowledge, such as “starting” cells, estimated growth rates and structure of the underlying cell state manifold ℳ. The problem of recovering *v* in the setting of snapshot datasets was studied in [21], in which the structure of the manifold ℳ (approximated by a *k*-NN graph 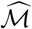) in combination with prior information on regions of influx and efflux of cells was used to infer the underlying flow, via techniques rooted in spectral graph theory. A similar approach based on regularised optimal transport was proposed in [39]. Both approaches are subject to the limitation that inference is restricted to dynamics that are driven by a vector field of gradient type, i.e. *v (x)* = − ∇Ψ (*x*). This limitation is in fact a fundamental one for problems of inferring dynamics from population data [21,39] since otherwise there may be many vector fields that explain the observed marginals. These inference approaches are therefore structurally limited to the types of dynamics they can infer – they cannot, for instance, model populations of cycling cells.

The population dynamics *p*_*t*_ that arise from (1) are described by a Fokker-Planck equation

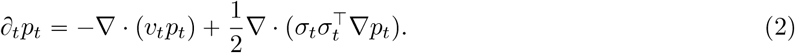

Let 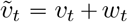 where *w*_*t*_ can be taken as any other vector field such that ∇· *w*_*t*_ = 0 and *w*_*t*_ · ∇*p*_*t*_ = 0. Then,

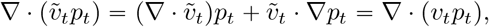

i.e. 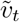 generates the exact same populations as *v*_*t*_. As a simple example, if *p*_*t*_ is an isotropic Gaussian centered at the origin for all *t >*0, then one may take 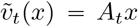 where *A* can be *any* antisymmetric matrix.

Pseudotemporal orderings are a subset of trajectory inference methods which aim to assign an ordering of cells in an observed population. This contrasts with methods like PBA and StationaryOT which aim to directly construct the Markov process. Methods such as diffusion pseudotime [38] require manual assignment of an initial cell, while some other approaches such as CytoTRACE [97] and SCENT [98] use heuristics such as transcriptional diversity to extract directionality. While these approaches aim to impose a partial or total ordering on cells and may appear incompatible with our Markov process point of view, we point out a straightforward approach for constructing Markov chain dynamics, given an ordering. Writing *T(x*_*i*_) to be the pseudotime assigned to cell *x*_*i*_, take

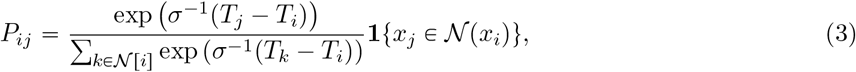

where *σ >*0 is a spread parameter, 𝒩 (*x*_*i*_)denotes the neighbourhood of cell *x*_*i*_ and **1** is an indicator function. In this way, the dynamics specified by *P* are adapted to the empirical cell-state manifold 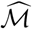 while still encoding the directionality of pseudotime estimates. Finally we remark that, different from the above methods which rely only on gene expression data, there are a variety of emerging trajectory inference approaches which use additional information to extract directionality. These include lineage tracing and phylogenetic information [89, 92], B-cell receptor dynamics [93], and multi-omics [99] information. The construction of *P* in this setting we find to be highly method-specific.

#### Undirected inference without dynamical information

Dynamical information is not always straight-forward to obtain for a given dataset. Metabolic labelling data are experimentally non-trivial to obtain, quality of RNA velocity outputs can be strongly dependent on dataset and modelling assumptions, and trajectory inference methods can be sensitive to parameter and preprocessing choices. Similarly, some snapshot measurements are made from the equilibrium distribution of the underlying biological process, at which point temporal resolution is lost. In these settings, inference of causal interactions between genes is not possible without additional (e.g. perturbation response) data or structural assumptions on a mechanistic model [100]. Instead, the goal of inferring undirected interactions in this setting has been addressed by a range of methods, of which GENIE3 [46] and PIDC [28] have been among the most successful [6]. The locaTE framework can be seamlessly applied to infer undirected cell-specific networks as a special case. In the absence of inferred dynamics, from an observed cell population, 𝒳, a Markov chain can be constructed as an undirected random walk on the cell state manifold. There are a variety of ways in which this random walk can be constructed: for example, a random walker located at cell *x* randomly chooses another cell *x*^′^ in its neighbourhood, 𝒩 (*x*), with a probability proportional to exp(− ‖ *x − x*′ ‖^2^ /*σ*^2^). Under an appropriate normalisation of the transition kernel [36, Section 3], the resulting random walk is consistent with the stochastic dynamics of a system at equilibrium where the vector field *f* is

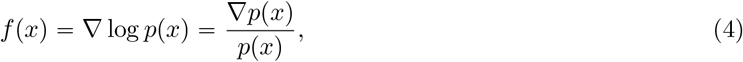

where *p(x*) is the population density of cells. In this setting, the potential − log *p (x*) is consistent with the observed data being at *equilibrium*. Since detailed balance holds at equilibrium, the dynamics are invariant under time reversal and therefore one can only hope to recover *undirected* interactions between variables.

### 4.2 Transfer entropy

#### Information measures

Information theoretic measures quantify dependences among random variables and do not depend upon restrictive modelling or parametric assumptions [101]. As introduced in brief earlier in Box 1, the central quantity of interest in information theory is the *entropy* of a random variable. Given a random variable *X*, the entropy of *X* can be written as

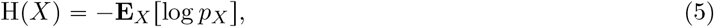

where *p*_*X*_ is the probability density of *X* relative to the Lebesgue measure (for the case of continuous random variables) or to counting measure (for the case of discrete random variables). H (*X*) measures, in *nats* (or bits, if the logarithm is taken base 2), the amount of *uncertainty* present in *X*. Given another variable *Y*, the mutual information between *X* and *Y* measures the degree of dependence. This is defined as the *reduction* in uncertainty in *Y* obtained by conditioning on *X*:

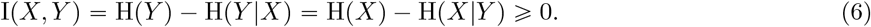

The mutual information provides a sufficient and necessary condition for independence of random variables:

*I*(*X, Y*) = 0 if and only if *X, Y* are independent. In the above, H(*X*|*Y*) is the *conditional* entropy:

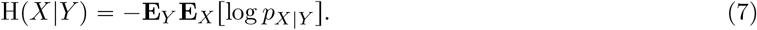

In other words (and abusing notation), writing *f (y)* = H (*X* |*Y =y*), we have that H (*X* |*Y)* = **E**_*Y*_ *f(y*). An alternative route to understanding the mutual information is that it is the relative entropy, or Kullback-Leibler divergence, between the joint distribution *p*_*XY*_ and the product of marginals *p*_*X*_ ⊗ *p*_*Y*_:

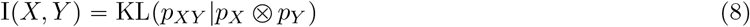

Viewed in this way, I (*X, Y*) is an explicit information-theoretic measurement of the deviation of (*X, Y*) away from independence. For a third random variable *Z*, the *conditional* mutual information I (*X, Y*| *Z*) can be defined as

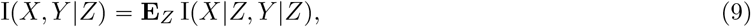

where (*X*|*Z* = *z*), (*Y* |*Z* = *z*) are random variables (*X, Y*) conditioned on {*Z* = *z*}.

#### Granger causality and transfer entropy

The problem of recovering dependency structure in dynamical systems is a classical problem in time series analysis: given trajectories *X (t) = [X*^(1)^ (*t*), …, *X*^(*d*)^ (*t)]* sampled from a *d*-dimensional stochastic dynamical system, the goal is to detect *causal* relationships between the variables. Here, temporal resolution allows us to apply a physical causality principle, namely that *causes* must precede their *effects*. The problem of network inference then amounts to pairwise testing of causal influence between pairs 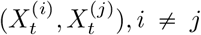. In what follows, we write (*X*_*t*_, *Y*_*t*_) to denote such a pair of stochastic processes.

An established approach to measuring causal influence is that of *Granger causality*, due to Norbert Wiener and Clive Granger [102]. In this framework, *X*_*t*_ is said to “Granger-cause” of *Y*_*t*_ if past knowledge of *X*_*t*_ “improves” forecasting *Y*_*t*_. In what follows, we provide a brief overview of Granger causality following Bressler and Seth [103]. In discrete time, define

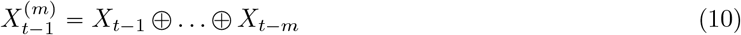

to be the *m* observed time-lagged states of *X* from times *t −m* up to *t −*1. One may then consider the following two linear autoregressive models for *X*_*t*_, one using only the history of *X*, and the other using the joint histories of (*X, Y*):

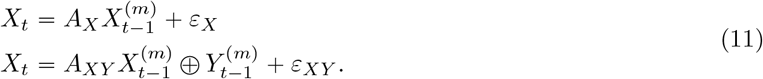

The insight of Wiener and Granger was that if there exists a *causal* (broadly interpreted as *time lagged*) relationship *Y* → *X*, then knowledge of the *m*-history of 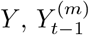, would improve the prediction of *X*_*t*_. In the setting of (11) this corresponds in a reduction of the variance of the residual term **Var**(*ε*_*X*_) < **Var**(*ε*_*X*_). The Granger causality score is defined by [104] to be

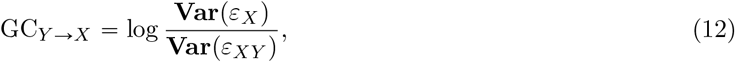

and a large reduction factor in in the residual variance implies a high Granger causality score. While Granger causality as a statistical measure of dependence is distinct from the physical notion of causality [105] it still has proven a useful tool in practice. In particular, it has been widely adopted to quantify temporal dependencies in neuroscience and finance. The linear nature of (11) is both a strength and a pitfall of the Granger causality framework: while it allows for straightforward estimation of Granger causality scores in practice, it assumes that the underlying dynamical process is linear and Gaussian.

Transfer entropy is the information-theoretic generalisation of Granger causality, aimed at quantifying temporal, directional *flow* of information between two time series. For a memory length *m*, the transfer entropy *Y* → *X* is

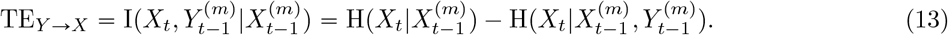

Similarly to the Granger causality score, TE_*Y* →*X*_ measures the reduction in uncertainty, as measured by the entropy, in *X*_*t*_ due to measurement of 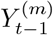, conditional on knowledge of its own history, i.e. 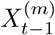. Notably, Barnett, Barrett and Seth [29] showed that the Granger causality score (11) is exactly the transfer entropy (13) in the case where all variables are Gaussian. This makes the link between the two quantities concrete. As an information theoretic measure, transfer entropy (13) does have assumptions on the distribution of (*X, Y*). This makes it ideal for applications to real-world, nonlinear complex systems.

#### Local transfer entropy for expression dynamics

We represent the transcriptional state of a cell as a vector of expression levels *X = (X*^(1)^, …, *X*^(*d*)^. Transfer entropy does not make assumptions on the specific form of a model, but for the sake of illustration let us consider a model of cellular dynamics where the expression level of each transcript *X*^p*i*q^ is governed by a stochastic differential equation (SDE) of the form

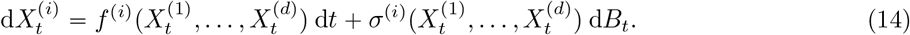

Note that this framework includes the chemical Langevin equation (CLE) [106], a biophysically derived model of single cell expression dynamics. Under the Itô interpretation [96, 107], for a small time interval *τ*, we may write approximately,

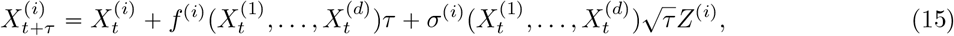

where the *Z*^(*k*)^ are independently and identically distributed Gaussian random variables with zero mean and unit variance. We can condition on 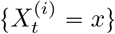.

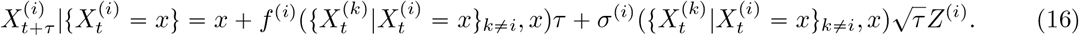

For any other gene *X*^(*j*)^, *j ≠ i*, the transfer entropy (13) with time lag *τ* is

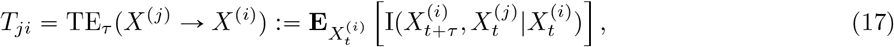

Under this discrete-time approximation, the transfer entropy *T*_*ji*_ will be positive whenever *f* ^(*i*)^ or *σ*^(*i*)^ have a functional dependance on *X*^(*j*)^, since we then have 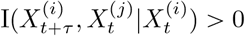.

Simply having a positive transfer entropy does not imply the existence of direct dependencies due to *indirect* relationships that may arise – for a third gene *k*, one in general will have *T*_*ki*_ *>*0 in the absence of a direct dependence of *X*^(*i*)^ on *X*^(*k*)^. As an illustrative example, consider a setting where for all *i, j, k* distinct, 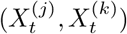 are non-independently distributed and 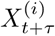 is a function of 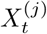 but not 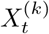, as would be the case in (16). Then 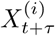 is independent of 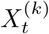, conditionally on 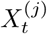. That is,

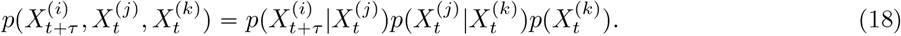

Whenever 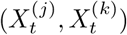 are not independent, one has *T*_*ki*_ > 0 since information about 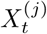 is also present in 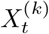, even though there is no direct dependency. Fortunately, the data processing inequality allows us to conclude that the transfer entropy for the direct interaction *X*^(*j*)^ → *X*^(*i*)^ will always dominate:

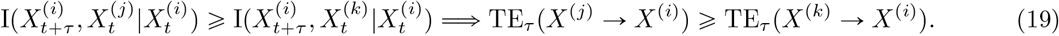

Although the data processing inequality gives us some assurances on detection of direct interactions, indirect interactions will still result in non-zero transfer entropies which is not ideal. This issue can be resolved by conditioning: define a *conditional* version of the transfer entropy, also known as *causation entropy* [108]:

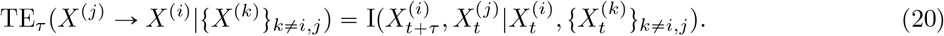

The conditional transfer entropy (20) measures the information conveyed to 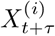 by 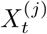, accounting for all possible confounding factors as well as the history 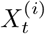. Revisiting the scenario (18), we have immediately that

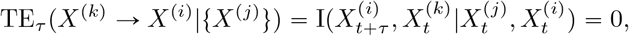

from conditional independence of 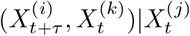 due to the Markov chain assumption (18).

The conditioned transfer entropy (20) accounts for the value of all confounder variables simultaneously, and can thus be re-written

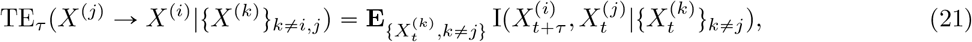

where we abuse notation to write as a random variable 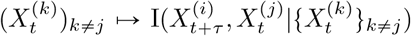. While (20) is theoretically sound and would ensure a score of zero for any indirect interactions, its computation is intractable whenever the number of variables to condition is even mildly large due to the number of data points required and computational cost [15]. Motivated by both this observation and the application to context-specific network inference in single cell datasets, we propose an *alternative* conditioning strategy that retains the ability to condition out confounders, while retaining a level of granularity. For any cell state *x*, let 𝒩(*x*) be a set with positive volume denoting its neighbourhood. We can define the *neighbourhood-conditioned* transfer entropy:

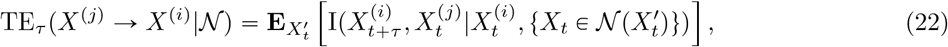

Instead of conditioning on a event fixing the *values* of all other variables 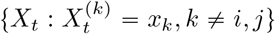, we now condition on an event that specifies a *neighbourhood* of the state space {*X*_*t*_ : *X*_*t*_ ∈ 𝒩(*x*)}. Equivalently, this can be understood in terms of a *nonlinear function* of all variables that parameterises the neighbourhood:

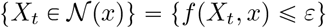

for some function *f* and neighbourhood size *ε*. In practice, the neighbourhood 𝒩 arises from some (potentially non-linear) function of the overall gene expression profile that is determined by the construction of the cell state manifold (e.g. in the case of PCA, this would be an affine combination of all genes). Writing *X*^−(*i,j*)^ to denote the vector *X* excluding the (*i, j*) th entries, we remark that for the choice 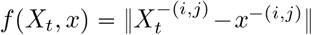 we have that as *ε* → 0,

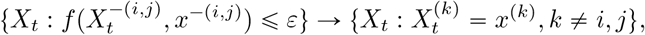

and we recover the conditional transfer entropy (20). Although transfer entropy has previously been used for inference of static networks from pseudotemporally ordered expression data [15, 17], existing methods calculate TE using lagged tuples of cells along a total ordering. Not only does this introduce a strong dependence upon the provided pseudotemporal ordering; it results in loss of resolution since conditioning (if any) is done across genes, not cells. This results in all interaction scores being population averages – any variation in regulatory activity over the expression space is integrated out to yield a static network. This can be problematic since biological trajectories may exhibit bifurcating, cyclic or converging trajectories, potentially requiring multiple runs of GRN inference on different subsets of a dataset.

Although neighbourhood-conditioned transfer entropy as written in (22) involves an expectation taken over all neighbourhoods, it can be decomposed as a sum of neighbourhood-wise contributions:

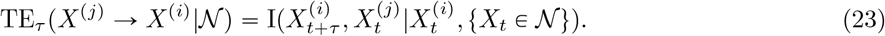

We call TE_*τ*_ (*X*^(*j*)^ − *X*^(*i*)^| 𝒩) the *local* transfer entropy for neighbourhood 𝒩. Intuitively, this can be understood as the transfer entropy (17) for trajectories started in the neighbourhood 𝒩, i.e. {*X*_*t*+*τ*_ : *X*_*t*_ ∈ 𝒩}.

#### Limitations of transfer entropy for network reconstruction

Limitations of the transfer entropy and related causal inference approaches have been the subject of protracted debate in the complex systems literature. We briefly discuss several fundamental assumptions and limitations in what follows: the first two of these limitations can be ameliorated by development of more sophisticated inference methods, while the third is fundamentally due to experimental measurement limitations.

- **Non-localisable or higher-order interactions** James, Barnett and Crutchfield [109] demonstrate that the transfer entropy is susceptible to overestimation of causal influence in cases where the true flow of information is not simply from one variable to another, but where multiple random variables *synergistically* drive another. The authors demonstrate that the *unconditional* transfer entropy can fail to detect dependencies where higher order interactions are present, while the *conditional* transfer entropy is successfully detects but overestimates its value. The authors ascribe this pitfall to the the ubiquitous use of pairwise interactions as a model of dependencies in complex systems.
- **Entropy estimation** The cost of adopting model-free information measures is that the entropy of a continuous random variable is not straightforward to estimate from samples. This has given rise to a wide range of estimators, and entropy estimation remains an open research problem – we refer the reader e.g. to [110] for a recent overview. Our application (network inference) requires fast computation of conditional mutual information for many triplets of variables, and so we use the discretisation approach of [28].
- **Hidden variables and undersampling of dynamics** Smirnov [111] identifies partial observations and undersampling of dynamics as two scenarios that can lead to spurious transfer entropies. More generally, hidden variables can result in delays in observed dynamics.

## 5 Methods

### Inferring cell-specific networks

In what follows, we write ⟦ *n* ⟧ = {1, …, *n*} for *n* ∈ ℕ. For each cell *x* ∈ 𝒳 we construct (using e.g. a uniform distribution, truncated Gaussian, etc.) a probability distribution 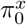 supported on the neighbourhood 𝒩 (*x*0. We then consider a Markov process 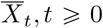 that starts from 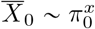 and evolves under *P*. For some timestep *τ* > 0 the resulting *coupling* is

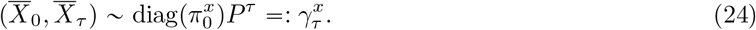

For a pair of genes, (*j, k*), we quantify the *local transfer entropy* in the neighbourhood 𝒩 (*x*) for the relationship *j* → *k* from knowledge of the coupling, 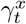, and the gene expression states:

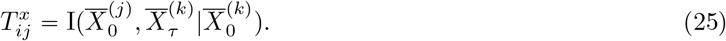

Repeating this computation for all cells, *x*_*i*_ *∈ 𝒳*, and gene pairs, *j, k* ∈ ⟦ *d* ⟧, we arrive at a tensor, *Ĝ* ∈ R^*N* × *d* × *d*^, such that 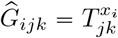, the local transfer entropy score of *j → k* in the neighbourhood of cell, *x*_*i*_.

The coupling defined in (24) is forward in time, between (0, *τ*). Resulting TE scores may reflect the underlying dynamics up to a shift in time. To remedy this, we consider couplings that are *symmetric* in time, between (−*τ, τ*). While time-reversal of a diffusion-drift process away from equilibrium is generally not a well-posed problem, in practice, suitable backward kernel have been constructed by transposing the transition matrix [23, 44]. Given such an approximate backward kernel, *Q*, a time-symmetric coupling can be constructed:

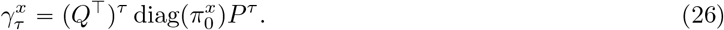

#### Constructing a backward kernel

It is straightforward to write:

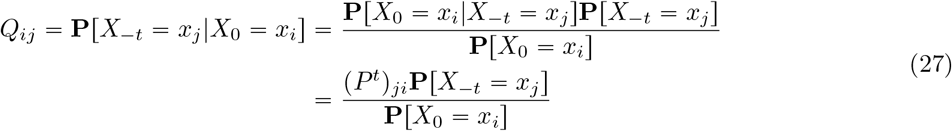

If *P* encodes a reversible Markov chain and we chose **P [***X* _*−t*_ *= ·]*to be the stationary distribution, we would have time-reversal at equilibrium. Since we are interested in the behaviour of the process away from equilibrium, we must prescribe **P [***X* _*−t*_ *= ·]*away from equilibrium. In practice, we find that taking **P** *X* _*t*_ Unif works well – this is equivalent to simply taking the transpose of *P* and rescaling, as done in [23]. The construction of this backward kernel is discussed in detail in [44, Section 4.3], where it is shown that this is consistent with negating the vector field *v* ↦ −*v* in the continuous limit.

#### Filtering and denoising

Since the TE score matrix, *Ĝ*_*i*.._, for each cell *x*_*i*_ has to be learned from its local neighbourhood 𝒩(*x*_*i*_), it is potentially noisy. Furthermore, it is impractical (due to limited data and computational resources) to condition on all potentially indirectly interacting genes [15]. Thus, TE scores may contain noise and spurious signals resulting from observational noise or indirect interactions. To deal with this, we first filter interaction scores using context likelihood of relatedness (CLR) algorithm [28, 112] and then solve a manifold-regularised optimisation problem to de-noise interaction signal [11, 15].

#### Context likelihood of relatedness

For raw TE scores, *Ĝ*, the data processing inequality (DPI) (19) gives us a basis for filtering for indirect interactions. While methods such as ARACNe [113] and TENET [17] apply the DPI by iterating through triplets (*X, Y, Z*) and filtering potential indirect edges, we use the context likelihood of relatedness [45] algorithm which was found to have better performance [45] and was also used in [28]. From a performance point of view, direct application of the DPI by iterating through triplets is *O* (*d*^3^) in the number of genes *d*, while the CLR algorithm is *O* (*d*^2^). Since we focus on context-specific network inference, we potentially need to filter many networks rather than a single static network. Thus we opt for CLR, although we mention that the direct application of DPI can be used.

For a pair of genes, (*i, j*), and given a interaction score matrix, *A*, we compute *z*_*i*_ (resp. *z*_*j*_) to be the *z*-score of *A*_*ij*_ with respect to *A*_*i*_. (resp. *A*_.*j*_). Then we define the *weighted* CLR score for (*i, j*) to be

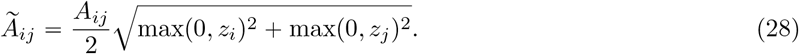

Applying CLR filtering along the first axis of *Ĝ*, we obtain the filtered tensor,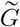 ;. We modify the original approach of [45] and weigh the CLR score by the initial MI value *A*_*ij*_. This is important since CLR was originally designed to filter interactions in *static* networks. In cell-specific networks, very few edges may be “active” in any given cellular context; then entire rows or columns of *A* may consist only of noise. Computing *z*-scores along those rows or columns would put both noise and signal on the same scale.

#### Denoising via manifold-regularised regression

The tensor 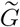 contains a (noisy) matrix of filtered interaction scores, one for each cell. For notational convenience we write 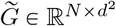, i.e. each row is a length-*d*^2^ flattened TE score matrix. We propose to solve the optimisation problem, 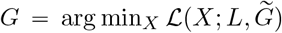, where

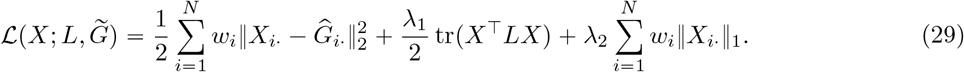

Here *λ*_1_, *λ*_2_ are hyper-parameters, *w* is a vector of cell weights, and *L* is the symmetric graph Laplacian on ℳ. The term associated with *λ*_1_ is a manifold regularisation term corresponding to an expectation that regulatory relationships should vary smoothly with changes in cell state, i.e. tr (*X* ^⊤^*LX*) is large for rapidly fluctuating *X*. The term associated to *λ*_2_ is a Lasso [2] term that encourages *X* to be sparse, reflecting our knowledge of biological networks [11, 16]. Together, the objective (29) encourages both parsimony and sharing of information along the cell state manifold. The problem is a case of L1-L2-regularised least squares, and can be solved efficiently using an iterative scheme.

The problem (29) is convex and a numerical algorithm for its solution can be derived using the alternating direction method of multipliers (ADMM) [114]. With auxiliary variables *W, Z* with the same dimensions as *X*, the corresponding ADMM scheme is

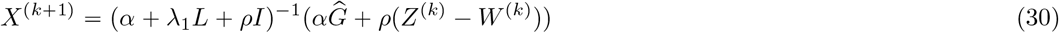

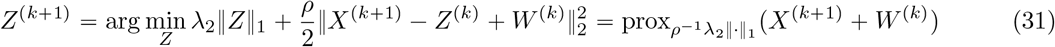

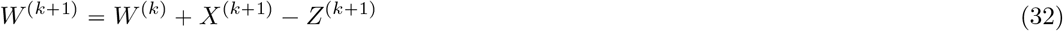

where *ρ* > 0 is the ADMM relaxation parameter (we take this to be 0.05), *α* = diag(*w*) and

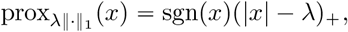

interpreted elementwise.

#### Uncovering regulatory modules via manifold-regularised NMF

Now we consider the same problem setting as (29), but this time we require that the *N* × *d*^2^ matrix of recovered interactions *G* be decomposed into two non-negative factors [115]. For a target rank *k*, we seek “tall” matrices 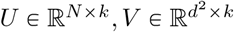 such that *G≈ UV* ^⊺^. That is, for a cell *i*, its network is given by a linear combination of the columns of *V* with coefficients *U*_*i*1_, …, *U*_*ik*_:

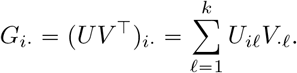

We consider a general loss function of the form

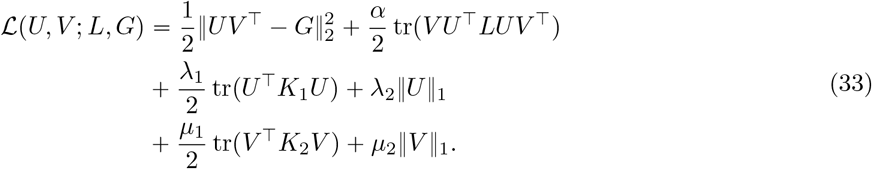

We explain the loss function term by term. The first term encourages a fit to the matrix *G* of raw TE values. As in the case of the denoising regresion, we impose a penalty on the smoothness of the low-rank reconstruction (w.r.t. the graph Laplacian *L*) with weight *α*. In addition, we can impose Tikhonov and sparsity penalties on the coefficients and atoms.

One can derive multiplicative update equations for the gradient descent, following the approach of [115]. For positivity-preserving updates, we decompose any Laplacian matrices into their positive and negative parts:

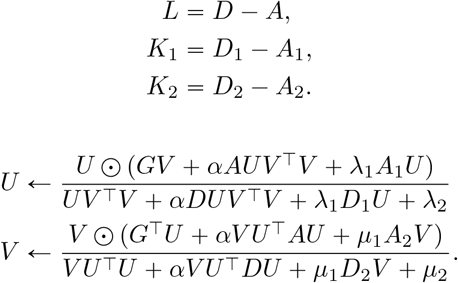

#### Normalisation of local transfer entropy scores

For information theoretic measures such as mutual information normalisation is not straightforward [28, 45], and therefore direct comparison of edge scores may not be an accurate reflection of relative edge confidence. We employ a cumulative distribution function (CDF) normalisation procedure to locaTE scores prior to downstream analysis which was found to work well in [28]. For a *d* × *d* reference matrix *A*, Gamma distributions are fit to each row and column to yield CDFs 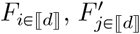. A CDF-transformed matrix 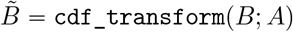 is then constructed as

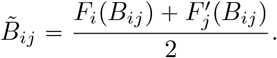

LocaTE produces a *N ×d ×d* tensor *G* which describes the smoothed *d ×d* cell-specific network in each of the *N* input cells. We construct the reference matrix as *A*_*jk*_ = max_*i*_ *G*_*ijk*_, where the maximum is taken since interactions may only be non-zero in a small subset of cells. The CDF-transformed tensor 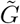 is thus constructed cell-wise as

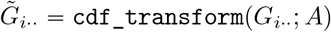

#### Computational complexity

The problem of cell-specific network inference requires 𝒪 (*Nd*^2^) transfer entropy calculations for *N* cells and *d* genes, since each cell-gene-gene triplet must be considered. For each triplet, (*i, j, k*), locaTE calculates the joint distributions of 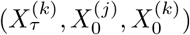 under the coupling *γ*^(*i*)^. While in the worst-case (if *γ* were dense) this would have complexity 𝒪 (*N* ^2^), in all our applications *γ* is sparse and transitions occur only between nearby cells; the complexity is therefore 𝒪 (|*γ*^(*i*)^|) ≪ *N* ^2^. The CLR filtering step is 𝒪 (*Nd*^2^), and the regression step primarily involves iterative matrix products of sparse *N* ×*N* against *N ×d*^2^. We implemented locaTE using the Julia programming language [116] to take advantage of its speed for numerical computation.

With exception of the regression step, which is global, all computational steps are easily parallelised across cells. The calculation of TE scores can alternatively be parallelised across gene pairs. This allows locaTE to take advantage of large scale parallelisation when available, potentially drastically reducing the compute time required. We find empirically that a large portion of computational burden arises from the construction of the joint distribution, 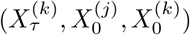. Since this is an accumulation operation, this can be accelerated using GPU computation. Other steps in the locaTE pipeline (regression, factor analysis) amount to linear algebra operations and are amenable to GPU acceleration. For datasets in excess of ∼10^4^ cells, the computational cost could be greatly reduced by first grouping sampled cells into metacells, using e.g. the approach in [117]. We discuss this in *Coarse graining for large datasets* and present an example in Figure S8.

#### A heuristic for selecting *τ*

The time-step *τ* is the key parameter underlying locaTE, as it is analogous to the delay parameter central to Granger causality and transfer entropy. In the case of locaTE, the choice of *τ* is dependent on the construction of the transition matrix *P* – taken together, (*P, τ*) determines the “timescale” along which transfer entropy scores will be calculated. To see this, note that *P* by itself has an intrinsic timescale – this could be affected e.g. by the number of nearest neighbours used to calculate transition probabilities using RNA velocity/metabolic labelling methods or PBA [21, 23, 24, 37]. In such a scenario, a large number of neighbours would increase the range of transitions a cell can take in a single step and therefore corresponds to a longer timescale. Similarly, a small number of neighbours would correspond to a shorter timescale.

Since locaTE can be applied with *any* choice of underlying dynamical inference method, giving general guidelines for selecting *τ* is difficult. Furthermore, quantifying the “timescale” encoded by a transition matrix *P* is also not straightforward. For a cellular population, we provide a heuristic method for identifying a range of reasonable values for *τ*, given knowledge of the scale of meaningful biological variation in the form of a coarse clustering of cells.

Let *P* be a given transition matrix supported on a discrete set of observed cells, 𝒳 = {*x*_1_, …, *x*_|*X* |_}. Furthermore, let 𝒞_1_ ⊔ … ⊔ 𝒞_*k*_ be a *partitioning* of X into *k* clusters (in practice, this can be constructed using any clustering method). Fix one cluster 𝒞_*i*_. Then, for any other cell state *x*_*j*_ ∉ 𝒞_*i*_ one may then define the *mean first passage time* for a random walk started at *x*_*j*_ to hit 𝒞_*i*_:

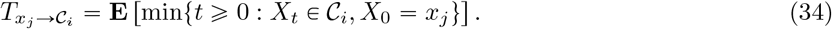

Clearly, 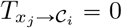 if *x*_*j*_ *∈ 𝒞*_*i*_ and 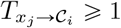 otherwise. Then, using a well-known formula for first passage times in discrete-time Markov chains [118]:

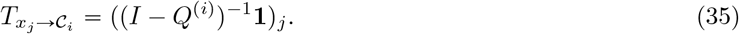

In the above, *Q*^(*i*)^ is the transient part of the fundamental matrix of the absorbing Markov chain obtained from *P* after modifying all states in 𝒞_*i*_ to be absorbing. That is, modify *P* by setting rows corresponding to *C*^*(i*)^ equal to rows of the identity *I*, forming *P*^*(i*)^. Then, up to reordering rows and columns, *P*^*(i*)^ can be written as

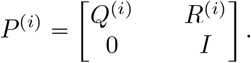

In practice, solution of the linear system (35) may be ill-conditioned, so we invert instead (1 + *ϵ*)*I* − *Q*^(*i*)^ where *ϵ* is taken to be a small positive constant, e.g. *ϵ* = 10^−3^ to ensure numerical stability.

Once we have calculated 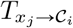 for all *x*_*j*_ ∉ 𝒞_*i*_, we set

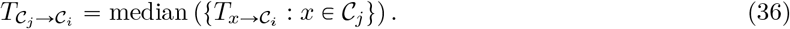

This gives a measure of the *typical* time for a transition 𝒞_*j*_ *→𝒞*_*i*_ between two partitions, in terms of steps of the Markov chain specified by *P*. This gives a heuristic for the time-step *τ*:

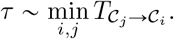

That is, to choose the time-step *τ* to be comparable to the shortest typical transition time between a pair of clusters. In Figure S4 we illustrate this heuristic for the bifurcating and switch examples from Section 2.2. Notably, in both cases the heuristic suggests *τ* ≈ 3, which is the value used to produce the results in Section 2.2.

#### Limitations and caveats

We now list and discuss several anticipated failure modes and caveats of locaTE which may be instructive.

- *Lack of continuous variation in time-series data* LocaTE relies upon a Markov chain approximation of cell dynamics and a manifold assumption on the single cell phenotypical space. These assumptions may be violated, for instance, in time-series settings where there are large “gaps” between timepoints. In this setting, we expect that Markov modelling approaches will fail in general, since they lack the ability to interpolate between observed populations. Appropriate modelling may involve the use of more sophisticated approaches, such as ODE or SDE modelling, or snapshot observations where only mature cell states are observed in disjoint clusters.
- *Lack of heterogeneity in tightly clustered data* Relatedly, in settings where distinct celltypes are grouped into tight, disjoint clusters, we expect that limited information can be obtained from context-specific inference approaches due to the lack of heterogeneity. Context-specific inference approaches are analogous to conditioning on cluster, and therefore may miss global dependencies that emerge across clusters. In settings with strong cluster structure, therefore, a population-level, static network inference approach may be more appropriate.
- *Errors in dynamical inference* Dynamical inference methods may suffer from inaccuracies and errors. Notably, RNA velocity approaches rely on kinetic models which may be invalid in certain systems, leading to random flows or even backflow [40]. In the context of locaTE, backflows in the dynamics inference could flip the direction of causal influence in inferred networks. We therefore advise quality and sanity checks of the inferred dynamics that are provided as input to locaTE using biological prior knowledge, using common approaches such as visualisation of streamplots [37] or fate probabilities [21,23]. In scenarios where no prior information is available on dynamics and high-confidence dynamics estimates are unavailable, we recommend use of the undirected kernel to infer undirected networks.
- *Misspecification of parameters* In addition to cell-gene expression data and inferred dynamics, locaTE depends on a number of parameters which are involved in the computation of transfer entropy scores and subsequent processing. In Table 1 we list parameters and provide guidelines for setting them. Most importantly, the neighbourhood bandwidth *ε* and time-lag *τ* affect the spatial and temporal scales across which networks are inferred. Choosing these scales too large risks losing resolution and inaccuracy in distinguishing between direct and indirect interactions.

**Table 1:** Key parameters and recommendations for choosing their values.

| Parameter | Required for | Type | Effect and default/recommended values |
| --- | --- | --- | --- |
| $\varepsilon$ | TE calculation | Positive real | **Bandwidth parameter for neighbourhood construction. Default:** $\varepsilon = 5\bar{C}$, where $\bar{C}$ is the average squared distance between pairs of cells. Note that we construct neighbourhoods using the approach [119] where the bandwidth is determined implicitly; however other constructions could be used (e.g. uniform on neighbourhood, truncated Gaussian). Choosing $\varepsilon$ too small (smaller than the scale of meaningful biological variation) may result in causal relationships being missed. Choosing $\varepsilon$ too large may result in estimation of indirect relationships. |
| $\tau$ | TE calculation | Positive integer | **Time-lag in terms of the transition matrix $P$. Default:** $\tau = 1$. Choice of $\tau$ depends on the construction of $P$: choosing $\tau$ too small (smaller than the scale of meaningful biological dynamics) may result in causal relationships being missed. Choosing $\tau$ too large may result in estimation of indirect relationships. Also see: *A heuristic for selecting $\tau$*. |
| $\lambda_1, \lambda_2$ | Denoising regression, NMF | Positive real | **Laplacian regularisation and LASSO parameters for smooth and sparse regression. Default:** $\lambda_1 = 25, \lambda_2 = 10^{-3}$. Choosing $\lambda_1$ too large may reduce resolution of inferred context-specific interactions, and choosing $\lambda_1$ too small may lead to noisy results. Choosing $\lambda_2$ determines the sparsity level in the regression result: too large a value may result in many zero scores, while too small a value may not reach the desired sparsity in the result. |
| $k$ | NMF | Positive integer | **Number of regulatory programs to learn** Choosing $k$ too large may result in spurious programs that only capture noise, while choosing $k$ too small may result in meaningful variation in inferred networks being missed. |

#### Simulation of bifurcating and switch systems (Section 2.2)

We simulate trajectories from each system using the BoolODE software package [6]. This uses a chemical Langevin equation scheme for simulating biophysically plausible expression dynamics: d*X*_*t*_ = *f*(*X*_*t*_)d*t* + *g*(*X*_*t*_)d*B*_*t*_ where *f, g* have specific forms dictated by the reaction network (see [6] and BoolODE documentation for more details). From independent realisations of the system’s trajectories, we generate a set of 1,000 sampled cells by sampling a cell state from each trajectory at a time chosen uniformly at random. As a measure of ground truth interactions for each sampled cell *x*_*i*_ we compute the corresponding Jacobians *J*_*ijk*_ *∂*_*j*_*f*_*k*_(*x*_*i*_). The Boolean rules used to implement each system are described in Appendix S1.1.

#### Inferring dynamics for BoolODE simulated data

For trajectories simulated using BoolODE (including the benchmark cases of Section 2.3), we construct the Markov transition matrix *P* using a number of different approaches. In each case, dimensionality reduction was performed using PCA and a *k*-NN constructed with *k* = 25. Diffusion pseudotime was then assigned with the root cell taken to be the sampled cell with the earliest ground-truth simulation time. Various transition kernels *P* were then constructed as described below.

- **Velocity kernel** For each observed cell, *x*_*i*_, we calculate the corresponding velocity vector, *v*_*i*_ = *f*(*x*_*i*_). From this, we can construct transition probabilities to a neighbourhood of *x*_*i*_ (in gene expression space) using a kernel function *K*:

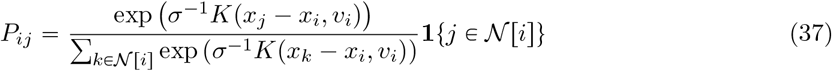

where **1** denotes the 0-1 indicator function. We selected the bandwidth *σ* following the median heuristic of [23]. Different choices of the function *K* can be used which lead to different theoretical interpretations as well as practical performance – we refer the interested reader to Section 4 for a more detailed discussion. We used the dot-product, cosine and correlation kernels.
- **Optimal transport kernel [39]** Cells in the top 92.5%-97.5% of pseudotime were assigned a negative flux rate so that the net negative flux was −50. Remaining cells were assigned a positive flux rate to satisfy the zero net flux requirement, ∑_*i*_ *R*_*i*_ = 0. StationaryOT [39] with quadratically regularised optimal transport was applied with Δ*t* = 1.0, *ε* = 0.05. The cost matrix was taken to be the matrix of pairwise graph distances constructed from the *k*-NN graph. To prevent short-circuiting resulting from spurious edges, edges were ranked by their random-walk betweenness centrality and the top 5% of edges were pruned.
- **Population balance analysis (PBA) kernel [21]** Population balance analysis [21] was applied using the previously constructed *k*-NN graph, the same flux rates *R*_*i*_ as used in the optimal transport kernel, and diffusivity *D* = 1.0.
- **Diffusion pseudotime kernel** The diffusion pseudotime (DPT) kernel was constructed from the previously estimated diffusion pseudotime ordering using the formula (3).
- **Undirected kernel** An undirected kernel was constructed by taking *P* to be the row-normalised neighbourhood kernel.

#### Inferring cell-specific networks (BoolODE simulated data)

LocaTE was applied using the simulated expression data and inferred transition kernels *P*. Expression values were discretised using the Bayesian blocks algorithm as implemented in [28]. The chemical Langevin simulation is a continuous approximation to biophysically realistic discrete expression dynamics, and as such produces expression values near zero. Simulated expression values < 10^−0.5^ were set to zero to prevent many bins from being created near zero. We remark that this is a simulation artifact specific to the BoolODE simulator and that this correction is not necessary for dyngen nor real data, which derive from counts. A neighbourhood kernel was constructed using quadratically regularised optimal transport [119], and the backward kernel was constructed from *P* following (27). For the examples shown in Figure 3, LocaTE was applied using *τ* = 3, *λ*_1_ = 25.0, *λ*_2_ = 10^−3^ for both systems.

#### Parameter sensitivity of locate

An exhaustive parameter sweep on (*τ, λ*_1_, *λ*_2_) was carried out (Figure S3), with

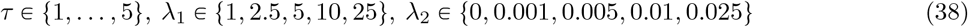

We find that the accuracy of locaTE depends mostly on the time step, *τ*, and the strength of the Laplacian regularisation, *λ*_1_. The choice of time step will depend on the nature of the input transition matrix: *τ* should be chosen large enough so that application of *P* ^*τ*^ captures changes in the cellular state, but not so large as to lose information of the cell-cell coupling between times *X* _−*τ*_, *X*_*τ*_. Similarly, *λ*_1_ should be chosen large enough to have a denoising effect, but not so large as to wash out signal. We find that *λ*_2_ has less of an effect on performance as measured by AUPRC and very little effect on the EP. We find that in both simulated datasets the AUPRC increases for increasing *λ*_1_ while the EP decreases for *λ*_1_ too large. This reflects a tradeoff to be made – at moderate levels, the smoothing amplifies signal and improves both the AUPRC (representing accuracy across all levels of recall) and the EP (representing accuracy at low recall, or high-confidence predictions). High levels of smoothing wash out high-confidence signal despite still increasing AUPRC due to improved performance at low confidence. Ideally therefore a balance should be struck between achieving good overall performance and good performance for high-confidence predictions.

#### Code availability

An open-source Julia implementation of locaTE is available at https://github.com/zsteve/locaTE.jl.

#### Benchmarking cell-specific and static network inference methods

To benchmark the performance of locaTE, we considered a number of other network inference methods. We considered three cell-specific network inference approaches: CeSpGRN, SpliceJAC and locCSN. Since locaTE can be used to infer static, population-level networks as well, we considered a number of static inference methods: PIDC, GENIE3, TENET, SCRIBE, SCODE, GRISLI, and SINCERITIES. We summarise the methods as follows.

- **CeSpGRN** infers cell-specific, signed, undirected networks from single cell expression data using a copula Gaussian graphical model.
- **SpliceJAC** infers cluster-specific, signed, directed networks from single cell spliced and unspliced expression data using a dynamical systems model.
- **locCSN** infers cluster-specific, unsigned, undirected networks from single cell expression data using pairwise independence tests.
- **PIDC** infers population-level undirected, unsigned networks from single cell expression data using partial information decomposition (PID). We chose to include PIDC due to its good performance in the benchmarking study of [6].
- **GENIE3** infers population-level undirected^1^, unsigned networks from single cell expression data using tree-based ensemble methods. We chose to include GENIE3 due to its good performance in the benchmarking study of [6].
- **TENET** and **SCRIBE** infer population-level directed, unsigned networks from single cell expression data together with pseudotime ordering. We chose to include TENET and SCRIBE since they, too, use the transfer entropy to infer directed networks.
- **SCODE, GRISLI** and **SINCERITIES** infer population-level, signed, directed networks from single cell expression data together with pseudotime ordering. We chose to include these methods as additional baselines for directed network inference.

CeSpGRN relies on several hyperparameters: neighbourhood size, kernel bandwidth, and sparsity level. In our benchmarks, we tried a range of 5 values for each of these three hyperparameters:

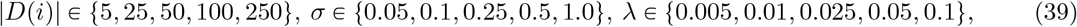

making a total of 125 parameter combinations for each test instance.

SpliceJAC and locCSN both require a cell cluster annotation to be provided: to generate this, we used the Louvain algorithm with resolution parameter set to 1.0. SpliceJAC may fail to handle all genes when cluster sizes are too small, and so if this occurred we reduced the resolution parameter by a factor of 2 and tried again.

SpliceJAC requires spliced and unspliced expression values. BoolODE does not model splicing but only mRNA and protein levels. We therefore used the protein and mRNA expression values as the “spliced” and “unspliced” data, as was done by the authors in the original publication [14].

PIDC and GENIE3 do not have hyperparameters that require significant tuning, and were found to have good performance with default options.

TENET requires a history length parameter *k*, for which we tried 1 ⩽ *k* ⩽ 8 and took the best performing value. The authors reported that they generally found *k* 1 to perform best [17]. SCRIBE requires several delays to be given, for which we used 5, 20, 40. For both SCRIBE and TENET, the true simulation time was provided in place of pseudotime.

SINCERITIES requires discrete timepoints. Following [6], we did this by binning the true simulation time into 10 timepoints.

For methods where we try multiple parameter settings, we report the best-case performance.

#### Benchmark test instance generation

Since we are interested in network inference for developmental systems with discernible trajectory, we consider BoolODE and dyngen, two simulation tools designed specifically for this task [6, 10].

We use BoolODE to simulate and sample expression profiles for TF-TF networks. For each cell, we record its true simulation time, velocity vector, and Jacobian matrix as a measure of the ground truth regulatory interactions. For each choice of trajectory structure, 10 test instances were generated each with 1,000 cells. Dynamics were inferred as previously done for bifurcating and switch systems.

Unlike BoolODE, dyngen models dynamics using a discrete model and produces spliced and unspliced counts by simulation using the Gillespie algorithm. Furthermore, dyngen allows for modelling of downstream TF-target interactions as well as TF-TF networks. For each dyngen trajectory we considered, 2,500 cells were sampled as a snapshot along with the true simulation time and spliced and unspliced counts. For dyngen instances, we construct the transition kernel *P* from the sampled spliced and unspliced expression values using dynamo [24].

#### Evaluation: cell-specific network inference

For benchmark instances generated using BoolODE, we record the Jacobian for each cell as a measure of the ground truth network activity. For each triple (*i, j, k*) with *i* ∈ ⟦*N*⟧ and *j, k* ∈ ⟦*d*⟧, we treat the problem of detecting an edge, *j* → *k*, in cell *x*_*i*_ as a binary classification problem with threshold *q*. To construct the set of true positives, we consider the matrix Π*J*, where Π is a neighbourhood transition matrix such that 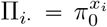. The motivation for considering this instead of simply using the raw Jacobians, *J*, is that cell-specific interactions are necessarily inferred using neighbourhood information. Strict cell-wise comparison to a ground truth would be overly stringent and sensitive to small perturbations in the expression space. By computing Π*J*, the ground truth signal is smoothed over the same neighbourhood that was used for inference, allowing for more robust assessment of classifiers. We classify all potential edges for which (Π*J*)_*ijk*_ > 0.5 to be “true” edges.

We consider unsigned, directed inference: therefore for methods that produce signed outputs we took the absolute value. Since some methods we compare to are only capable of inferring undirected networks, we provide an additional comparison where all inferred directed networks are first symmetrised, and then compared to the ground truth undirected network. This provides a fair comparison of locaTE to other approaches when considering only the undirected inference problem. Inferred networks were evaluated against the ground truth by constructing precision-recall (PR) curves and reporting the area under precision-recall curve (AUPRC), AUPRC ratio relative to a random predictor, and early precision (EP, which we define as the area under the PR curve up to 10% recall).

#### Evaluation: static network inference

Cell-specific networks inferred by locaTE, CeSpGRN, SpliceJAC and locCSN are averaged across sampled cells to produce population-level static networks. As in the cell-specific case, where signed networks are available we convert them to unsigned networks by taking an absolute value. Inferred networks for each test instance are then compared to the ground truth static network, and we report inference accuracy in terms of the AUPRC ratio. We also consider an undirected comparison as described previously in order to ensure a fair evaluation against undirected inference approaches.

#### Effect of vector field noise and dropout

We investigate in Figure S7 the effect of noise in the vector field estimates as well as dropout for the bifurcating network and switch network considered in Section 2.2. To model noise in the vector field *v*(*x*), we construct a noisy vector field 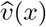 for a given noise level *η* ∈ [0, 1] componentwise:

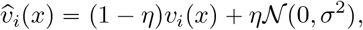

where *σ* is a scale factor to computed as the root-mean-square value of gene expression. We consider *η* 0, 0.1, 0.25, 0.5, 0.75. We implement dropout following the approach of [6, 28]. For a given dropout rate *q* ∈ [0, 1], for each gene *i* we select cells with the bottom 100 × *q*% expression and set the expression level to zero with probability *q*. We consider *q* = 0, 0.1, 0.25, 0.5. For each combination of (*η, q*), we apply locaTE with parameters (*τ, λ*_1_, *λ*_2_) = (3, 25, 0.001) using dot-product, cosine, diffusion pseudotime and undirected kernels. Summarising these results, we show the AUPRC ratio for static network inference over 10 sampled simulation datasets. We find that in the bifurcating network, dropout has relatively little influence on the performance of locaTE, while in the switch network we find that performance drops for higher dropout rates (*q* = 0.5) but is relatively stable for lower rates (*q* = 0.1, 0.25). Similarly, we find that performance of locaTE with dot-product and cosine kernels degrades at the highest noise level (*η* = 0.75), but is robust to lower noise levels. Curiously, we find that the cosine kernel suffers to a lesser extent than the dot-product kernel: this may be because the cosine kernel is only sensitive to the orientation of the vector field and is thus robust to the effect of noise on the magnitude of the vector field.

#### Coarse graining for large datasets

Interaction scoring by locaTE requires 𝒪(*Nd*^2^) TE scores to be calculated, where *N* is the number of cells or contexts to be considered and *d* is the number of genes. Each TE calculation involves construction of a joint probability distribution of expression levels over a cell state neighbourhood, whose size may depend on *N*. If the neighbourhood size scales linearly with *N* (and this is the setting that would be reasonable as *N* tends to infinity), the overall complexity of locaTE is 𝒪(*N*^2^*d*^2^). For datasets consisting of more than several thousand cells, this may result in significant computational load. A straightforward to scale this analysis to efficiently handle large numbers of cells is to employ a coarse graining of the cell population ahead of transfer entropy computation. This allows us to perform our analysis at a mesoscopic scale of *metacells*, instead of the microscopic scale of single cells [117,120]. We note that this is distinct from cluster-wise analysis since coarse graining typically produces still on the order of 10^2^ − 10^3^ metacells, compared to cluster analyses which typically produce ≪ 10^2^ clustered states [23].

LocaTE can handle coarse grained data by accepting a single cell dataset 𝒳 along with a partitioning {*C*_1_, …, *C*_*K*_} into *K* metacells, where each metacell is comprised of cells with sufficiently similar transcriptional states. We investigate this in terms of the switch-like system of Section 2.2 in Figure S8. While a variety of methods for constructing metacell partitionings already exist, to illustrate the efficacy of coarse graining we chose to construct metacells by a simple Leiden partitioning of the cell state graph. In terms of computational speed, we found that using coarse-grained cell states led to a vast speedup in computation times on the CPU: applying locaTE directly to the 10,000 cell states took approximately 2.1 hours, compared to just 3 seconds using the coarse grained states. Investigating the accuracy of the inferred networks, we find that locaTE with coarse-grained cell infers more accurate static networks compared to direct application of locaTE to the full cell states. For inference of cell-specific networks, we find that locaTE with coarse-grained cell states results in a modest decrease in performance as measured by the AUPRC. This may be an acceptable compromise however, considering the drastic speedup by several orders of magnitude.

#### Mouse embryonic stem cell dataset

Log-transformed expression values for 100 TFs and pseudotime estimates were fetched from the SCODE example “data1” [18]. Cells were embedded into the top 25 principal components and a *k*-NN graph was constructed with *k* = 25. Using this *k*-NN graph, a cost matrix *C* of squared pairwise shortest-path distances was calculated using the Floyd-Warshall algorithm. StationaryOT [39] was then applied using this cost matrix, treating cells in the final 10% of pseudotime as sink cells. The flux rate for sink cells was taken to be uniform such that ∑_*i*∈sinks_ *R*_*i*_ = −50, and similarly for non-sink cells so that ∑_*i*∉sinks_ *R*_*i*_ = −50. Quadratically regularised optimal transport was applied with 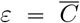, i.e. the mean value of *C*. The resulting transition matrix was used as the forward transition matrix for locaTE. The ESCAPE reference network of [19] was retrieved from https://github.com/gitter-lab/SINGE-supplemental. LocaTE was applied to estimate TE using the transition kernel derived from StationaryOT with *τ* = 1. Expression values were discretised using the Bayesian blocks algorithm as implemented in [28], with artifactually small values (< 10^−2^) set to zero in order to prevent many bins being created near zero. Raw TE values were first filtered using weighted CLR, and denoising regression was applied with *λ*_1_ = 10, *λ*_2_ = 10^−3^ using the normalised *k*-NN graph Laplacian. To apply locaTE-NMF, the array of raw TE values was scaled to have unit 0.9-quantile. The target rank was set to *k* = 10, *L* was taken to be the normalised *k*-NN graph Laplacian, and *K*_1_ = *K*_2_ = *I*. We chose *α* = 5, *β* = 0, *λ*_1_ = *λ*_2_ = *µ*_1_ = *µ*_2_ = 1. For comparison, PIDC [28], GRISLI [20], TENET [17], SCODE [18] and CeSpGRN [11] were applied using the same log-transformed expression values and pseudotime estimates. For TENET a history length of 1 was used, following the usage guidelines in its documentation. We found that TENET tended to perform worse with longer history lengths. For SCODE, the rank parameter was set to *D* = 4 which is the recommended value. For CeSpGRN, we took a neighbourhood size |*N*(*i*)| = 25, bandwidth *σ* = 5, and sparsity level *λ* = 0.1 in keeping with the suggested parameter ranges in [11, Section 3.1]. All other parameters were taken to be their default values.

#### Pancreatic endocrinogenesis dataset

Expression data was processed and RNA velocity was estimated following the CellRank tutorial available at https://cellrank.readthedocs.io/en/stable/cellrank_basics.html. A transition kernel derived from RNA velocity was produced using cellrank.tl.kernels-.VelocityKernel using the cosine similarity scheme. Pseudotime estimates for visualisation were calculated using the function scvelo.tl.latent_time. Intersecting the genes that passed filtering with the list of TFs from [20] yielded a subset of 82 TFs. LocaTE was applied using log-transformed expression values of the TF subset and the transition kernel derived from RNA velocity, with *τ* = 1. All other parameters were chosen to be the same as for the mESC dataset. Prior to analysis of the inferred networks, we filtered out edges involving the stress-related genes Fos and Egr1 as described in the main text.

#### Haematopoiesis (Marot-Lassauzaie et al.) dataset

Expression data was obtained from the companion Github repository of [41] available at https://github.com/HaghverdiLab/velocity_notebooks and a transition kernel derived from RNA velocity was produced following the code available in HSPC_kappa-velo.ipynb. Log-normalised gene expression matrices were prepared by filtering and normalisation using the function scvelo.pp.filter_and_normalize function with min_shared_counts = 20 and n_top_genes = 2000. Intersecting the genes that passed filtering with the list of Mus musculus TFs from [121] available at http://bioinfo.life.hust.edu.cn/static/AnimalTFDB3/download/Mus_musculus_TF yielded a subset of 202 TFs, from which we further filtered for the top 100 highly variable TFs using the function sc.pp.highly_variable_genes. Velocity pseudotime was estimated from the *κ*-velo dynamics using the function scvelo.tl.velocity_pseudotime. LocaTE was applied using the log-transformed expression values of the TF subset and the transition kernel derived from *κ*-velo with *τ* = 1. All other parameters for locaTE were chosen to be the same as for the mESC dataset.

#### Haematopoiesis (Tusi et al.) dataset

Preprocessed single-cell expression data was obtained from the Gene Expression Omnibus (GEO) using accession GSE89754 from GSM2388072_basal_bone_marrow.filtered_-normalized_counts.csv.gz. Due to batch effects described in [84, Methods], cells from the library basal_-bm1 were discarded. Preprocessed gene expression matrices were prepared by filtering and normalisation with sc.pp.normalize_per_cell followed by a log1p-transformation. We retained the top 100 highly variable TFs, as was described previously for the Marot-Lassauzaie et al. dataset. To focus on cells close to the fate decision boundary, we filtered out cells estimated by PBA to be highly fate committed, keeping only cells with PBA_Potential > 20. LocaTE was applied using the log-transformed expression values of the TF subset and a transition kernel derived from PBA with *τ* = 3. Results are shown in Figure S19.

#### Cell-cycle (Battich et al.) dataset

ScEU-seq metabolic labelling data of [86] was obtained and pre-processed following the tutorial of Dynamo [24] available at https://dynamo-release.readthedocs.io/en/latest/notebooks/scEU_seq_rpe1_analysis_kinetic.html. First we selected the top 1,000 highly variable genes using scanpy.pp.highly_variable_genes, before computing for each gene its Spearman correlation with the provided relative cell cycle position Cell_cycle_relativePos. We selected the 200 most strongly correlated (in terms of the absolute value) genes for network inference. Dynamo [24] was employed to estimate cell-wise velocities from metabolic labelling data, after which a global vector field was learned in 30-dimensional PCA space using dyn.vf.VectorField. A transition matrix was calculated from the global vector field using the dot product kernel and the *k*-NN graph with *k* = 50. LocaTE and locaTE-NMF were applied using the normalised expression matrix with *τ* = 1. All other parameters were chosen to be the same as for the mESC dataset (see Section 5). Results are shown in Figure S21.

## Acknowledgements

SYZ would like to thank Geoffrey Schiebinger, Léo Diaz and Wilder Scott for insightful discussions. The first author acknowledges support from the Australian Government Research Training Program and an Elizabeth and Vernon Puzey Scholarship. MPHS gratefully acknowledges funding from the Australian Research Council through a Laureate Fellowship (FL220100005), and the University of Melbourne through DRM funding.

## Footnotes

1 although GENIE3 produces directed networks as output, since it does not use any temporal or causal information, we find that its inference outputs are almost always close to being symmetric.

## Notes

https://github.com/zsteve/locaTE.jl

### Competing Interest Statement

The authors have declared no competing interest.

### Summary of Updates

Main text and figures revised. Portions of supplementary text moved to ``Methods'' and ``Theory'' sections.

## References

[1] Florian Buettner, Kedar N Natarajan, F Paolo Casale, Valentina Proserpio, Antonio Scialdone, Fabian J Theis, Sarah A Teichmann, John C Marioni, and Oliver Stegle. Computational analysis of cell-to-cell heterogeneity in single-cell rna-sequencing data reveals hidden subpopulations of cells. Nature biotechnology, 33(2):155–160, 2015.

[2] Bradley Efron and Trevor Hastie. Computer Age Statistical Inference, Student Edition: Algorithms, Evidence, and Data Science, volume 6. Cambridge University Press, 2021.

[3] Michael PH Stumpf. Inferring better gene regulation networks from single-cell data. Current Opinion in Systems Biology, 27:100342, 2021.

[4] Kyle Akers and TM Murali. Gene regulatory network inference in single-cell biology. Current Opinion in Systems Biology, 26:87–97, 2021.

[5] Shuonan Chen and Jessica C Mar. Evaluating methods of inferring gene regulatory networks highlights their lack of performance for single cell gene expression data. BMC bioinformatics, 19(1):232–221, 2018.

[6] Aditya Pratapa, Amogh P Jalihal, Jeffrey N Law, Aditya Bharadwaj, and TM Murali. Benchmarking algorithms for gene regulatory network inference from single-cell transcriptomic data. Nature methods, 17(2):147–154, 2020.

[7] Patrick S Stumpf, Rosanna CG Smith, Michael Lenz, Andreas Schuppert, Franz-Josef Müller, Ann Babtie, Thalia E Chan, Michael PH Stumpf, Colin P Please, Sam D Howison, et al. Stem cell differentiation as a non-markov stochastic process. Cell Systems, 5(3):268–282, 2017.

[8] Elias Ventre, Ulysse Herbach, Thibault Espinasse, Gérard Benoit, and Olivier Gandrillon. One model fits all: combining inference and simulation of gene regulatory networks. bioRxiv, 2022.

[9] Thomas W Thorne, Pietro Fratta, Michael G Hanna, Andrea Cortese, Vincent Plagnol, Elizabeth M Fisher, Michael P H Stumpf, and Michael P H Stumpf. Graphical modelling of molecular networks underlying sporadic inclusion body myositis. Molecular Biosystems, 9(7):1736–1742, 2013.

[10] Robrecht Cannoodt, Wouter Saelens, Louise Deconinck, and Yvan Saeys. Spearheading future omics analyses using dyngen, a multi-modal simulator of single cells. Nature Communications, 12(1):1–9, 2021.

[11] Ziqi Zhang, Jongseok Han, L. Song, and Xiuwei Zhang. Inferring cell-specific gene regulatory networks from single cell gene expression data. bioRxiv, 2022.

[12] Xuran Wang, David Choi, and Kathryn Roeder. Constructing local cell-specific networks from single-cell data. Proceedings of the National Academy of Sciences, 118(51), 2021.

[13] Hao Dai, Lin Li, Tao Zeng, and Luonan Chen. Cell-specific network constructed by single-cell rna sequencing data. Nucleic acids research, 47(11):e62–e62, 2019.

[14] Federico Bocci, Peijie Zhou, and Qing Nie. splicejac: transition genes and state-specific gene regulation from single-cell transcriptome data. Molecular Systems Biology, 18(11):e11176, 2022.

[15] Xiaojie Qiu, Arman Rahimzamani, Li Wang, Bingcheng Ren, Qi Mao, Timothy Durham, José L McFaline-Figueroa, Lauren Saunders, Cole Trapnell, and Sreeram Kannan. Inferring causal gene regulatory networks from coupled single-cell expression dynamics using scribe. Cell systems, 10(3):265– 274, 2020.

[16] Sophie Lèbre, Jennifer Becq, Frédéric Devaux, Michael P H Stumpf, and Gaëlle Lelandais. Statistical inference of the time-varying structure of gene-regulation networks. BMC Systems Biology, 4(1):130, 2010.

[17] Junil Kim, Simon T. Jakobsen, Kedar N Natarajan, and Kyoung-Jae Won. Tenet: gene network reconstruction using transfer entropy reveals key regulatory factors from single cell transcriptomic data. Nucleic acids research, 49(1):e1–e1, 2021.

[18] Hirotaka Matsumoto, Hisanori Kiryu, Chikara Furusawa, Minoru SH Ko, Shigeru BH Ko, Norio Gouda, Tetsutaro Hayashi, and Itoshi Nikaido. Scode: an efficient regulatory network inference algorithm from single-cell rna-seq during differentiation. Bioinformatics, 33(15):2314–2321, 2017.

[19] Atul Deshpande, Li-Fang Chu, Ron Stewart, and Anthony Gitter. Network inference with granger causality ensembles on single-cell transcriptomics. Cell reports, 38(6):110333, 2022.

[20] Pierre-Cyril Aubin-Frankowski and Jean-Philippe Vert. Gene regulation inference from single-cell rna-seq data with linear differential equations and velocity inference. Bioinformatics, 36(18):4774–4780, 2020.

[21] Caleb Weinreb, Samuel Wolock, Betsabeh K Tusi, Merav Socolovsky, and Allon M Klein. Fundamental limits on dynamic inference from single-cell snapshots. Proceedings of the National Academy of Sciences, 115(10):E2467–E2476, 2018.

[22] Geoffrey Schiebinger. Reconstructing developmental landscapes and trajectories from single-cell data. Current Opinion in Systems Biology, 27:100351, 2021.

[23] Marius Lange, Volker Bergen, Michal Klein, Manu Setty, Bernhard Reuter, Mostafa Bakhti, Heiko Lickert, Meshal Ansari, Janine Schniering, Herbert B Schiller, et al. Cellrank for directed single-cell fate mapping. Nature methods, pages 1–12, 2022.

[24] Xiaojie Qiu, Yan Zhang, Jorge D Martin-Rufino, Chen Weng, Shayan Hosseinzadeh, Dian Yang, Angela N Pogson, Marco Y Hein, Kyung Hoi Joseph Min, Li Wang, et al. Mapping transcriptomic vector fields of single cells. Cell, 185(4):690–711, 2022.

[25] Kevin R Moon, Jay S Stanley Iii, Daniel Burkhardt, David van Dijk, Guy Wolf, and Smita Krishnaswamy. Manifold learning-based methods for analyzing single-cell rna-sequencing data. Current Opinion in Systems Biology, 7:36–46, 2018.

[26] Juliane Liepe, Aaron Sim, Helen Weavers, Laura Ward, Paul Martin, and Michael PH Stumpf. Accurate reconstruction of cell and particle tracks from 3d live imaging data. Cell Systems, 3(1):102–107, 2016.

[27] Thomas Schreiber. Measuring information transfer. Physical review letters, 85(2):461, 2000.

[28] Thalia E Chan, Michael PH Stumpf, and Ann C Babtie. Gene regulatory network inference from single-cell data using multivariate information measures. Cell systems, 5(3):251–267, 2017.

[29] Lionel Barnett, Adam B Barrett, and Anil K Seth. Granger causality and transfer entropy are equivalent for gaussian variables. Physical review letters, 103(23):238701, 2009.

[30] Naomi Moris, Cristina Pina, and Alfonso Martinez Arias. Transition states and cell fate decisions in epigenetic landscapes. Nature Reviews Genetics, 17(11):693–703, 2016.

[31] Rowan D Brackston, Eszter Lakatos, and Michael P H Stumpf. Transition state characteristics during cell differentiation. PLoS computational biology, 14(9):e1006405, 2018.

[32] Conrad Hal Waddington. The strategy of the genes. 1957.

[33] Adam L MacLean, Tian Hong, and Qing Nie. Exploring intermediate cell states through the lens of single cells. Current Opinion in Systems Biology, 9:32–41, 2018.

[34] Peijie Zhou, Shuxiong Wang, Tiejun Li, and Qing Nie. Dissecting transition cells from single-cell transcriptome data through multiscale stochastic dynamics. Nature communications, 12(1):5609, 2021.

[35] Sophie Tritschler, Maren Büttner, David S Fischer, Marius Lange, Volker Bergen, Heiko Lickert, and Fabian J Theis. Concepts and limitations for learning developmental trajectories from single cell genomics. Development, 146(12):dev170506, 2019.

[36] Ronald R Coifman and Stéphane Lafon. Diffusion maps. Applied and computational harmonic analysis, 21(1):5–30, 2006.

[37] Volker Bergen, Marius Lange, Stefan Peidli, F Alexander Wolf, and Fabian J Theis. Generalizing rna velocity to transient cell states through dynamical modeling. Nature biotechnology, 38(12):1408–1414, 2020.

[38] Laleh Haghverdi, Maren Büttner, F Alexander Wolf, Florian Buettner, and Fabian J Theis. Diffusion pseudotime robustly reconstructs lineage branching. Nature methods, 13(10):845–848, 2016.

[39] Stephen Zhang, Anton Afanassiev, Laura Greenstreet, Tetsuya Matsumoto, and Geoffrey Schiebinger. Optimal transport analysis reveals trajectories in steady-state systems. PLoS computational biology, 17(12):e1009466, 2021.

[40] Gennady Gorin, Meichen Fang, Tara Chari, and Lior Pachter. Rna velocity unraveled. bioRxiv, 2022.

[41] Valérie Marot-Lassauzaie, Brigitte Joanne Bouman, Fearghal Declan Donaghy, Yasmin Demerdash, Marieke Alida Gertruda Essers, and Laleh Haghverdi. Towards reliable quantification of cell state velocities. PLoS Computational Biology, 18(9):e1010031, 2022.

[42] Yan Zhang, Xiaojie Qiu, Jonathan S Weissman, Ivet Bahar, and Jianhua Xing. Graph-dynamo: Learning stochastic cellular state transition dynamics from single cell data. bioRxiv, pages 2023–09, 2023.

[43] Steven L Brunton, Joshua L Proctor, and J Nathan Kutz. Discovering governing equations from data by sparse identification of nonlinear dynamical systems. Proceedings of the national academy of sciences, 113(15):3932–3937, 2016.

[44] Tiejun Li, Jifan Shi, Yichong Wu, and Peijie Zhou. On the mathematics of rna velocity i: theoretical analysis. bioRxiv, 2020.

[45] Jeremiah J Faith, Boris Hayete, Joshua T Thaden, Ilaria Mogno, Jamey Wierzbowski, Guillaume Cottarel, Simon Kasif, James J Collins, and Timothy S Gardner. Large-scale mapping and validation of escherichia coli transcriptional regulation from a compendium of expression profiles. PLoS biology, 5(1):e8, 2007.

[46] Vân Anh Huynh-Thu, Alexandre Irrthum, Louis Wehenkel, and Pierre Geurts. Inferring regulatory networks from expression data using tree-based methods. PloS one, 5(9):e12776, 2010.

[47] Nan Papili Gao, SM Minhaz Ud-Dean, Olivier Gandrillon, and Rudiyanto Gunawan. Sincerities: inferring gene regulatory networks from time-stamped single cell transcriptional expression profiles. Bioinformatics, 34(2):258–266, 2018.

[48] Tetsutaro Hayashi, Haruka Ozaki, Yohei Sasagawa, Mana Umeda, Hiroki Danno, and Itoshi Nikaido. Single-cell full-length total rna sequencing uncovers dynamics of recursive splicing and enhancer rnas. Nature communications, 9(1):1–16, 2018.

[49] Huilei Xu, Caroline Baroukh, Ruth Dannenfelser, Edward Y Chen, Christopher M Tan, Yan Kou, Yujin E Kim, Ihor R Lemischka, and Avi Ma’ayan. Escape: database for integrating high-content published data collected from human and mouse embryonic stem cells. Database, 2013, 2013.

[50] Wenjing Shi, Hui Wang, Guangjin Pan, Yijie Geng, Yunqian Guo, and Duanqing Pei. Regulation of the pluripotency marker rex-1 by nanog and sox2. Journal of biological chemistry, 281(33):23319–23325, 2006.

[51] Guangjin Pan and James A Thomson. Nanog and transcriptional networks in embryonic stem cell pluripotency. Cell research, 17(1):42–49, 2007.

[52] Chin Yan Lim, Wai-Leong Tam, Jinqiu Zhang, Haw Siang Ang, Hui Jia, Leonard Lipovich, Huck-Hui Ng, Chia-Lin Wei, Wing Kin Sung, Paul Robson, et al. Sall4 regulates distinct transcription circuitries in different blastocyst-derived stem cell lineages. Cell stem cell, 3(5):543–554, 2008.

[53] Kelly L Covello, James Kehler, Hongwei Yu, John D Gordan, Andrew M Arsham, Cheng-Jun Hu, Patricia A Labosky, M Celeste Simon, and Brian Keith. Hif-2α regulates oct-4: effects of hypoxia on stem cell function, embryonic development, and tumor growth. Genes & development, 20(5):557–570, 2006.

[54] Hong Doan, Alexander Parsons, Shruthi Devkumar, Jogitha Selvarajah, Francesc Miralles, and Veronica A Carroll. Hif-mediated suppression of deptor confers resistance to mtor kinase inhibition in renal cancer. Iscience, 21:509–520, 2019.

[55] Pooja Agrawal, Joseph Reynolds, Shereen Chew, Deepak A Lamba, and Robert E Hughes. Deptor is a stemness factor that regulates pluripotency of embryonic stem cells. Journal of Biological Chemistry, 289(46):31818–31826, 2014.

[56] Edward E Morrisey, Zhihua Tang, Kirsten Sigrist, Min Min Lu, Fang Jiang, Hon S Ip, and Michael S Parmacek. Gata6 regulates hnf4 and is required for differentiation of visceral endoderm in the mouse embryo. Genes & development, 12(22):3579–3590, 1998.

[57] Yingyao Zhou, Bin Zhou, Lars Pache, Max Chang, Alireza Hadj Khodabakhshi, Olga Tanaseichuk, Christopher Benner, and Sumit K Chanda. Metascape provides a biologist-oriented resource for the analysis of systems-level datasets. Nature communications, 10(1):1–10, 2019.

[58] Sayali Chowdhary and Anna-Katerina Hadjantonakis. Journey of the mouse primitive endoderm: from specification to maturation. Philosophical Transactions of the Royal Society B, 377(1865):20210252, 2022.

[59] Irene Aksoy, Ralf Jauch, Jiaxuan Chen, Mateusz Dyla, Ushashree Divakar, Gireesh K Bogu, Roy Teo, Calista Keow Leng Ng, Wishva Herath, Sun Lili, et al. Oct4 switches partnering from sox2 to sox17 to reinterpret the enhancer code and specify endoderm. The EMBO journal, 32(7):938–953, 2013.

[60] Aimée Bastidas-Ponce, Sophie Tritschler, Leander Dony, Katharina Scheibner, Marta Tarquis-Medina, Ciro Salinno, Silvia Schirge, Ingo Burtscher, Anika Böttcher, Fabian J Theis, et al. Comprehensive single cell mrna profiling reveals a detailed roadmap for pancreatic endocrinogenesis. Development, 146(12):dev173849, 2019.

[61] Ashleigh E Schaffer, Brandon L Taylor, Jacqueline R Benthuysen, Jingxuan Liu, Fabrizio Thorel, Weiping Yuan, Yang Jiao, Klaus H Kaestner, Pedro L Herrera, Mark A Magnuson, et al. Nkx6. 1 controls a gene regulatory network required for establishing and maintaining pancreatic beta cell identity. PLoS genetics, 9(1):e1003274, 2013.

[62] EC Baechler, FM Batliwalla, George Karypis, PM Gaffney, K Moser, WA Ortmann, KJ Espe, S Balasubramanian, KM Hughes, JP Chan, et al. Expression levels for many genes in human peripheral blood cells are highly sensitive to ex vivo incubation. Genes & Immunity, 5(5):347–353, 2004.

[63] Léo Machado, Frédéric Relaix, and Philippos Mourikis. Stress relief: emerging methods to mitigate dissociation-induced artefacts. Trends in Cell Biology, 31(11):888–897, 2021.

[64] Brandon L Taylor, Fen-Fen Liu, and Maike Sander. Nkx6. 1 is essential for maintaining the functional state of pancreatic beta cells. Cell reports, 4(6):1262–1275, 2013.

[65] >H Efsun Arda, Cecil M Benitez, and Seung K Kim. Gene regulatory networks governing pancreas development. Developmental cell, 25(1):5–13, 2013.

[66] Philip A Seymour, Kristine K Freude, Man N Tran, Erin E Mayes, Jan Jensen, Ralf Kist, Gerd Scherer, and Maike Sander. Sox9 is required for maintenance of the pancreatic progenitor cell pool. Proceedings of the National Academy of Sciences, 104(6):1865–1870, 2007.

[67] Richard I Sherwood, Tzong-Yang Albert Chen, and Douglas A Melton. Transcriptional dynamics of endodermal organ formation. Developmental dynamics: an official publication of the American Association of Anatomists, 238(1):29–42, 2009.

[68] Shin-Heng Chiou, Madeleine Dorsch, Eva Kusch, Santiago Naranjo, Margaret M Kozak, Albert C Koong, Monte M Winslow, and Barbara M Grüner. Hmga2 is dispensable for pancreatic cancer development, metastasis, and therapy resistance. Scientific reports, 8(1):1–10, 2018.

[69] C Dorrell, J Schug, CF Lin, PS Canaday, AJ Fox, O Smirnova, R Bonnah, PR Streeter, CJ Stoeckert, KH Kaestner, et al. Transcriptomes of the major human pancreatic cell types. Diabetologia, 54:2832– 2844, 2011.

[70] Kailun Lee, Jeng Yie Chan, Cassandra Liang, Chi Kin Ip, Yan-Chuan Shi, Herbert Herzog, William E Hughes, Mohammed Bensellam, Viviane Delghingaro-Augusto, Mark E Koina, et al. Xbp1 maintains beta cell identity, represses beta-to-alpha cell transdifferentiation and protects against diabetic beta cell failure during metabolic stress in mice. Diabetologia, 65(6):984–996, 2022.

[71] Ingmar Glauche and Carsten Marr. Mechanistic models of blood cell fate decisions in the era of single-cell data. Current Opinion in Systems Biology, 28:100355, 2021.

[72] Jan Krumsiek, Carsten Marr, Timm Schroeder, and Fabian J Theis. Hierarchical differentiation of myeloid progenitors is encoded in the transcription factor network. PloS one, 6(8):e22649, 2011.

[73] Shijie C Zheng, Genevieve Stein-O’Brien, Leandros Boukas, Loyal A Goff, and Kasper D Hansen. Pumping the brakes on rna velocity–understanding and interpreting rna velocity estimates. bioRxiv, page 2022–06, 2022.

[74] Cuiping Zhang, Yvonne N Fondufe-Mittendorf, Chi Wang, Jin Chen, Qiang Cheng, Daohong Zhou, Yi Zheng, Hartmut Geiger, and Ying Liang. Latexin regulation by hmgb2 is required for hematopoietic stem cell maintenance. haematologica, 105(3):573, 2020.

[75] Michael J Nemeth, Martha R Kirby, and David M Bodine. Hmgb3 regulates the balance between hematopoietic stem cell self-renewal and differentiation. Proceedings of the National Academy of Sciences, 103(37):13783–13788, 2006.

[76] Xunde Wang, Nikolaos Angelis, and Swee Lay Thein. Myb–a regulatory factor in hematopoiesis. Gene, 665:6–17, 2018.

[77] Thomas Graf and Tariq Enver. Forcing cells to change lineages. Nature, 462(7273):587–594, 2009.

[78] Rachel M Gerstein. Deciding the decider: Mef2c in hematopoiesis. Nature Immunology, 10(3):235–236, 2009.

[79] Hongsheng Wang and Herbert C Morse. Irf8 regulates myeloid and b lymphoid lineage diversification. Immunologic research, 43:109–117, 2009.

[80] Tomohiko Tamura, Daisuke Kurotaki, and Shin-ichi Koizumi. Regulation of myelopoiesis by the transcription factor irf8. International journal of hematology, 101:342–351, 2015.

[81] Voahangy Randrianarison-Huetz, Benoit Laurent, Valérie Bardet, Gerard C Blobe, François Huetz, and Dominique Duménil. Gfi-1b controls human erythroid and megakaryocytic differentiation by regulating tgf-β signaling at the bipotent erythro-megakaryocytic progenitor stage. Blood, The Journal of the American Society of Hematology, 115(14):2784–2795, 2010.

[82] Emmanuelle Passegué, Erwin F Wagner, and Irving L Weissman. Junb deficiency leads to a myeloproliferative disorder arising from hematopoietic stem cells. Cell, 119(3):431–443, 2004.

[83] Karthik Arumugam, William Shin, Valentina Schiavone, Lukas Vlahos, Xiaochuan Tu, Davide Carnevali, Jordan Kesner, Evan O Paull, Neus Romo, Prem Subramaniam, et al. The master regulator protein baz2b can reprogram human hematopoietic lineage-committed progenitors into a multipotent state. Cell reports, 33(10):108474, 2020.

[84] Betsabeh Khoramian Tusi, Samuel L Wolock, Caleb Weinreb, Yung Hwang, Daniel Hidalgo, Rapolas Zilionis, Ari Waisman, Jun R Huh, Allon M Klein, and Merav Socolovsky. Population snapshots predict early haematopoietic and erythroid hierarchies. Nature, 555(7694):54–60, 2018.

[85] Terry Bossomaier, Lionel Barnett, Michael Harré, and Joseph T Lizier. Transfer entropy. In An introduction to transfer entropy, pages 65–95. Springer, 2016.

[86] Nico Battich, Joep Beumer, Buys de Barbanson, Lenno Krenning, Chloé S Baron, Marvin E Tanenbaum, Hans Clevers, and Alexander van Oudenaarden. Sequencing metabolically labeled transcripts in single cells reveals mrna turnover strategies. Science, 367(6482):1151–1156, 2020.

[87] Lam-Ha Ly and Martin Vingron. Effect of imputation on gene network reconstruction from single-cell rna-seq data. Patterns, 3(2):100414, 2022.

[88] Juan Zhao, Yiwei Zhou, Xiujun Zhang, and Luonan Chen. Part mutual information for quantifying direct associations in networks. Proceedings of the National Academy of Sciences, 113(18):5130–5135, 2016.

[89] Shou-Wen Wang, Michael J Herriges, Kilian Hurley, Darrell N Kotton, and Allon M Klein. Cospar identifies early cell fate biases from single-cell transcriptomic and lineage information. Nature Biotechnology, 40(7):1066–1074, 2022.

[90] Louise Deconinck, Robrecht Cannoodt, Wouter Saelens, Bart Deplancke, and Yvan Saeys. Recent advances in trajectory inference from single-cell omics data. Current Opinion in Systems Biology, 27:100344, 2021.

[91] Jun Ding, Nadav Sharon, and Ziv Bar-Joseph. Temporal modelling using single-cell transcriptomics. Nature Reviews Genetics, pages 1–14, 2022.

[92] Kun Wang, Liangzhen Hou, Xin Wang, Xiangwei Zhai, Zhaolian Lu, Zhike Zi, Weiwei Zhai, Xionglei He, Christina Curtis, D. Zhou, et al. Phylovelo enhances transcriptomic velocity field mapping using monotonically expressed genes. Nature Biotechnology, 42(5):778–789, 2024.

[93] Joseph CF Ng, Guillem Montamat Garcia, Alexander T Stewart, Paul Blair, Claudia Mauri, Deborah K Dunn-Walters, and Franca Fraternali. scicsr infers b cell state transition and predicts class-switch recombination dynamics using single-cell transcriptomic data. Nature Methods, 21(5):823–834, 2024.

[94] Gennady Gorin and Lior Pachter. Length biases in single-cell rna sequencing of pre-mrna. Biophysical Reports, 3(1), 2023.

[95] Rory J Maizels, Daniel M Snell, and James Briscoe. Reconstructing developmental trajectories using latent dynamical systems and time-resolved transcriptomics. Cell Systems, 15(5):411–424, 2024.

[96] MA Coomer, L Ham, and MPH Stumpf. Noise distorts the epigenetic landscape and shapes cell-fate decisions. Cell Syst, 13(1):83–102, 2022.

[97] Gunsagar S Gulati, Shaheen S Sikandar, Daniel J Wesche, Anoop Manjunath, Anjan Bharadwaj, Mark J Berger, Francisco Ilagan, Angera H Kuo, Robert W Hsieh, Shang Cai, et al. Single-cell transcriptional diversity is a hallmark of developmental potential. Science, 367(6476):405–411, 2020.

[98] Andrew E Teschendorff and Tariq Enver. Single-cell entropy for accurate estimation of differentiation potency from a cell’s transcriptome. Nature communications, 8(1):15599, 2017.

[99] Martina Tedesco, Francesca Giannese, Dejan Lazarević, Valentina Giansanti, Dalia Rosano, Silvia Monzani, Irene Catalano, Elena Grassi, Eugenia R Zanella, Oronza A Botrugno, et al. Chromatin velocity reveals epigenetic dynamics by single-cell profiling of heterochromatin and euchromatin. Nature biotechnology, 40(2):235–244, 2022.

[100] Ulysse Herbach, Arnaud Bonnaffoux, Thibault Espinasse, and Olivier Gandrillon. Inferring gene regulatory networks from single-cell data: a mechanistic approach. BMC systems biology, 11(1):1–15, 2017.

[101] Thomas M. Cover and Joy A. Thomas. Elements Of Information Theory. J. Wiley, 2005.

[102] Clive WJ Granger. Investigating causal relations by econometric models and cross-spectral methods. Econometrica: journal of the Econometric Society, pages 424–438, 1969.

[103] Steven L Bressler and Anil K Seth. Wiener–granger causality: a well established methodology. Neuroimage, 58(2):323–329, 2011.

[104] John F Geweke. Measures of conditional linear dependence and feedback between time series. Journal of the American Statistical Association, 79(388):907–915, 1984.

[105] Judea Pearl et al. Models, reasoning and inference. Cambridge, UK: CambridgeUniversityPress, 19(2):3, 2000.

[106] Daniel T Gillespie. The chemical langevin equation. The Journal of Chemical Physics, 113(1):297–306, 2000.

[107] Crispin Gardiner. Stochastic Methods: A Handbook For The Natural And Social Sciences. Springer, 2009.

[108] Jie Sun and Erik M Bollt. Causation entropy identifies indirect influences, dominance of neighbors and anticipatory couplings. Physica D: Nonlinear Phenomena, 267:49–57, 2014.

[109] Ryan G James, Nix Barnett, and James P Crutchfield. Information flows? a critique of transfer entropies. Physical review letters, 116(23):238701, 2016.

[110] Pawel Czyz, Frederic Grabowski, Julia Vogt, Niko Beerenwinkel, and Alexander Marx. Beyond normal: On the evaluation of mutual information estimators. Advances in Neural Information Processing Systems, 36, 2024.

[111] Dmitry A Smirnov. Spurious causalities with transfer entropy. Physical Review E—Statistical, Nonlinear, and Soft Matter Physics, 87(4):042917, 2013.

[112] Aviv Madar, Alex Greenfield, Eric Vanden-Eijnden, and Richard Bonneau. Dream3: network inference using dynamic context likelihood of relatedness and the inferelator. PloS one, 5(3):e9803, 2010.

[113] Adam A Margolin, Ilya Nemenman, Katia Basso, Chris Wiggins, Gustavo Stolovitzky, Riccardo Dalla Favera, and Andrea Califano. Aracne: an algorithm for the reconstruction of gene regulatory networks in a mammalian cellular context. In BMC bioinformatics, volume 7, pages 1–15. BioMed Central, 2006.

[114] Stephen Boyd, Neal Parikh, Eric Chu, Borja Peleato, Jonathan Eckstein, et al. Distributed optimization and statistical learning via the alternating direction method of multipliers. Foundations and Trends® in Machine learning, 3(1):1–122, 2011.

[115] Daniel D Lee and H Sebastian Seung. Learning the parts of objects by non-negative matrix factorization. Nature, 401(6755):788–791, 1999.

[116] Jeff Bezanson, Alan Edelman, Stefan Karpinski, and Viral B Shah. Julia: A fresh approach to numerical computing. SIAM review, 59(1):65–98, 2017.

[117] Yael Baran, Akhiad Bercovich, Arnau Sebe-Pedros, Yaniv Lubling, Amir Giladi, Elad Chomsky, Zohar Meir, Michael Hoichman, Aviezer Lifshitz, and Amos Tanay. Metacell: analysis of single-cell rna-seq data using k-nn graph partitions. Genome biology, 20(1):1–19, 2019.

[118] Charles Miller Grinstead and James Laurie Snell. Introduction to probability. American Mathematical Soc., 2012.

[119] Stephen Zhang, Gilles Mordant, Tetsuya Matsumoto, and Geoffrey Schiebinger. Manifold learning with sparse regularised optimal transport. arXiv preprint 2307.09816, 2023.

[120] F Alexander Wolf, Fiona K Hamey, Mireya Plass, Jordi Solana, Joakim S Dahlin, Berthold Göttgens, Nikolaus Rajewsky, Lukas Simon, and Fabian J Theis. Paga: graph abstraction reconciles clustering with trajectory inference through a topology preserving map of single cells. Genome biology, 20:1–9, 2019.

[121] Hui Hu, Ya-Ru Miao, Long-Hao Jia, Qing-Yang Yu, Qiong Zhang, and An-Yuan Guo. Animaltfdb 3.0: a comprehensive resource for annotation and prediction of animal transcription factors. Nucleic acids research, 47(D1):D33–D38, 2019.

