## Supplementary information for "Learning cell-specific networks from dynamics and geometry of single cells"

#### S1 Supplementary results

##### S1.1 Simulated data

For each boolean network considered, simulated datasets were generated using the BoolODE package [1], modified to record additionally the velocity vector of simulated cells as well as the Jacobian matrix of expression state. We provide below the boolean rules that we used for simulating the two examples in the main text; all other simulated trajectories were produced using rules provided as part of the BoolODE package.

**Boolean rules: bifurcating**

$$
\begin{aligned}
g1 &\leftarrow \neg(g4 \vee g5)\\
g2 &\leftarrow g1\\
g3 &\leftarrow g2\\
g4 &\leftarrow ((g3 \vee g4) \wedge (\neg g5))\\
g5 &\leftarrow ((g5 \vee g3) \wedge (\neg g4))\\
g6 &\leftarrow ((g4 \wedge (\neg(g5 \vee g9))) \vee ((g7 \vee g6) \wedge (g5 \wedge (\neg g4))))\\
g7 &\leftarrow ((g6 \wedge (g4 \wedge (\neg g5))) \vee (g8 \wedge (g5 \wedge (\neg g4))))\\
g8 &\leftarrow ((g7 \wedge (g4 \wedge (\neg g5))) \vee (g9 \wedge (g5 \wedge (\neg g4))))\\
g9 &\leftarrow (((g8 \vee g9) \wedge (g4 \wedge (\neg g5))) \vee (g5 \wedge (\neg(g4 \vee g6))))
\end{aligned}
$$

See Figure S1.

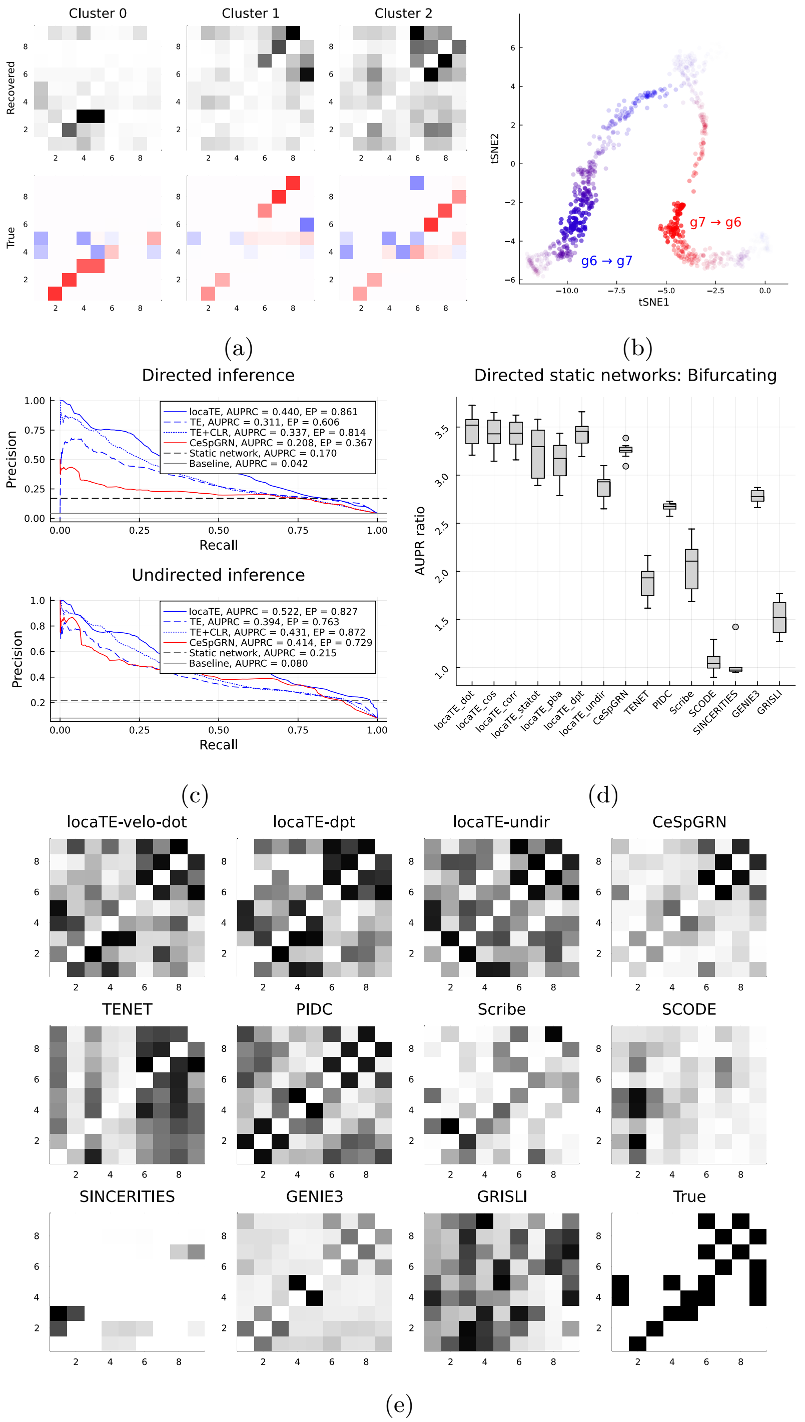


Supplementary Figure S1: **Supplementary results for simulated bifurcating network.** (a) Interactions inferred by locaTE averaged by cluster (top) shown against ground truth interactions averaged by cluster. (b) Interaction scores for $g7 \to g6$ and $g6 \to g7$ shown for each cell on the tSNE embedding. (c) Precision-Recall (PR) curves for cell-specific edge detection from $\hat{G}$ (raw TE values), $\tilde{G}$ (CLR filtered TE values), or $G$ (locaTE). Area under PR curve (AUPRC) and early precision at 10% recall (EP) is shown. (d) AUPRC ratio for locaTE and other methods across 10 sampled simulation datasets, for static (i.e. population average) inference. (e) Best inferred static networks.

**Boolean rules: switch**

$$
\begin{aligned}
g1 &\leftarrow (\neg g6)\\
g2 &\leftarrow ((g1 \wedge (\neg(g6 \wedge (\neg g5)))) \vee (g7 \wedge ((g6 \wedge (\neg g5)))))\\
g3 &\leftarrow ((g2 \wedge (\neg(g6 \wedge (\neg g5)))) \vee ((g4 \vee g3) \wedge ((g6 \wedge (\neg g5)))))\\
g4 &\leftarrow ((g3 \wedge (\neg(g6 \wedge (\neg g5)))) \vee (g2 \wedge ((g6 \wedge (\neg g5)))))\\
g5 &\leftarrow ((g4 \wedge (\neg(g6 \wedge (\neg g5)))))\\
g6 &\leftarrow ((g5 \vee g6))\\
g7 &\leftarrow (g5 \vee (g7 \wedge (g6 \wedge (\neg g5)) \wedge (\neg g2)))
\end{aligned}
$$

See Figure S2.

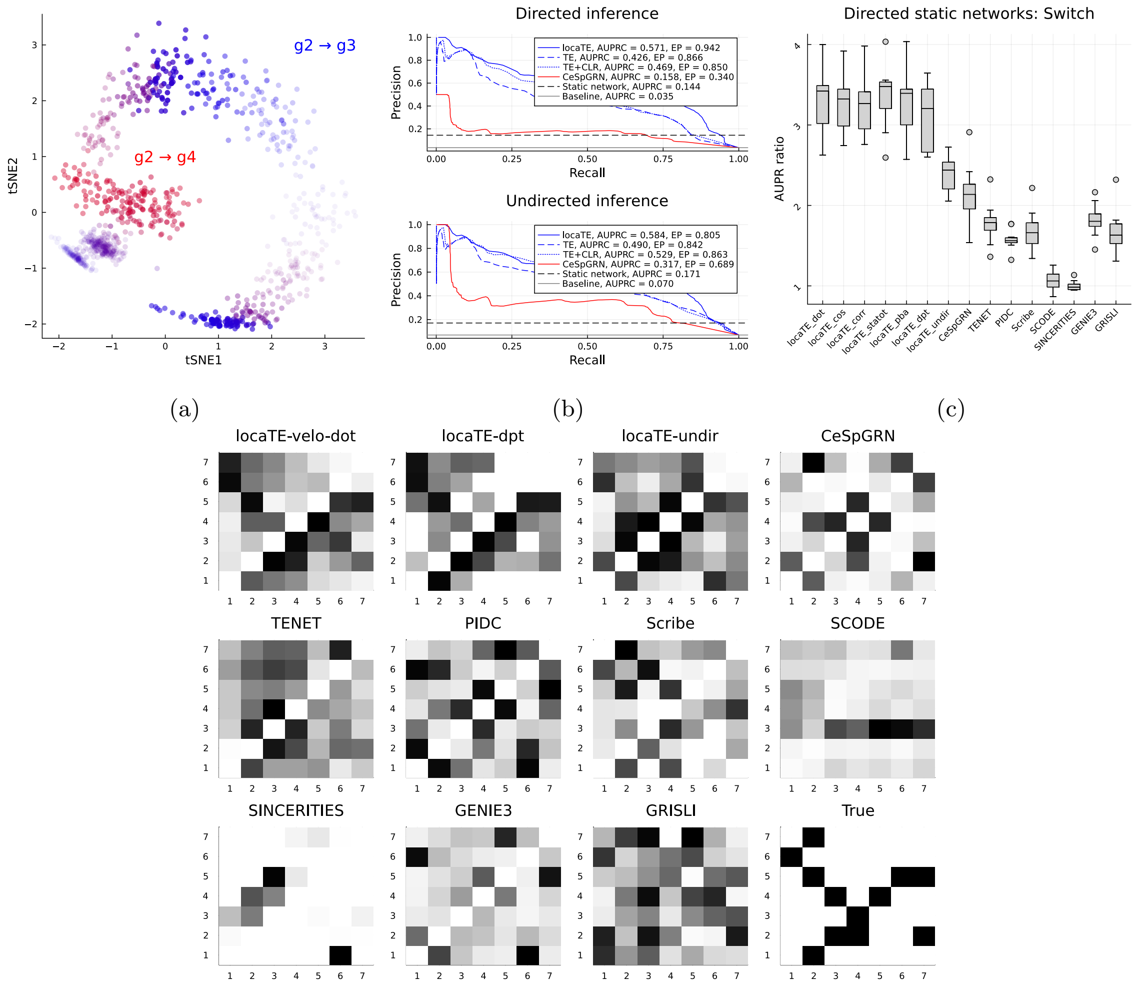


(d)

Supplementary Figure S2: **Supplementary results for simulated switch network.** (a) Interaction scores for $g2 \to g3$ and $g2 \to g4$ for each cell on the tSNE embedding. (b) Precision-Recall (PR) curves for cell-specific edge detection from $\hat{G}$ (raw TE values), $\tilde{G}$ (CLR filtered TE values), or $G$ (locaTE). Area under PR curve (AUPRC) and early precision at 10% recall (EP) is shown. (c) AUPRC ratio for locaTE and other methods across 10 sampled simulation datasets, for static (i.e. population average) inference. (d) Best inferred static networks.

**Parameter sweep results** See Figure S3.

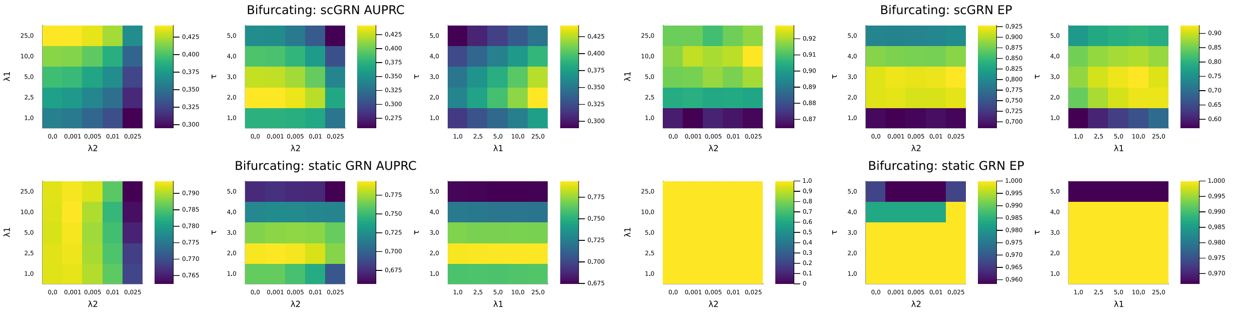


(a)

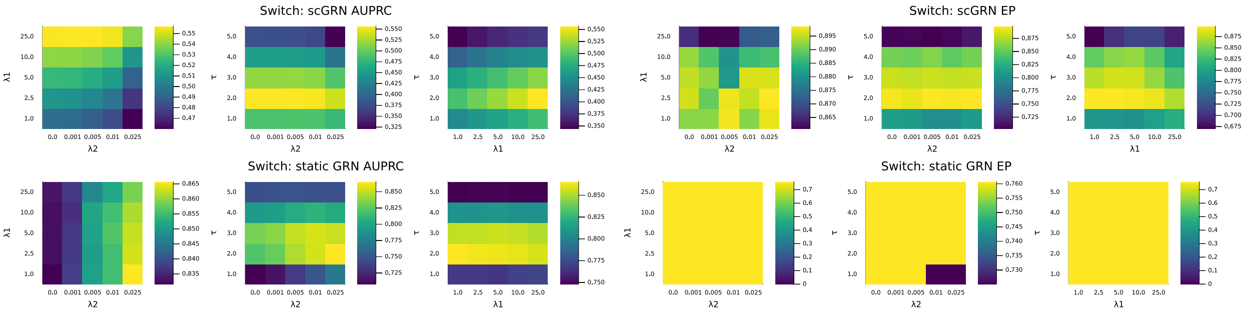


(b)

Supplementary Figure S3: Parameter sweep results over $(\tau, \lambda_1, \lambda_2)$ for (a) bifurcating simulation (b) switch simulation using dot-product velocity kernel. Heatmaps show, for each pair of parameters, the best average score (maximum AUPRC and maximum EP respectively) achieved along the remaining parameter.

**Time-scale selection heuristic** See Figure S4.

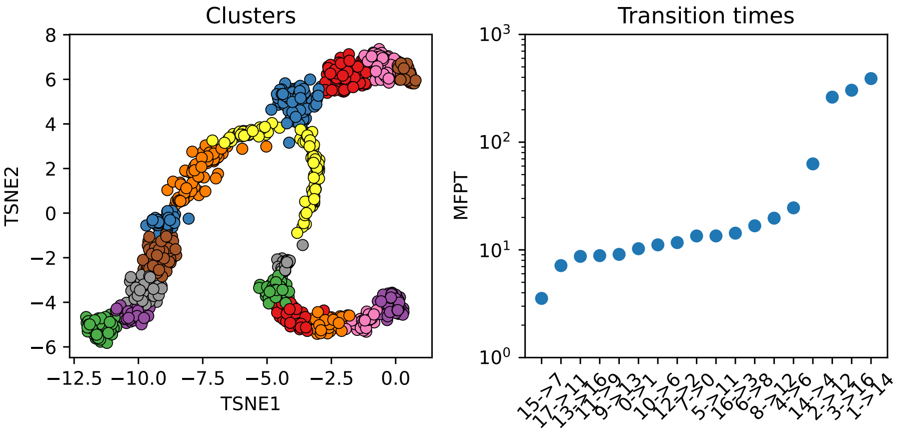


(a) Bifurcating system

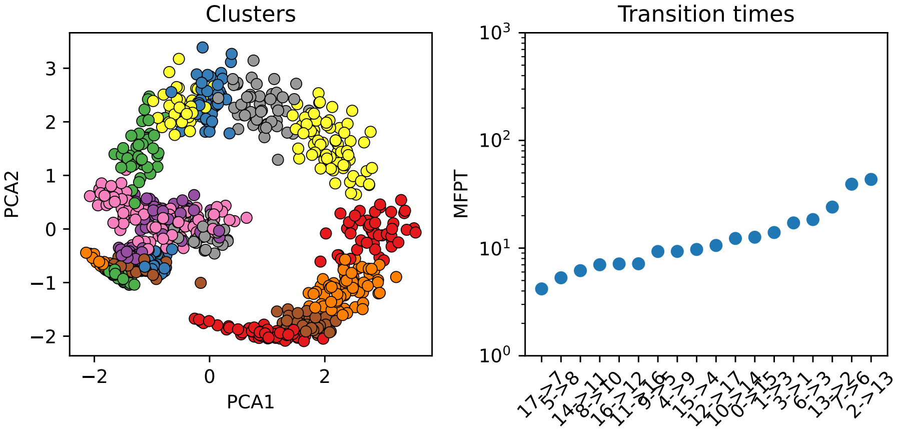


(b) Switch system

Supplementary Figure S4: Left: Leiden clusters found with `resolution = 2.5`, for $P_{\texttt{velo\_dot}}$. Right: Sorted mean-first passage times (MFPTs) between pairs of clusters. A heuristic for selecting $\tau$ (see *Methods*) is to pick $\tau$ on the order of the shortest transition time between any pair of clusters.

#### S1.2 Results: BoolODE trajectories

We applied locaTE with several choices of kernel to the simulated trajectory examples from [1]. For comparison, we also applied CeSpGRN, TENET, PIDC, Scribe and SCODE. A parameter sweep was performed for locaTE, CeSpGRN and TENET, and default parameters were used for the remaining methods. Performance for each method was assessed in terms of the AUPRC ratio for directed edge inference. We summarise performance with boxplots below, and show representative examples of the recovered static networks.

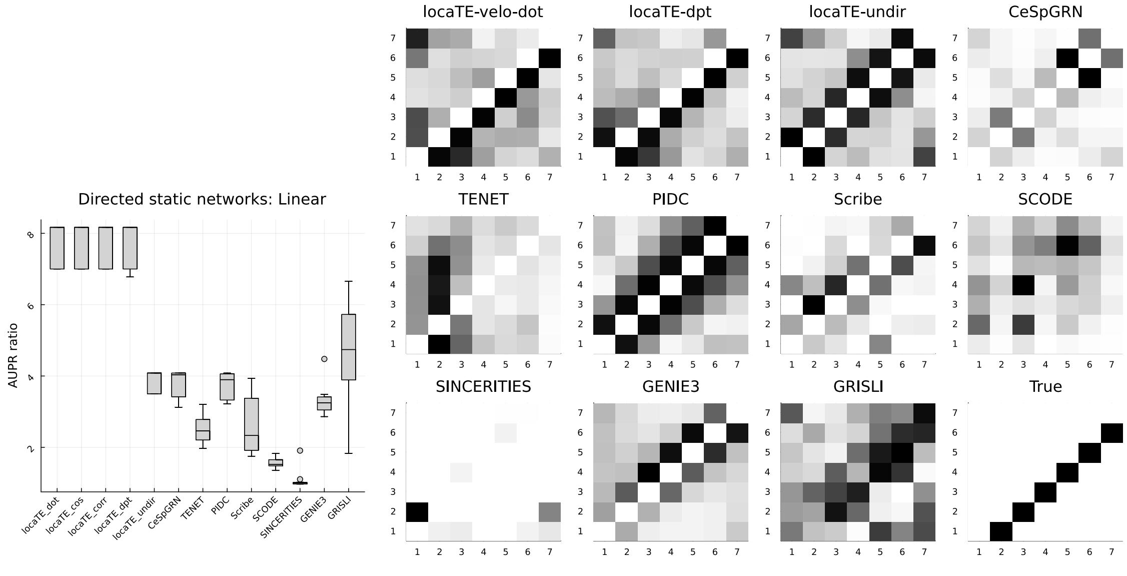


(a) Linear network, 1000 cells.

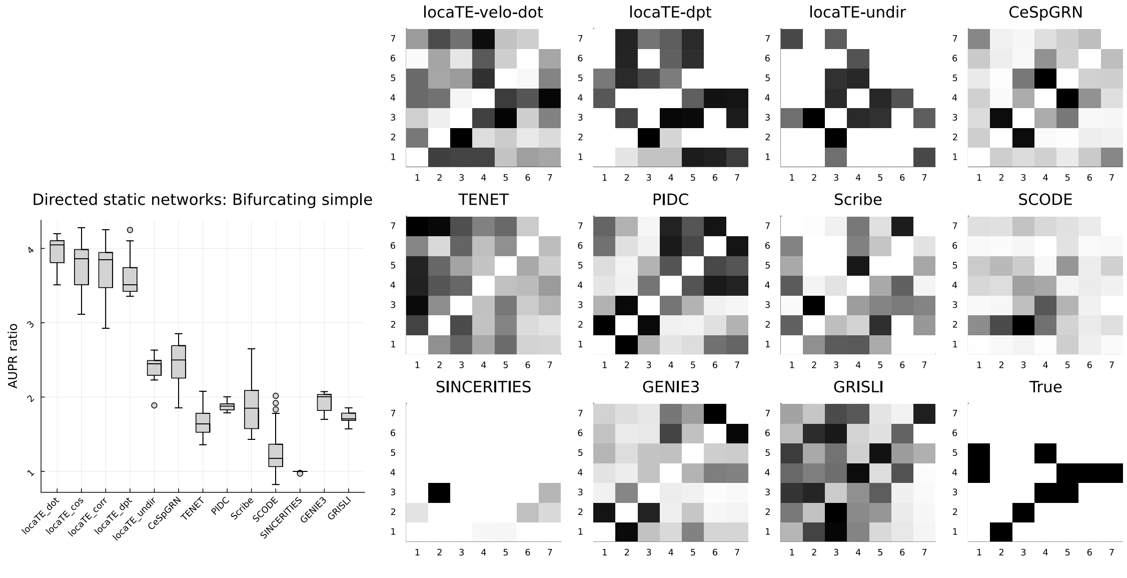


(b) Simple bifurcating network, 1000 cells.

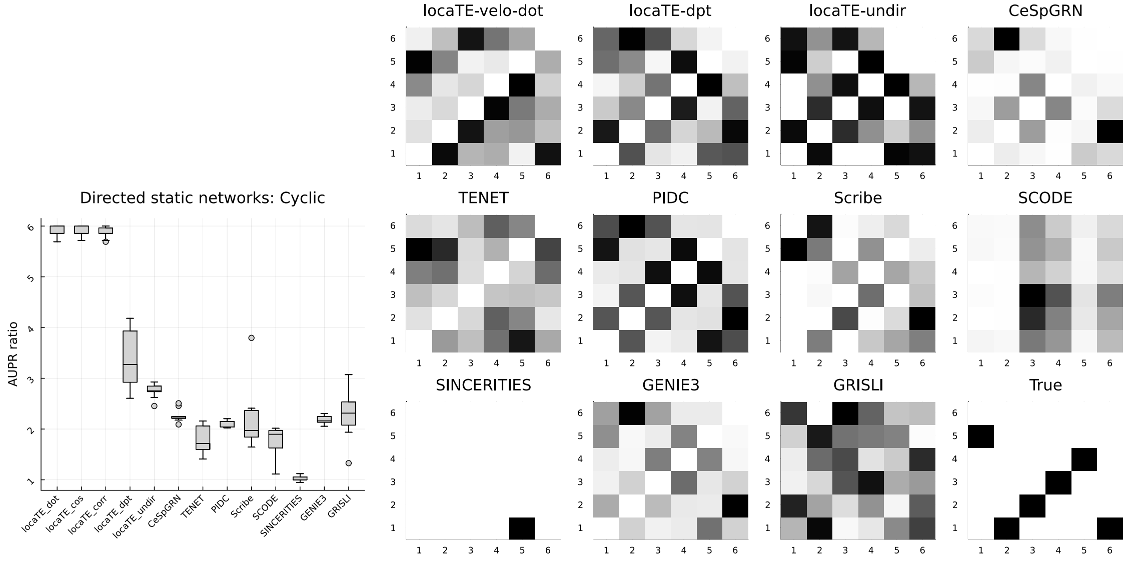


(c) Cycling network, 1000 cells.

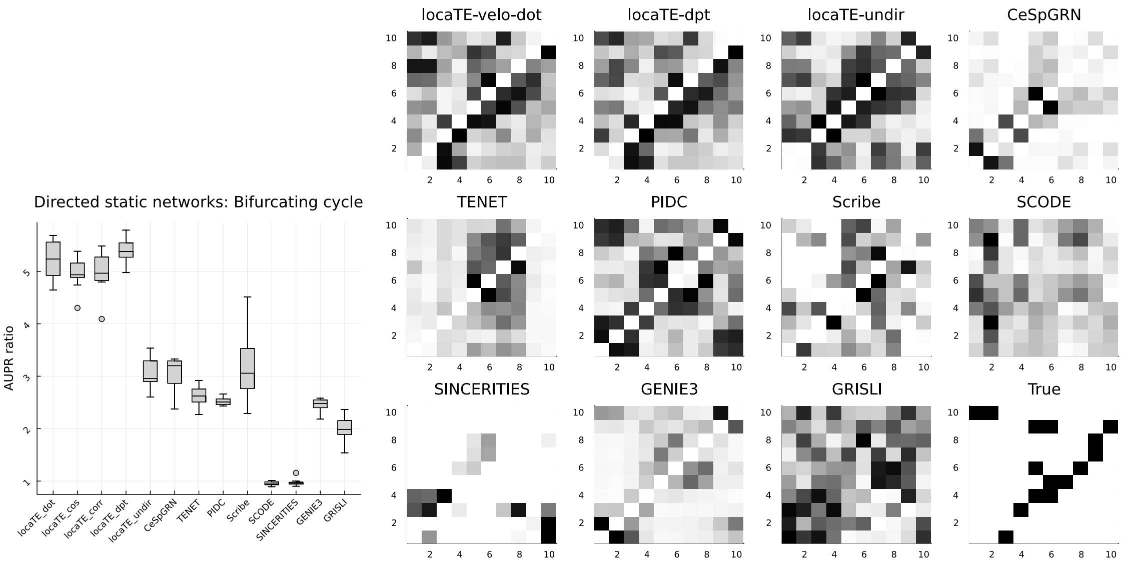


(a) Bifurcating cycling network, 1000 cells.

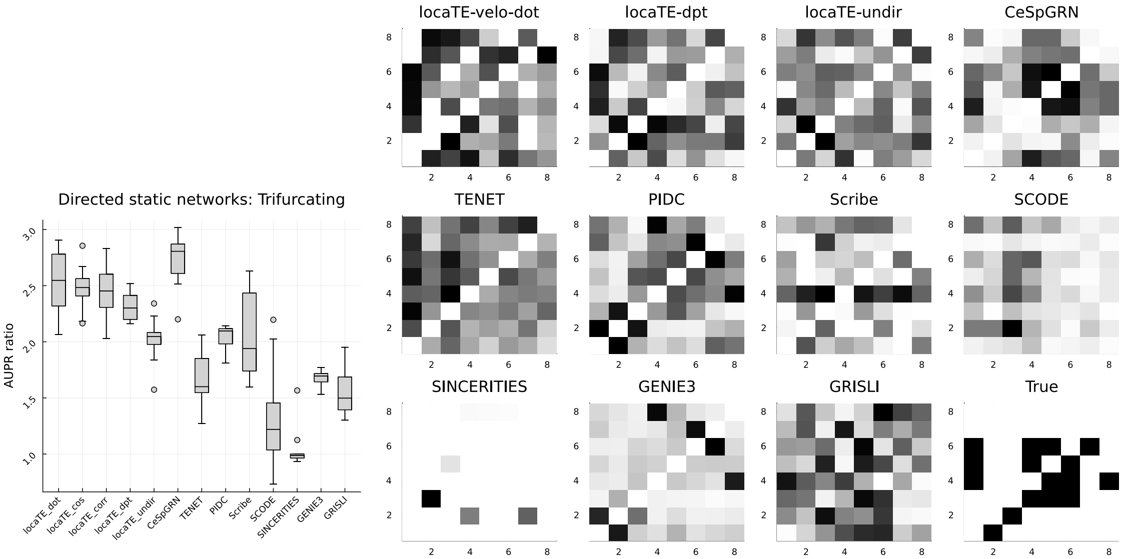


(b) Trifurcating network, 1000 cells.

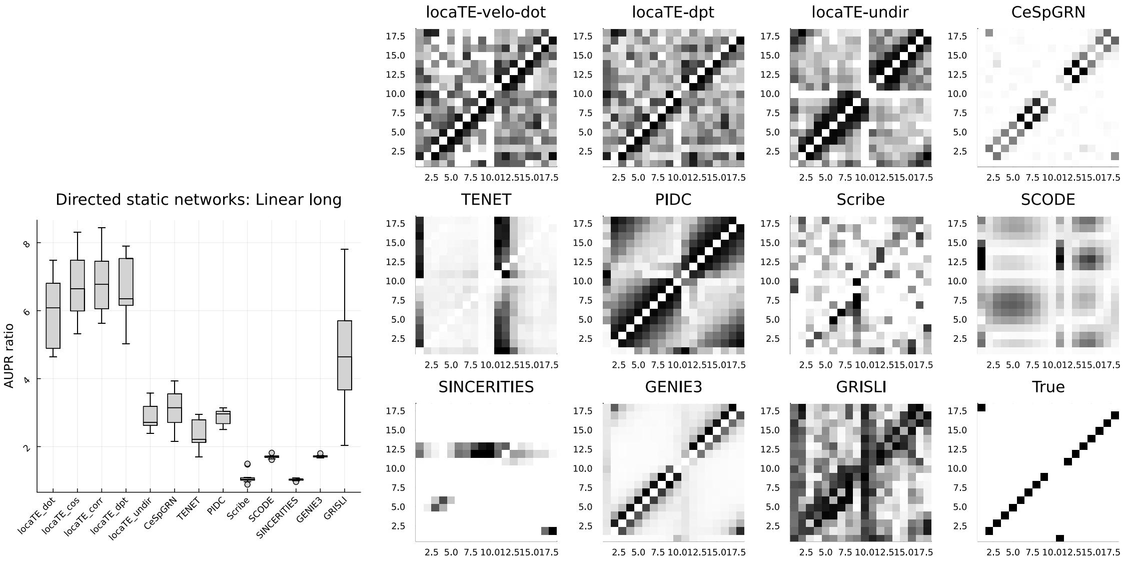




(c) Linear long network, 1000 cells.

#### S1.3 Effect of vector field noise and dropout

See Figure S7.

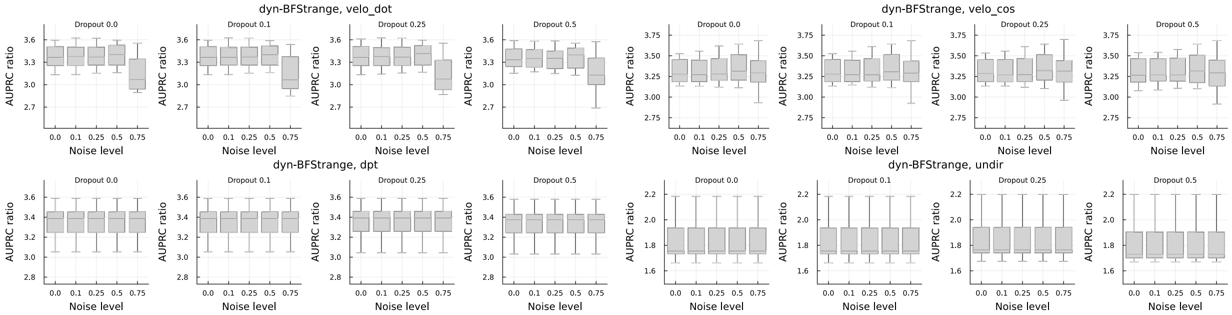


(a) Bifurcating network

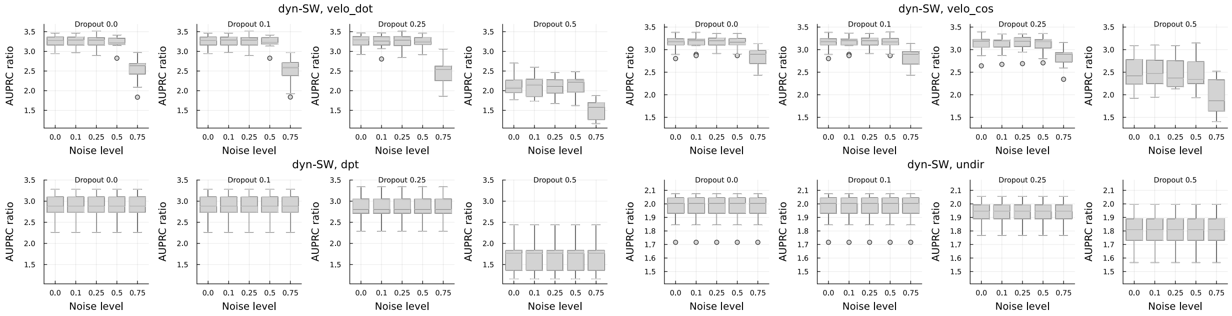


(b) Switch network

Supplementary Figure S7: Impact of vector field noise and dropout on performance of locaTE.

#### S1.4 Coarse-graining for large datasets

See Figure S8.

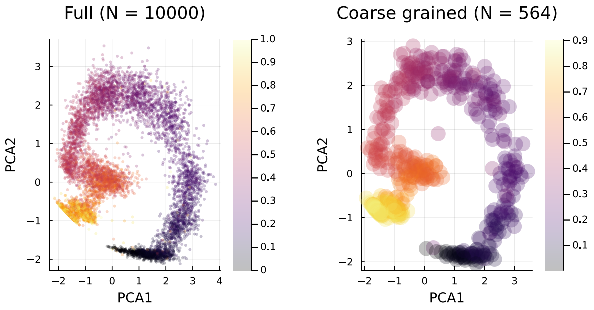


(a)

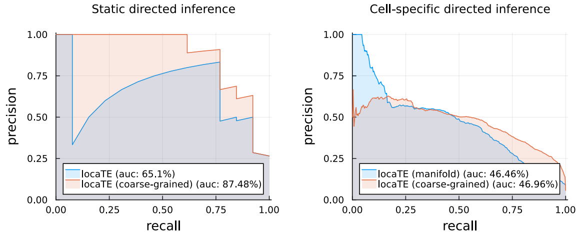


(b)

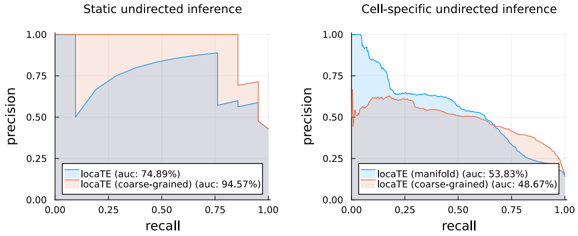


(c)

Supplementary Figure S8: (a) For the simulated switch-like network of Figure 3, from 10,000 simulated cell states the Leiden algorithm with resolution parameter 100 followed by PAGA [2] produces 564 coarse grained states, shown in PCA coordinates and coloured by simulation time. (b, c) Precision-recall curves for directed and undirected inference, using the full set of 10,000 cell states and $k$-NN graph, or using the 564 coarse grained states and PAGA state graph.

#### S1.5 mESC dataset

See Figures S9, S10.

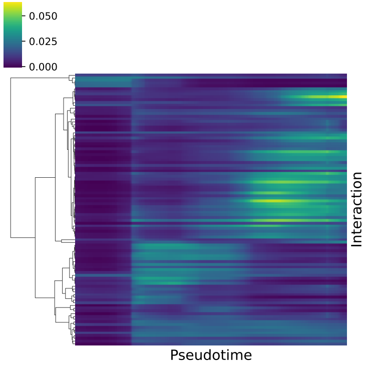


(a)

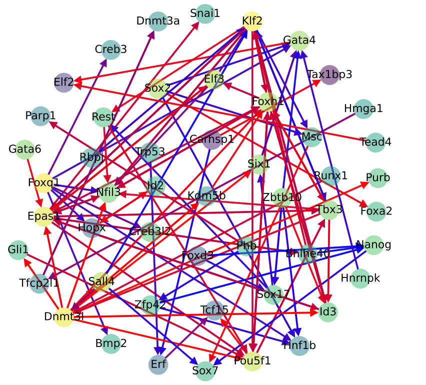


(b)

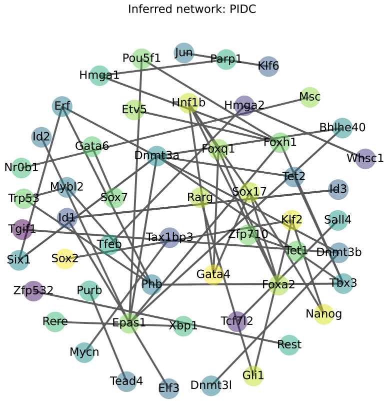


(c)

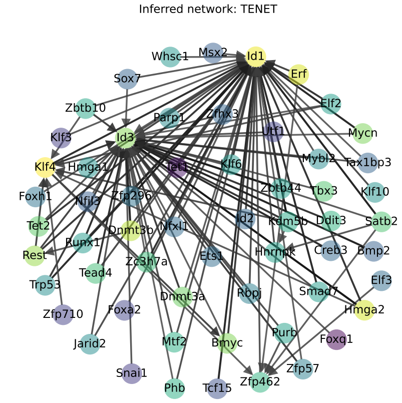


(d)

Supplementary Figure S9: Smoothed plot of top 1% of inferred interactions (from static network) (a) clustered and shown against pseudotime (b) coloured by the time of peak activity, where blue corresponds to earliest, and red corresponds to latest. (c, d) inferred networks (top 1% of inferred interactions) from PIDC [3] and TENET [4] respectively. For TENET, a history length of 3 was used.

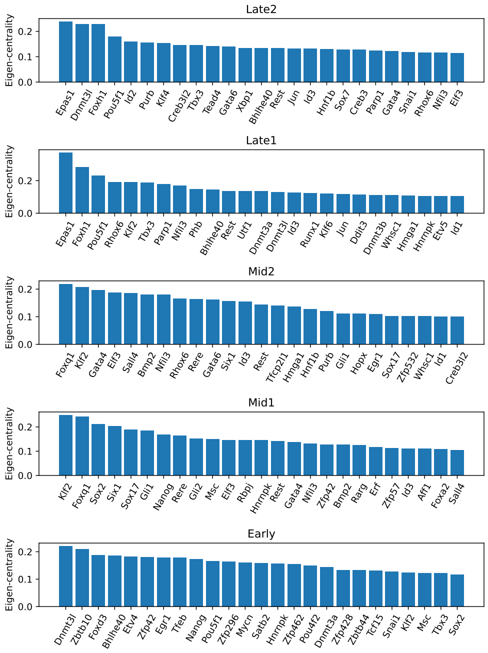


(a)

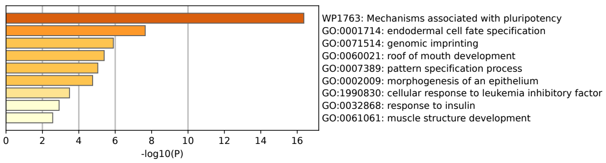


(b) Early

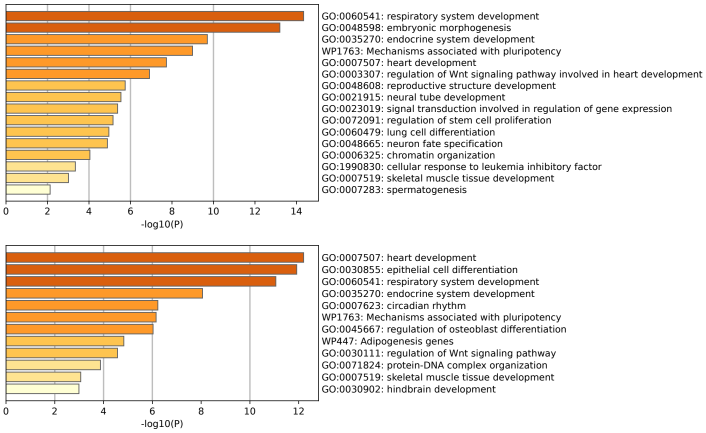


(c) Mid(1,2)

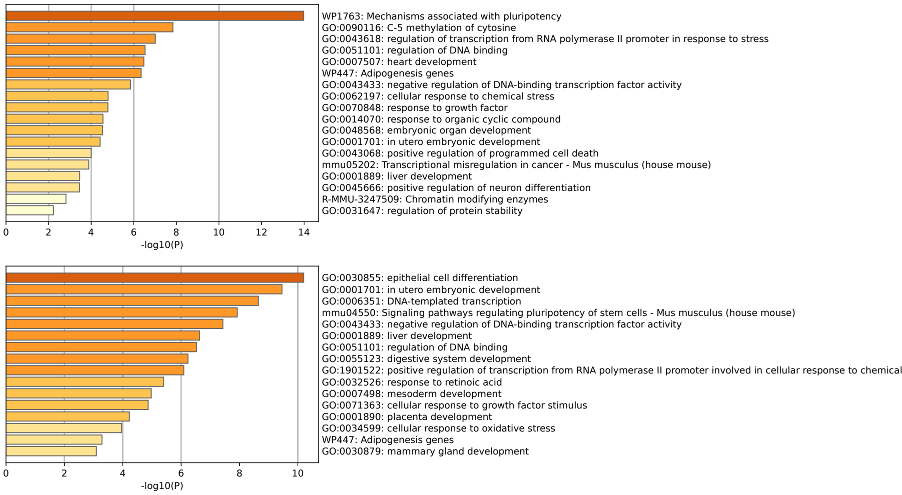


(d) Late(1,2)

Supplementary Figure S10: (a) Top 25 TFs for each regulatory module by outgoing eigenvector centrality (b-d) GO terms found by Metascape for top 25 TFs for each regulatory module.

#### S1.6 Pancreatic development dataset

See Figures S11, S12, S13, S14.

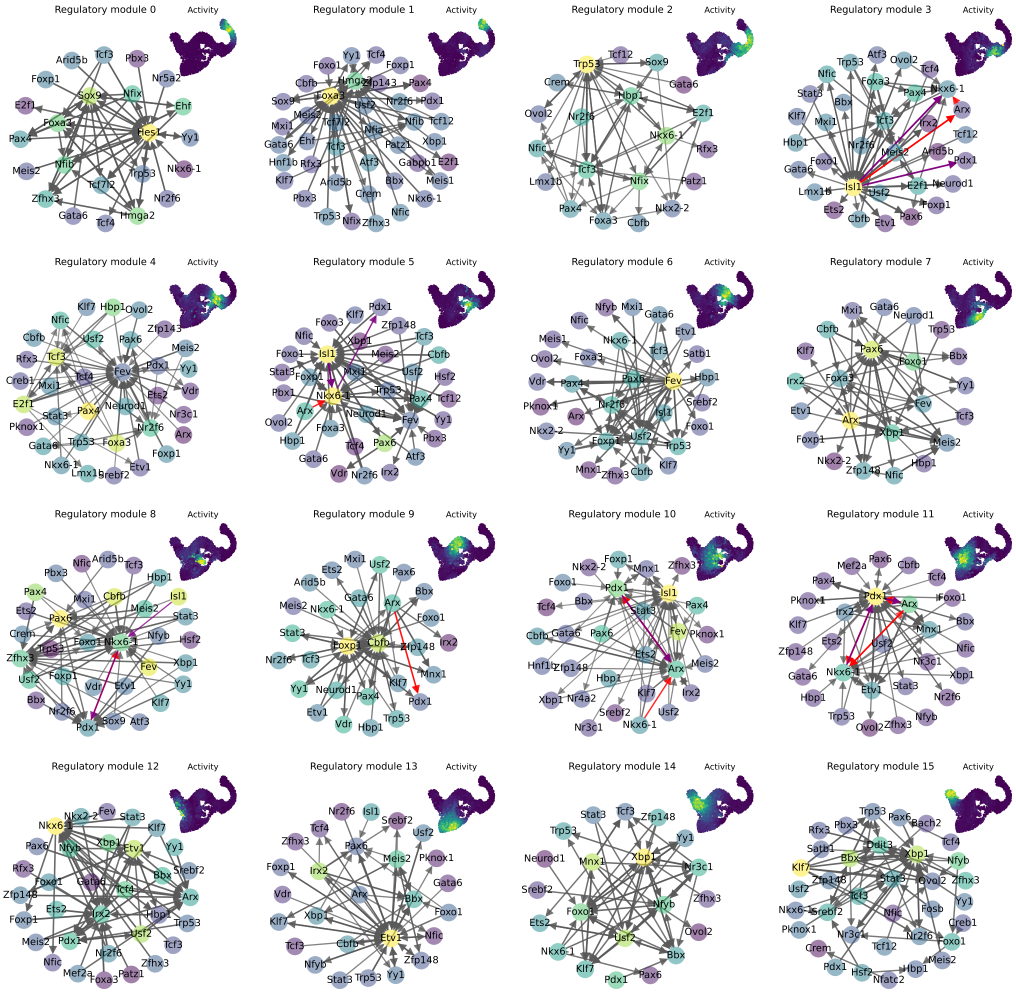


Supplementary Figure S11: Pancreatic development regulatory modules (top 1% of interactions) and module activities (inset) on UMAP coordinates.

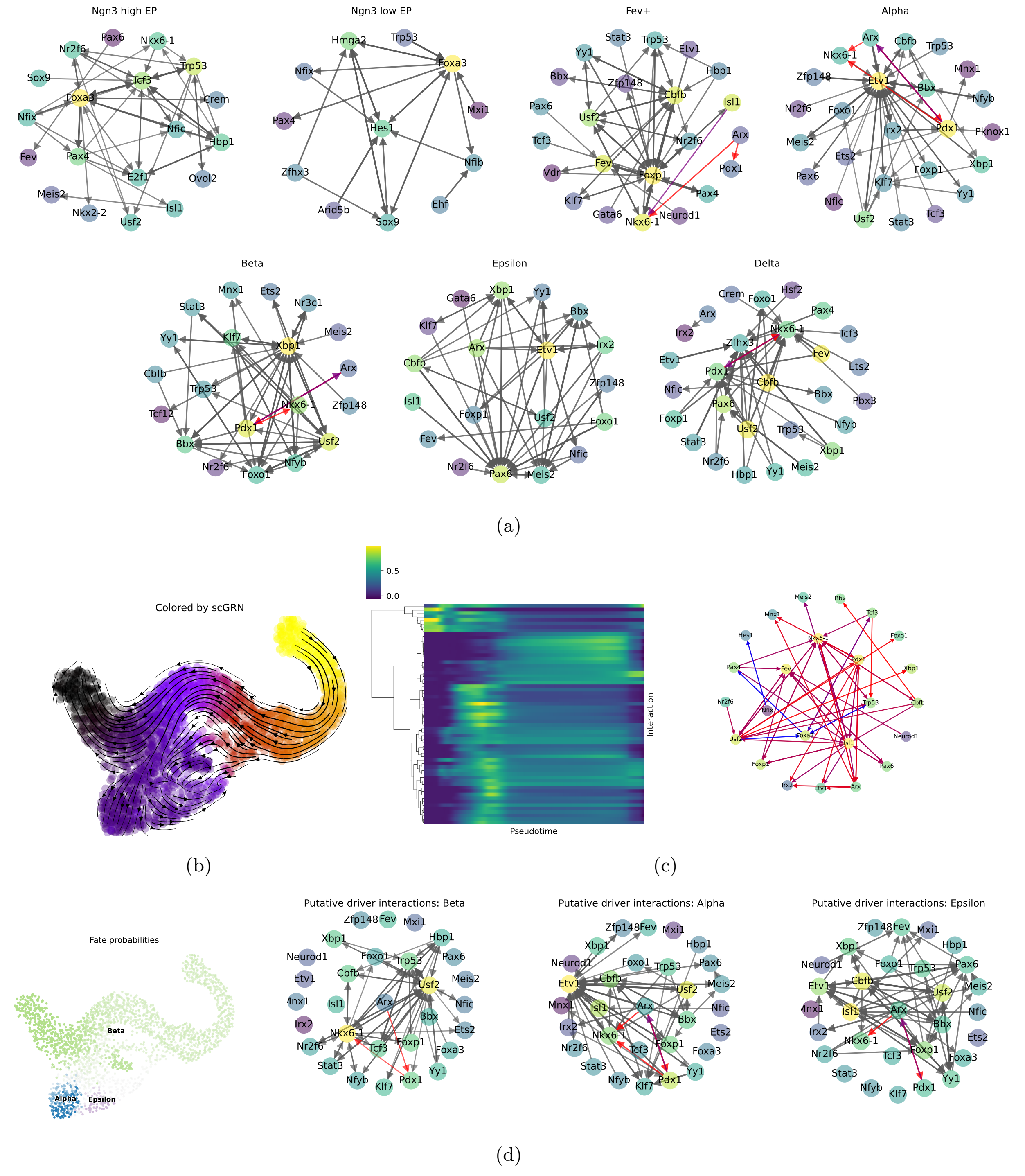


Supplementary Figure S12: (a) Per-cluster networks (top 1% of interactions) (b) First diffusion component of cell-specific GRNs found by locaTE overlaid on UMAP coordinates (c) Smoothed plot of interactions (top 1% by confidence) clustered and shown against pseudotime, and corresponding static network with edges coloured by time of peak activity (d) Putative driver networks (top 1% by confidence) for alpha, beta and epsilon fates.

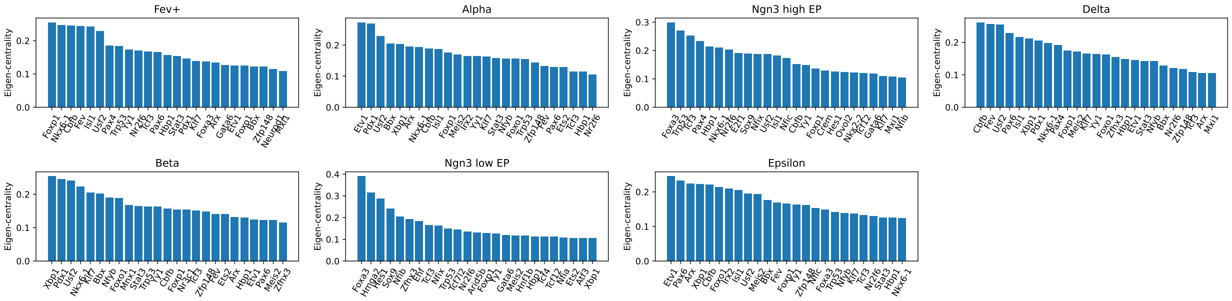


(a)

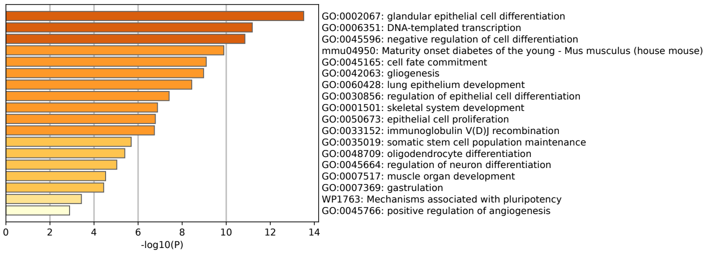


(b) Ngn3- EP cluster

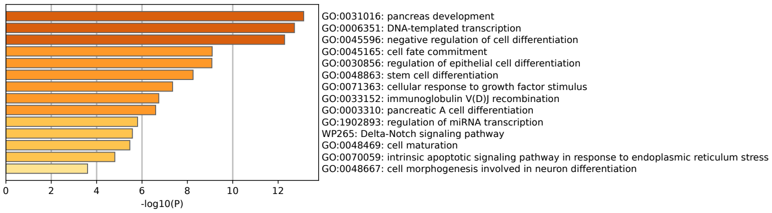


(c) Ngn3+ EP cluster

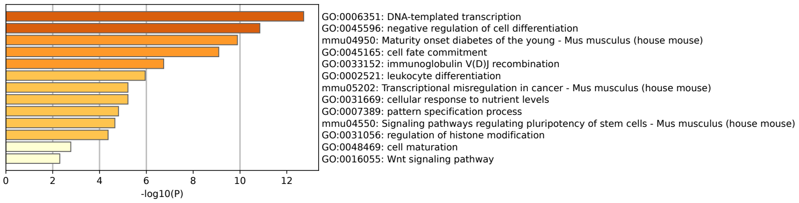


(d) Beta cluster

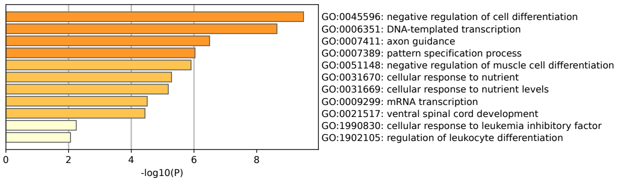


(e) Alpha cluster

Supplementary Figure S13: **(a)** Top 25 TFs for each stage by outgoing eigenvector centrality **(b-e)** GO terms found by Metascape for top 25 TFs from Ngn3- EP, Ngn3+ EP, Beta, Alpha clusters.

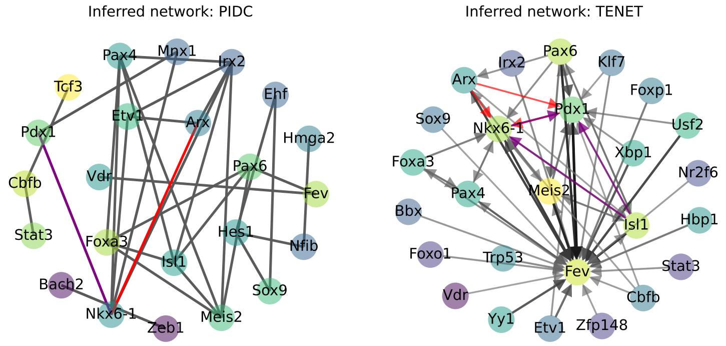


Supplementary Figure S14: Inferred networks (top 1% of inferred interactions) from PIDC [3] and TENET [4] respectively. For TENET, a history length of 3 was used.

#### S1.7 Haematopoiesis (Lassauzaie et al.) dataset

See Figures S15, S16, S17, S18.

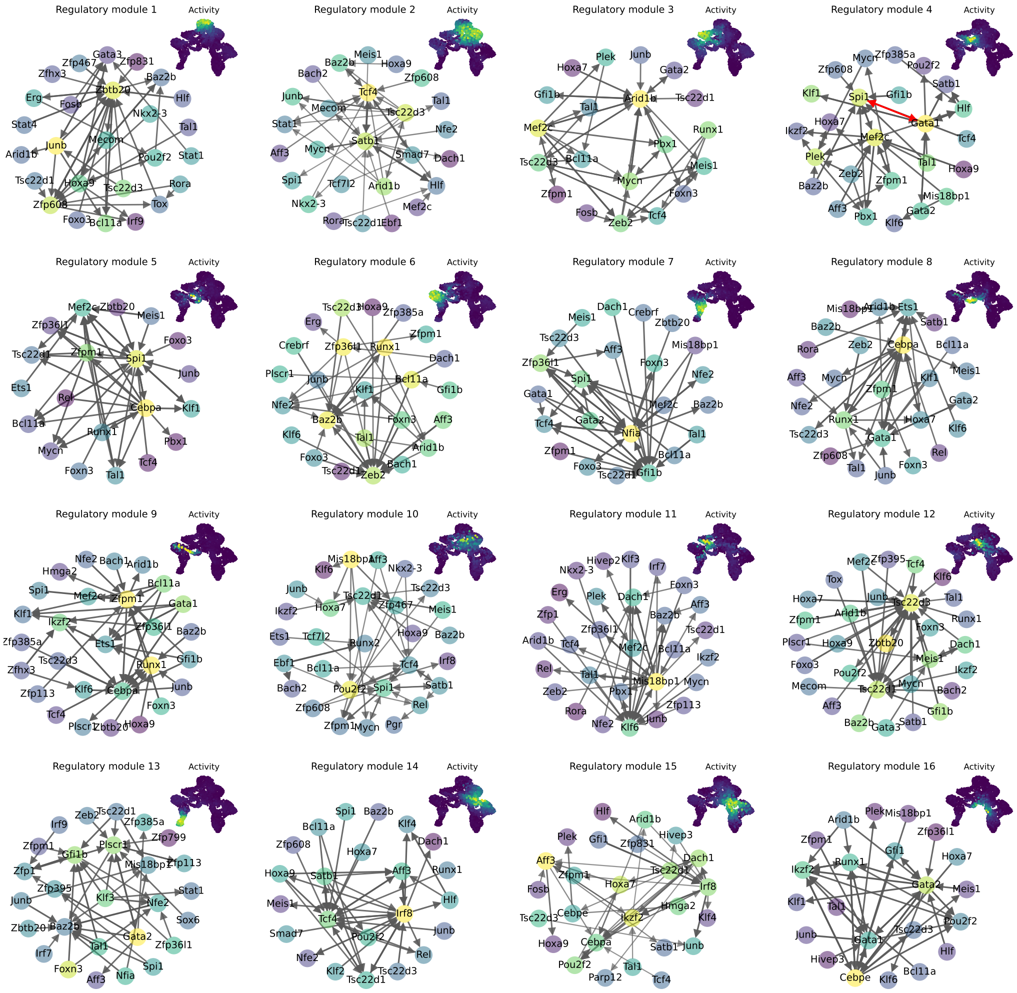


(a)

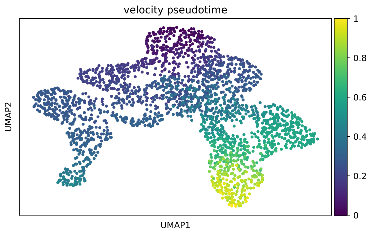


(b)

Supplementary Figure S15: (a) Haematopoiesis regulatory modules (top 0.5% of edges by confidence) and module activities (inset) on UMAP coordinates. (b) Velocity pseudotime shown on UMAP coordinates highlights insufficiency of a 1D ordering.

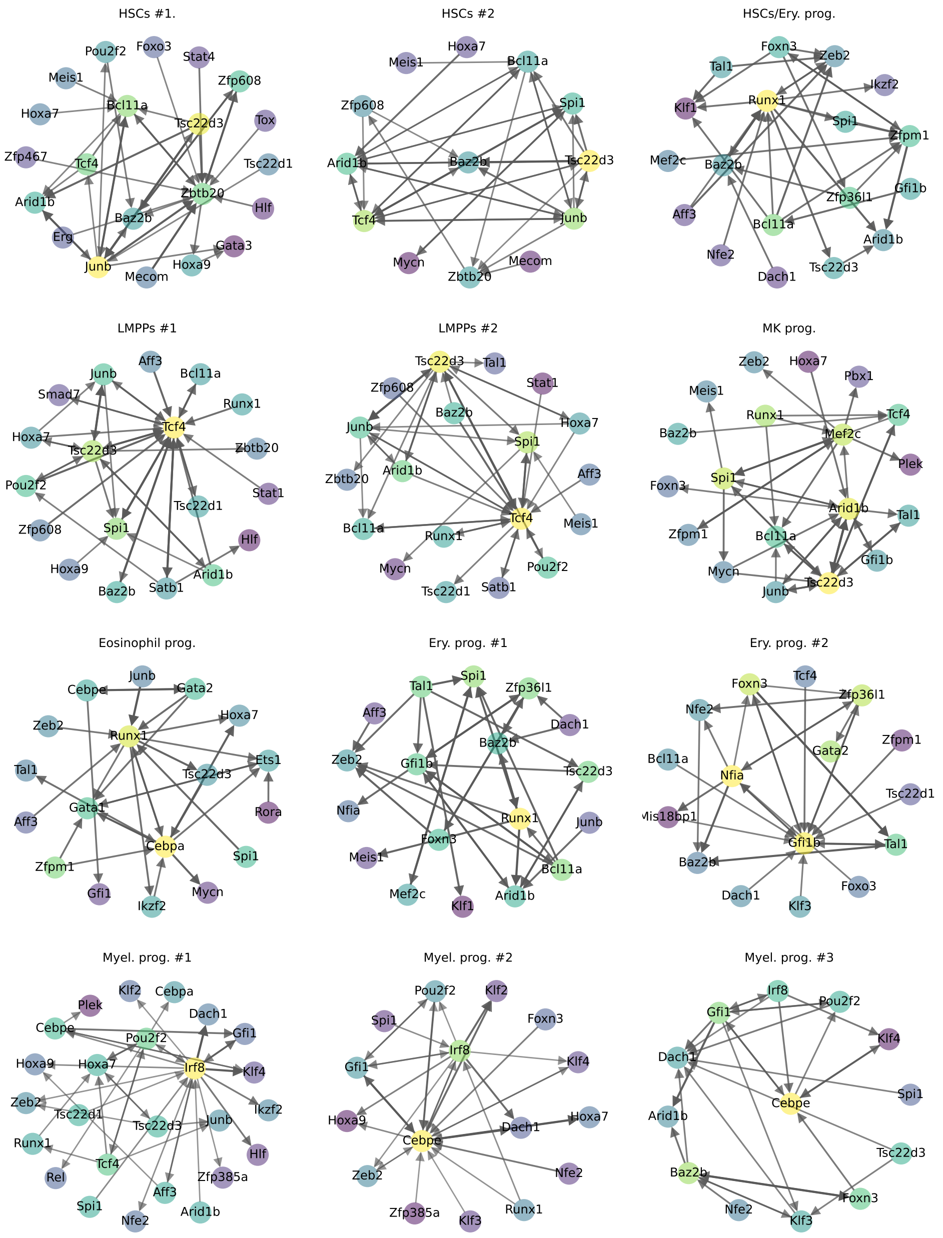


Supplementary Figure S16: Cluster-averaged inferred networks for annotated clusters (top 0.5% of edges by confidence).

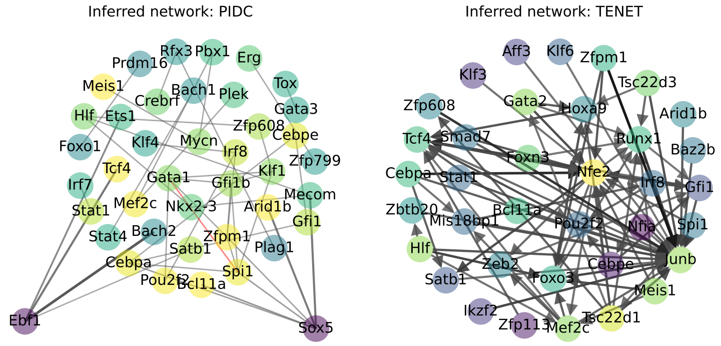


Supplementary Figure S17: Inferred networks (top 1% of inferred interactions) from PIDC [3] and TENET [4] respectively. For TENET, a history length of 3 was used.

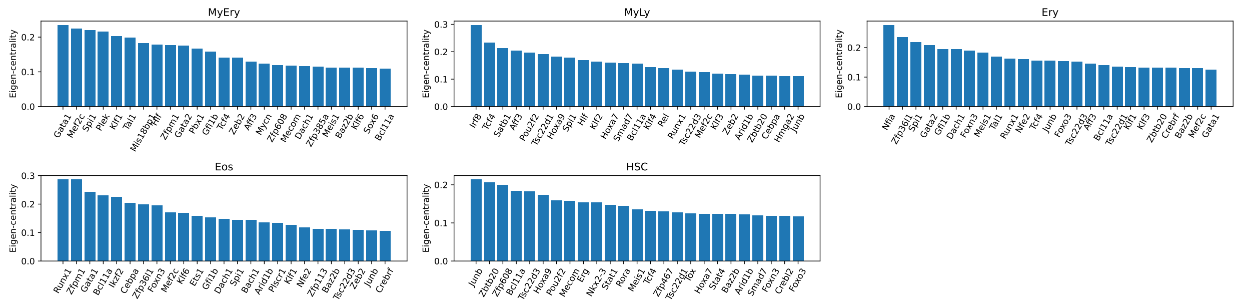


(a)

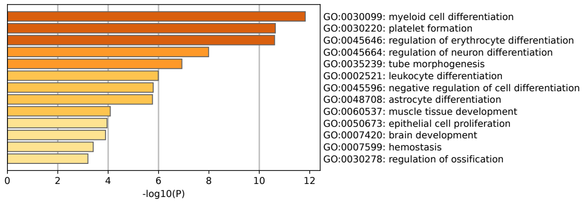


(b) Myeloid-Erythroid module

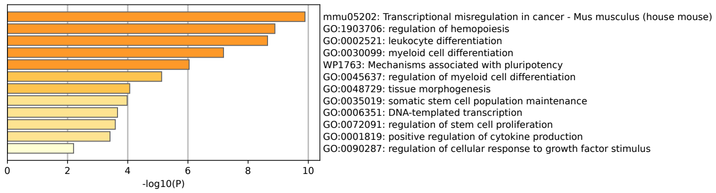


(c) Myeloid-Lymphoid module

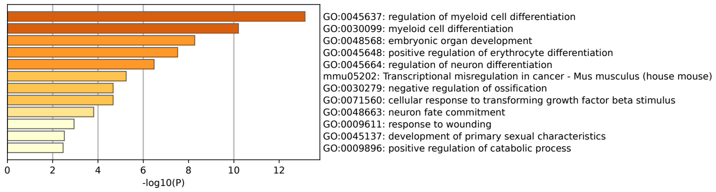


(d) Erythroid module

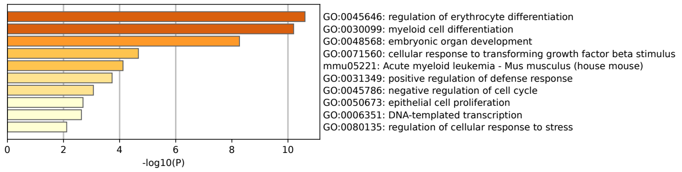


(e) Eosinophil module

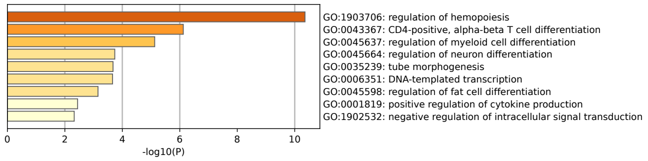


(f) HSC module

Supplementary Figure S18: (a) Top 25 TFs for each regulatory module ranked by out-edge eigenvector centrality. (b) Corresponding GO terms found by Metascape.

#### S1.8 Haematopoiesis (Tusi et al.) dataset

See Figure S19.

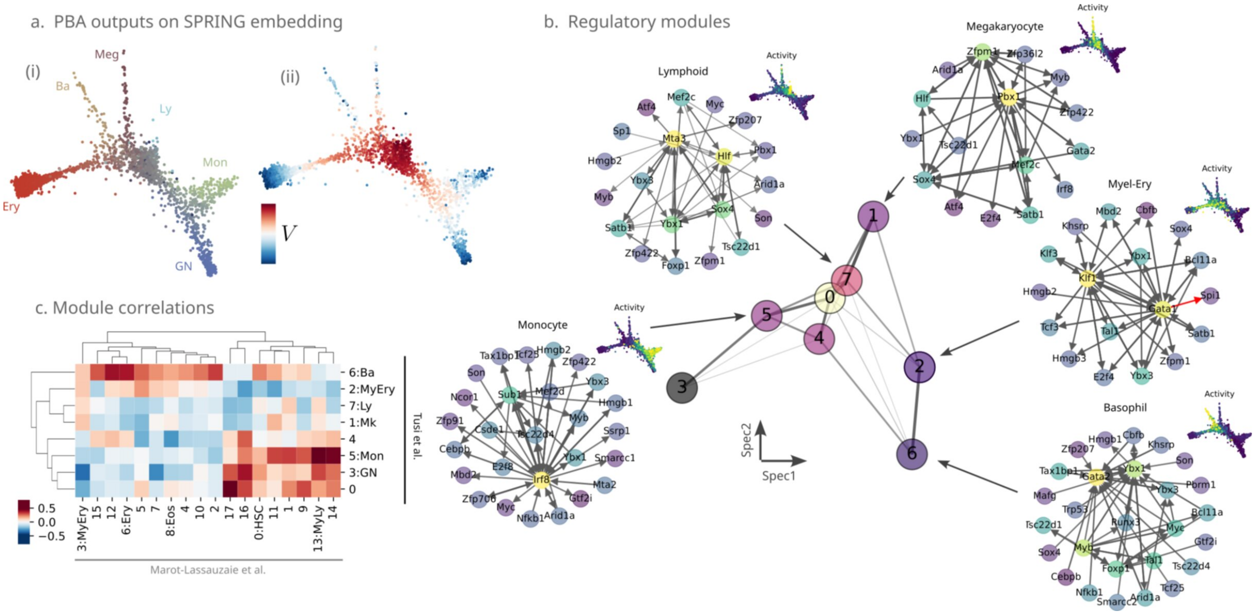


Supplementary Figure S19: **Tusi et al. haematopoiesis dataset (a)** Single cell SPRING embeddings from the dataset of Tusi et al. [5, 6] shown by (i) fate annotation and (ii) inferred population balance analysis (PBA) potential, $V(x)$. **(b)** Graph layout of regulatory modules found by locaTE-NMF colored by proximity from a progenitor module, shown alongside representative networks of selected modules (top 0.5% of all filtered edges), corresponding to Monocyte, Lymphoid, Megakaryocyte, Myeloid-Erythroid, and Basophil. **(c)** Similarities between regulatory modules found for each dataset, measured in terms of Spearman rank correlation of overlapping gene eigencentralities.

#### S1.9 Cell-cycle dataset

See Figure S21, S20.

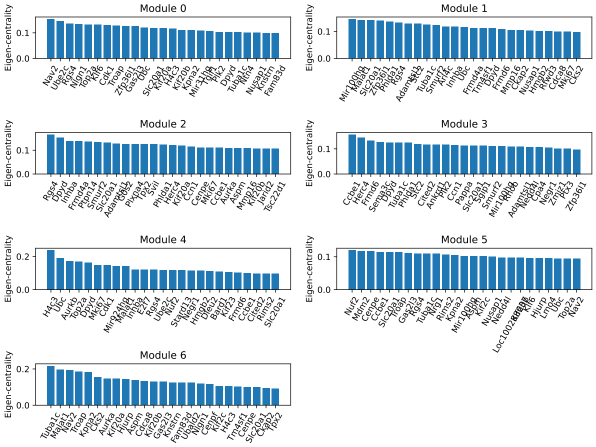


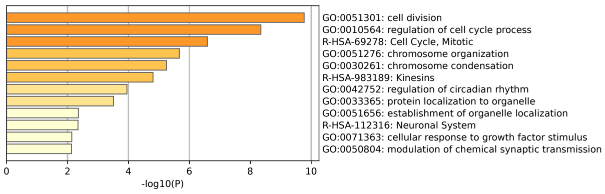


(a) Module 1

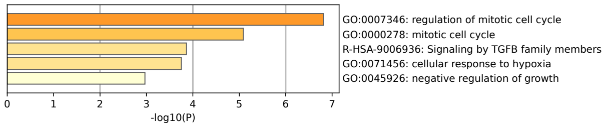


(b) Module 2

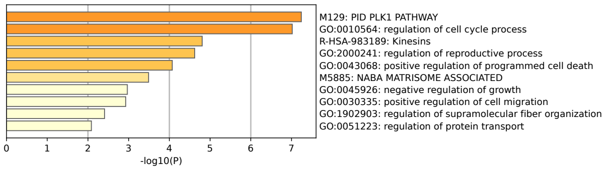


(c) Module 3

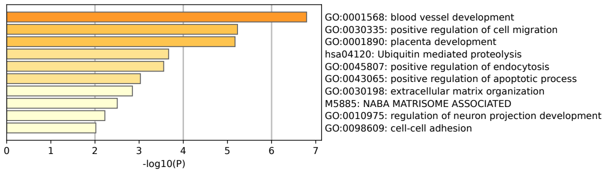


(d) Module 4

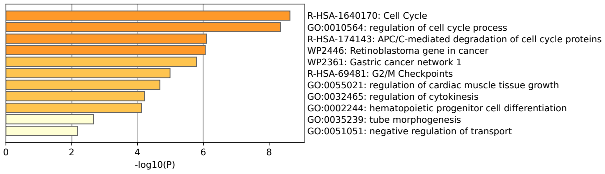


(e) Module 5

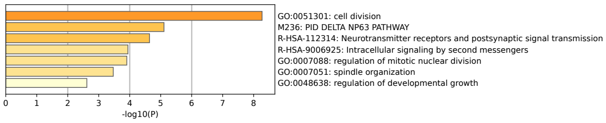


(f) Module 6

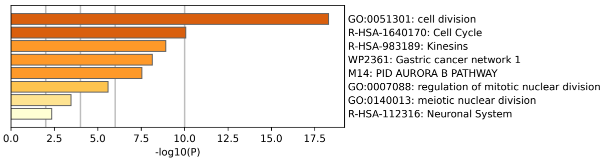


(g) Module 7

Supplementary Figure S20: (a) Top 25 genes for each interaction module by outgoing eigenvector centrality. (b) GO terms found by Metascape for top 25 genes for each interaction module.

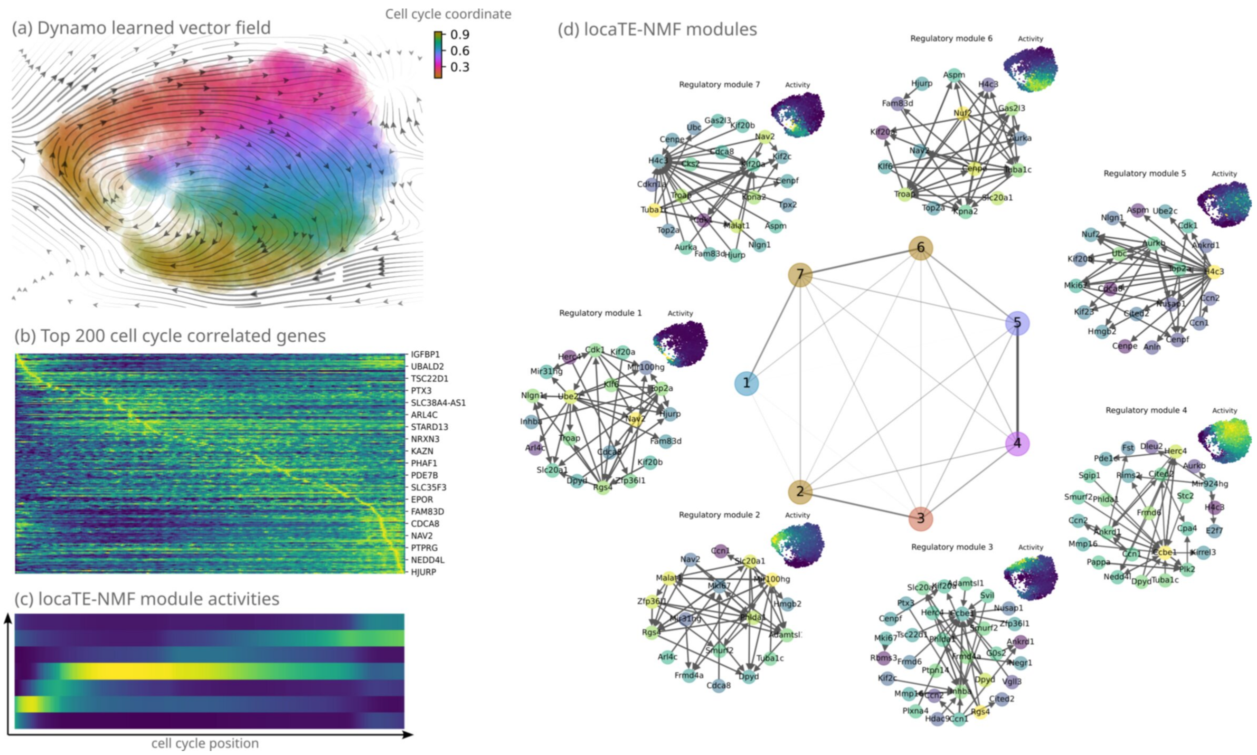


Supplementary Figure S21: (a) Cell-cycle dataset from [7, 8] shown in UMAP coordinates and coloured according to cell cycle progression, with vector field learned using Dynamo [8] overlaid. (b) Expression profiles of top 200 cell-cycle correlated genes against cell cycle progression. (c, d) Interaction modules and module activities learned by locaTE-NMF with $\alpha = 0.75$.

### References


[1] Aditya Pratapa, Amogh P Jalihal, Jeffrey N Law, Aditya Bharadwaj, and TM Murali. Benchmarking algorithms for gene regulatory network inference from single-cell transcriptomic data. *Nature methods*, 17(2):147–154, 2020.

[2] F Alexander Wolf, Fiona K Hamey, Mireya Plass, Jordi Solana, Joakim S Dahlin, Berthold Göttgens, Nikolaus Rajewsky, Lukas Simon, and Fabian J Theis. Paga: graph abstraction reconciles clustering with trajectory inference through a topology preserving map of single cells. *Genome biology*, 20:1–9, 2019.

[3] Thalia E Chan, Michael PH Stumpf, and Ann C Babtie. Gene regulatory network inference from single-cell data using multivariate information measures. *Cell systems*, 5(3):251–267, 2017.

[4] Junil Kim, Simon T. Jakobsen, Kedar N Natarajan, and Kyoung-Jae Won. Tenet: gene network reconstruction using transfer entropy reveals key regulatory factors from single cell transcriptomic data. *Nucleic acids research*, 49(1):e1–e1, 2021.

[5] Betsabeh Khoramian Tusi, Samuel L Wolock, Caleb Weinreb, Yung Hwang, Daniel Hidalgo, Rapolas Zilionis, Ari Waisman, Jun R Huh, Allon M Klein, and Merav Socolovsky. Population snapshots predict early haematopoietic and erythroid hierarchies. *Nature*, 555(7694):54–60, 2018.

[6] Caleb Weinreb, Samuel Wolock, Betsabeh K Tusi, Merav Socolovsky, and Allon M Klein. Fundamental limits on dynamic inference from single-cell snapshots. *Proceedings of the National Academy of Sciences*, 115(10):E2467–E2476, 2018.

[7] Nico Battich, Joep Beumer, Buys de Barbanson, Lenno Krenning, Chloé S Baron, Marvin E Tanenbaum, Hans Clevers, and Alexander van Oudenaarden. Sequencing metabolically labeled transcripts in single cells reveals mrna turnover strategies. *Science*, 367(6482):1151–1156, 2020.

[8] Xiaojie Qiu, Yan Zhang, Jorge D Martin-Rufino, Chen Weng, Shayan Hosseinzadeh, Dian Yang, Angela N Pogson, Marco Y Hein, Kyung Hoi Joseph Min, Li Wang, et al. Mapping transcriptomic vector fields of single cells. *Cell*, 185(4):690–711, 2022.
